# Self-Organizing Global Gene Expression Regulated Through Criticality: Mechanism of the Cell-Fate Change

**DOI:** 10.1101/066498

**Authors:** Masa Tsuchiya, Alessandro Giuliani, Midori Hashimoto, Jekaterina Erenpreisa, Kenichi Yoshikawa

## Abstract

**Background:** A fundamental issue in bioscience is to understand the mechanism that underlies the dynamic control of genome-wide expression through the complex temporal-spatial self-organization of the genome to regulate the change in cell fate. We address this issue by elucidating a physically motivated mechanism of self-organization.

**Principal Findings:** Building upon transcriptome experimental data for seven distinct cell fates, including early embryonic development, we demonstrate that self-organized criticality (SOC) plays an essential role in the dynamic control of global gene expression regulation at both the population and single-cell levels. The novel findings are as follows:

**i) Mechanism of cell-fate changes**: A sandpile-type critical transition self-organizes overall expression into a few transcription response domains (critical states). A cell-fate change occurs by means of a dissipative pulse-like global perturbation in self-organization through the erasure of initial-state critical behaviors (criticality). Most notably, the reprogramming of early embryo cells destroys the zygote SOC control to initiate self-organization in the new embryonal genome, which passes through a stochastic overall expression pattern.

**ii) Mechanism of perturbation of SOC controls**: Global perturbations in self-organization involve the temporal regulation of critical states. Quantitative evaluation of this perturbation in terminal cell fates reveals that dynamic interactions between critical states determine the critical-state coherent regulation. The occurrence of a temporal change in criticality perturbs this between-states interaction, which directly affects the entire genomic system. Surprisingly, a sub-critical state, corresponding to an ensemble of genes that shows only marginal changes in expression and consequently are considered to be devoid of any interest, plays an essential role in generating a global perturbation in self-organization directed toward the cell-fate change.

**Conclusion and Significance:** ‘Whole-genome’ regulation of gene expression through self-regulatory SOC control complements gene-by-gene fine tuning and represents a still largely unexplored non-equilibrium statistical mechanism that is responsible for the massive reprogramming of genome expression.

**A List of Abbreviations**
SOC: self-organized criticality
CP: critical point
CM: center of mass
CES: coherent expression state nrmsf; normalized root mean square fluctuation
CSB: coherent-stochastic behavior
ABS: autonomous bistable switch

## Introduction

In mature mammalian stem cells, the cell fate/state can be reprogrammed to provoke a shift between two stable (and very different) gene expression profiles involving tens of thousands of genes by means of a few reprogramming stimuli [1-5].

The coordinated control of a large number of genes must overcome several difficulties, such as the substantial instability of genetic products due to the stochastic noise stemming from the low copy number of specific gene mRNAs per cell and the lack of a sufficient number of molecules to reach a thermodynamic limit [6,7]. Due to the complexity of the interaction between molecular effectors and changes in the structure of chromatin, it has been a challenging issue to understand how globally coordinated control can determine the cell fate/state from a genomic point of view. In this respect, it is important to gain a comprehensive understanding of dynamic control mechanisms that could help us to obtain a quantitative appreciation of the still largely qualitative notion of the epigenetic landscape [8]. The existence of global gene regulation implies that the driving force of genomic expression acts through only a small number of control parameters that underpin highly complex molecular genetics reaction mechanisms. The hypothesis that a reliable model of a complex system can be obtained through the use of few relevant parameters was aptly addressed by Transtrum *et al*. [9] in terms of ‘sloppiness’:

> *“First, in spite of the large number of parameters, complex biological systems typically exhibit simple behavior that requires only a few parameters to describe, analogous to how the diffusion equation can describe microscopically diverse processes. Attempting to accurately infer all of the parameters in a complex biological model is analogous to learning all of the mechanical and electrical properties of water molecules in order to accurately predict a diffusion constant. It would involve considerable effort (measuring all the microscopic parameters accurately), while the diffusion constant can be easily measured using collective experiments and used to determine the result of any other collective experiment. Second, in many biological systems, there is considerable uncertainty about the microscopic structure of the system. Sloppiness suggests that an effective model that is microscopically inaccurate may still be insightful and predictive in spite of getting many specific details wrong.”*

Here we explore the potential of coarse-grain statistical metrics regarding the expression levels of gene ensembles [10,11] to sketch a model of biological regulation within the framework of ‘statistical mechanics’. This approach may clarify how a small number of hidden control parameters can provoke a global change in the expression profile involving thousands of genes.

Scientists working with microarray technology (cell population) are familiar with the strict profile-invariance of independent samples relative to the same type of tissue. The expression vectors for two independent samples of the same tissue, which consist of about 20,000 ORFs, show a near-unity Pearson correlation, which points to a very strict global integration of cell populations in terms of gene expression.

The presence of global tissue-level control is evident when we compare two samples from different tissues (Figure 1). A near-unity correlation is observed between the profiles of independent samples of the same kind of tissue (Figure 1A). In contrast, when we consider samples from different tissues, this near-unity correlation breaks down (Figure 1B). This simple plot suggests that global self-organization supports the phenotype that corresponds to different cell types and involves the whole expression profile in cell population dynamics. It is very interesting to look at the results depicted in Figure 1 from the point of view of criticality-induced complexity matching [12,13] in the field of complex networks.

**Figure 1:**
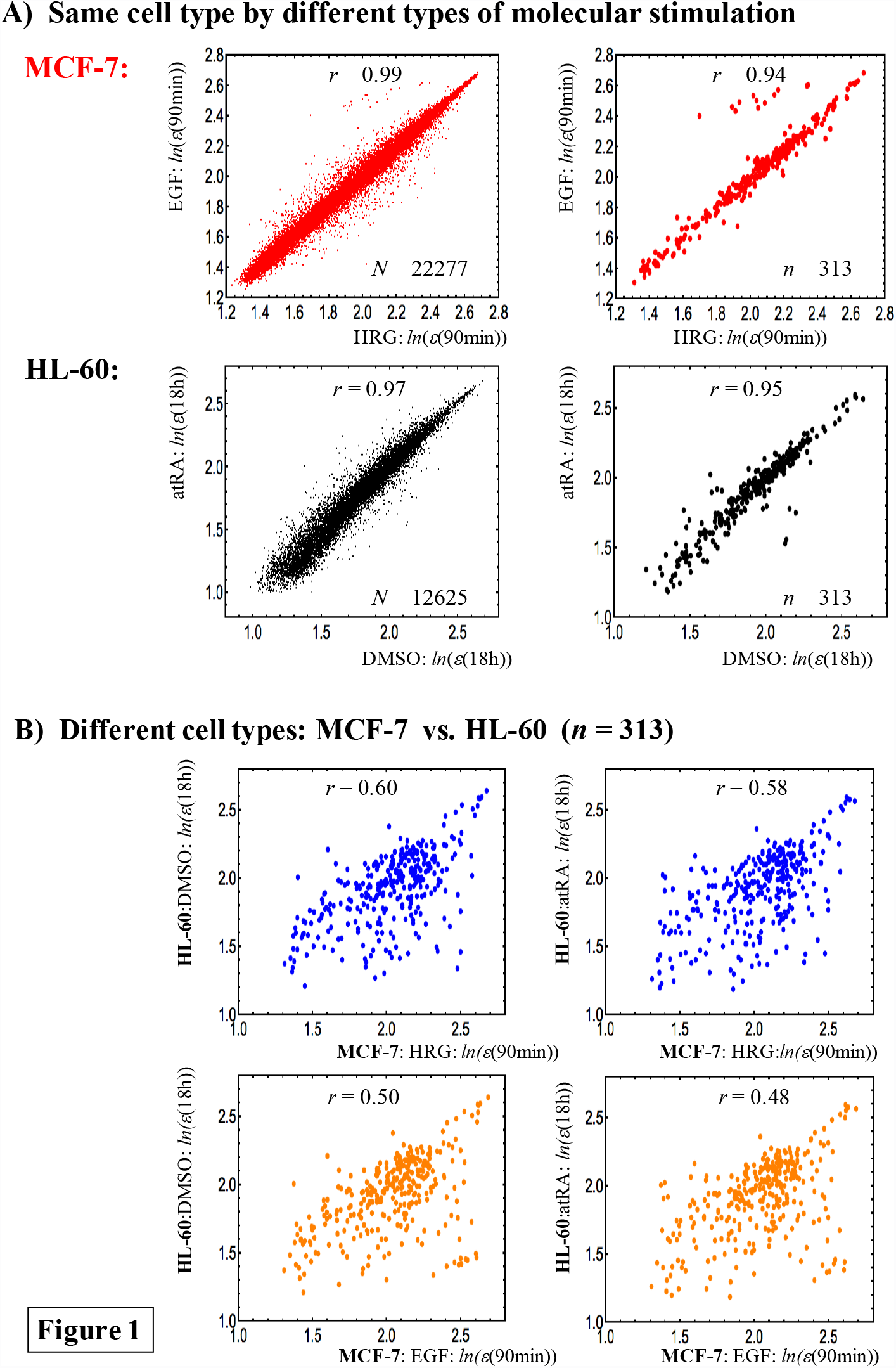
Correlation of gene expression profiles: *A) **The same cell types**: right panels: a near-unity Pearson correlation, r, in whole gene expression (N: total number of mRNAs) within the same cell type is shown for different types of molecular stimulation (first row: HRG- vs. EGF-stimulated MCF-7 cells; second row: DMSO- vs. atRA-stimulated HL-60 cells). Left panels: 313 (n) gene expressions, which have a common probe ID among four transcriptome microarray expression data sets (see **Methods**) also show a near-unity Pearson correlation within the same cell type.* *B) **Different cell types**: The near-unity Pearson correlation between independent samples of the same cell type breaks down when the gene expression profiles come from different (HL-60 and MCF-7) cell types.* e*(t) represents the ensemble of expression at time point t (N: the whole set; n: an ensemble set) and ln(*e*(t)) represents its natural logarithm, where the natural log of an individual expression value is taken. Plots show t = 90min for MCF-7-stimulated cells and t = 18h for HL-60-stimulated cells.*

Furthermore, Figure 2 suggests that the observed profile invariance is an emergent property that strictly depends on the range of gene expression being considered. If we change the box size from genes with very similar expression levels (low between-gene expression variance) to the whole set, gene expression shifts from a stochastic to a genome-wide attractor profile (which causes a near-unity Pearson correlation). The development of this correlation demonstrates the presence of a transition that follows a tangent hyperbolic function (inset in Figure 2). This implies that, while myriad transcriptional regulation control circuits are active at the same time at a local level (which gives a stochastic distribution; refer to **section IV**), at the global level of genome expression, very efficient tissue-level self-organization accompanied by “higher-order cooperativity” [14] emerges. Such self-organization involves the parallel regulation of more than 20,000 of different and functionally heterogeneous genes. This in turn suggests that the ordination of gene ensembles (a coarse-grained approach [15-18]) according to their expression level could be useful candidate for exploring genome-wide regulation.

**Figure 2:**
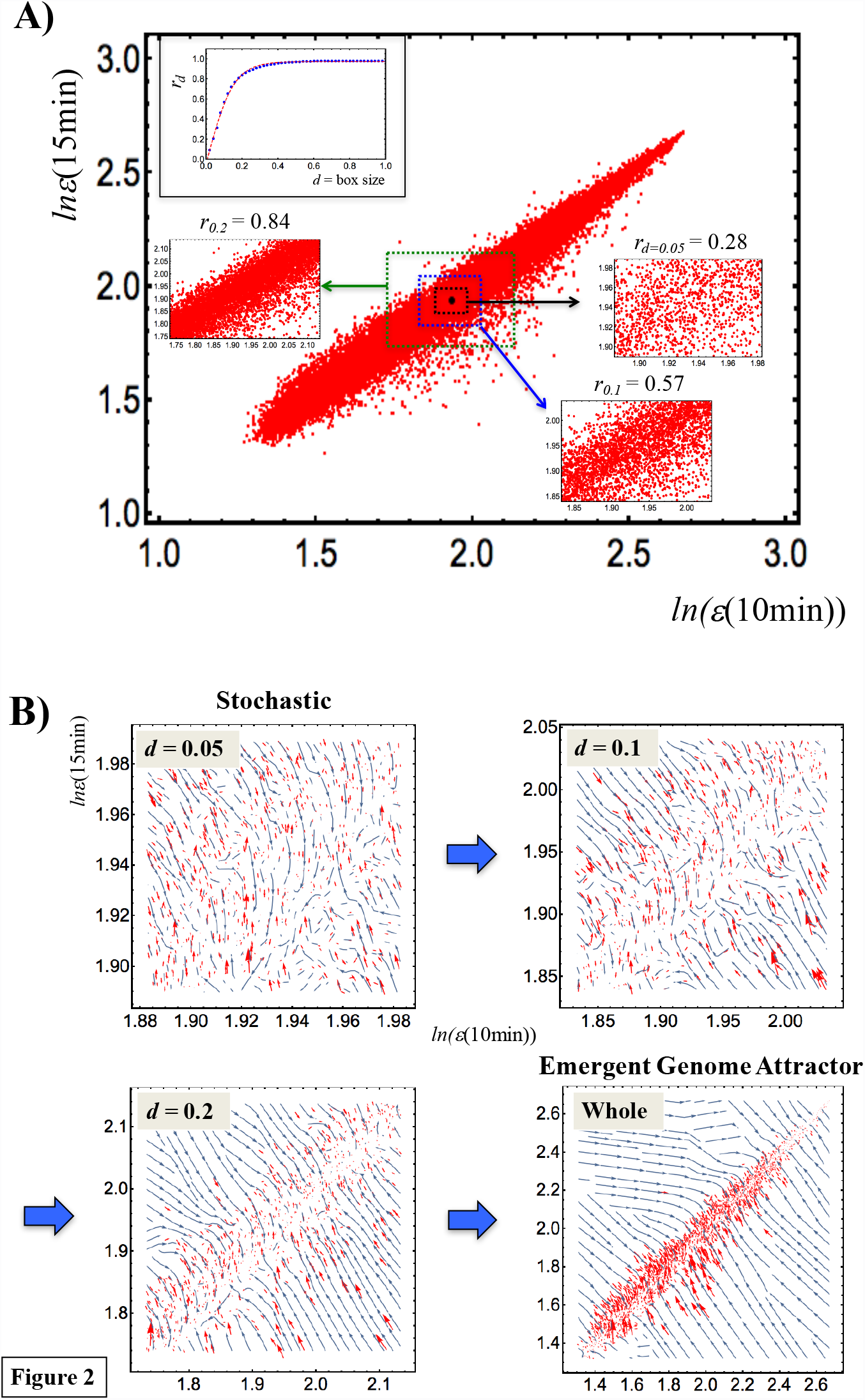
Transition of gene expression from a stochastic to a genome-wide attractor profile: *A) Plot shows the whole expression profiles at 10min (x-axis) and 15min (y-axis) for the HRG response in MCF-7 cells. A box is constructed from the center of mass (i.e., average of whole expression), (CM(10min), CM(15min)) (black dot); a box contains gene expression within the range from CM(t*_*j*_*) - d to CM(t*_*j*_*) + d with a variable box size, d. To highlight the scaling of the Pearson correlation with box size d, r*_*d*_ *for d = 0.05, 0.1 and 0.2 are reported. The plot in the upper left corner shows that, between gene expression profiles, the Pearson correlation r*_*d*_ *follows a tangent hyperbolic function: r*_*d*_ = 0.97. *tanh* (6.79 · *d*-0.039) *(p<10^−4^), which reveals a critical transition in the correlation development.* *B) Stream plots for the box sizes in Panel A. These plots are generated from the vector field values {*Δ*x*_*i*_, *Δy*_*i*_*} at expression points {x*_*i*_*(10min), x*_*i*_*(15min)}, where Δx*_*i*_ = *x*_*i*_*(15min) - x*_*i*_*(10min), Δy*_*i*_= *x*_*i*_*(20min) - x*_*i*_*(15min), and x*_*i*_*(t*_*j*_*) is the natural log of the i*^*th*^ *expression: x*_*i*_*(t*_*j*_*) = ln(ε*_*i*_*(t*_*j*_*)) at t = t*_*j*_ *(t*_*j*_ = *10min or 15min; i = 1,2,.., N = 22,277). Blue lines represent streamlines and red arrows represent vectors at a specified expression point (plot every 2*^*nd*^, *6*^*th*^ *10*^*th*^ and *20*^*th*^ *point for d = 0.05, 0.1, 0.2, and the whole set, respectively). When we move from a small number of genes to the whole set, gene expression shifts from a stochastic to a genome-wide attractor profile.*

While by far the great majority of scientists have focused on the details of local gene-expression control, in this work we approach gene-expression regulation at the global level as an open thermodynamic (non-equilibrium) system by trying to answer some general questions:

- What is the underlying principle that regulates whole-genome expression through a global expression transition?
- Are there some differences among different biological systems regarding the global dynamics of genome expression?
- Is there a key player in the self-organization of expression?
- What is the mechanism of the self-organization that determines the change in the cell fate?

To address these important and still largely unanswered questions, we analyzed experimental transcriptome time-series of both microarray and RNA sequencing (RNA-Seq) data. We sought to demonstrate the presence of critical transitions in different biological processes associated with changes in the cell fate. We considered (i) early embryonic development in human and mouse, (ii) the induction of terminal differentiation in human leukemia HL-60 cells by dimethyl sulfoxide (DMSO) and all-trans-retinoic acid (atRA), (iii) the activation of ErbB receptor ligands in human breast cancer MCF-7 cells by epidermal growth factor (EGF) and heregulin (HRG), and (iv) T helper 17 cell differenation induced by Interleukin-6 (IL-6) and transforming growth factor-β (TGF-β) (**Methods**).

Our approach is based on an analysis of the dynamics of transcriptome data by means of the grouping (gene ensembles) of gene expression (averaging behaviors) built upon the results obtained in our recent papers [10,11] dealing with an MCF-7 cell population (see more in **Methods**). These previous studies revealed that self-organizing whole-genome expression coexisted with distinct response domains (critical states), where the self-organization exhibits criticality (critical behaviors) and self-similarity at a critical point (CP) - self-organized criticality control (SOC control) of overall expression.

To understand the current analysis based on our previous studies, it is important to elucidate the following points:

(i) In each critical state, coherent (collective/coordinated) behavior emerges in ensembles of stochastic expression by more than 50 elements [11]. Due to this coherent-stochastic behavior, *it is important to stress that the characteristics of the self-organization through SOC become apparent only in the collective behaviors of groups* with more than 50 genes in terms of their average value (mean-field approach).

(ii) SOC control of overall expression through a critical transition explains self-organization and the coexistence of critical states at a certain time point. This phenomenon cannot be interpreted in terms of the occurrence of a (first- or second-order) phase transition [19] in an equilibrium system, i.e., a phase transition in the overall expression from one critical state to another through a critical transition such as the ferromagnetic transition of iron at a critical temperature (*T*_c_) (*T*<*T*_c_: ferromagnetic-ordered magnetic moments; *T*>*T*_c_: paramagnetic-disordered). The coexistence of critical states demonstrates the existence of an internal order parameter such as the amount of gene-expression variance as in our formalism, but not of a control parameter such as temperature. In our previous work [10], use of the metaphorical example of the ferromagnetic transition to explain the self-organization of the coexistence of critical states could mislead regarding the true picture of SOC control.

(iii) Self-organization exhibits super-, near- and sub-critical states corresponding to the ensemble of high-, intermediate-, and low-variance gene expression, respectively, and their coherent oscillatory dynamics [10,11]. This difference in expression variance in critical states reveals the degree of cooperation of critical states based on the change in autocorrelation (Pearson correlation between neighboring time points in a critical state): highest in the super-critical state, medium in the near- critical state, and lowest (almost no change) in the sub-critical state (Figure 4A in [11]). This shows that stochastic perturbation in expression can spread in the sub-critical state, whereas the perturbation in the sub-critical state is locally confined in time.

(iv) Despite of considerable research over several decades, a general mathematical argument or formulation regarding the SOC hypothesis under non-equilibrium conditions is still in a primitive stage. This is due to the fact that there are a myriad of scenarios of self-organization with critical behaviors under non-equilibrium conditions, where “*a universal classification scheme is still missing for non-equilibrium phase transitions and the full spectrum of universality classes is unknown; it may be large or even infinite*.” [20]. Thus, to date, there is no stereotypic view of SOC in non-equilibrium systems (see more in subsection (iv) of **Discussion**).

In the present report, we describe the existence of a temporal interval (which differs for each system analyzed), where the change in transcriptome expression occurs via SOC at both single-cell (based on RNA-Seq data) and population levels (microarray data). Notably, the erasure of an initial SOC state, i.e., the disappearance of a sandpile-type critical behavior (criticality) of the initial state (*t* = 0 or initial cell state) determines *when and how a crucial change in the genome state occurs* (**sections I and II**), which intriguingly coincides with real biological critical events that determines the change in cell fate (**Discussion**).

SOC control occurs in a model-specific manner, which reveals that the spatio-temporal profiles of self-organization in overall expression regulation differ among the different tested systems; distinct critical states can coexist (**section III**). Furthermore, the emergent property of the coherent dynamics in critical states helps us to understand how the emergent sloppiness is exhibited in the genome-wide expression dynamics (**section IV**).

In **sections V and VI**, we demonstrate that a molecular stressor such as HRG in MCF-7 cells and DMSO in HL-60 cells, which induce cell differentiation, dynamically perturbs the genome-wide self-organization in SOC, and as a result, terminal cell fates occur at the end of a dissipative pulse-like global perturbation in self-organization. The perturbation of SOC occurs due to the exchange of expression flux among critical states through the cell nucleus environment as an open thermodynamic system. The quantitative evaluation of such flux flow reveals a mechanical picture of the interactions of critical states (“genome engine”; **Discussion**), and their roles in self-organization; most notably, sub-critical states (ensembles of genes with low-variance expression) are the central players for deriving the temporal development of self-organization. There is no fine-tuning by an external driving parameter to maintain critical dynamics in the SOC control of genome expression.

The elucidation of a statistical mechanism of the cell-fate change revealed through the perturbation of SOC in open thermodynamic gene regulation may contribute to new advances in our understanding of the dynamic aspects of epigenomics and help to clarify the material bases of biological regulation (**Discussion**).

## Results

The coherent dynamics of an ensemble of stochastic expression can be represented as a hill-like function, which is defined as a ‘coherent expression state (CES)’, with regard to the probability density profile in the regulatory space (expression vs. fold change in the expression) [10,11]. The emergence of a CES at around the critical point (CP) that marks the transition allows us to describe critical transitions in distinct cell types. Figure 3A shows, as an example, that, in DMSO-stimulated HL-60 cell differentiation, through the grouping of expression based on the fold change in expression (see why in **Methods**), a sandpile-type critical behavior is observed at 18-24h. These critical dynamics regarding the change in expression (e.g., fold change) emerges due to stochastic resonance [11].

**Figure 3:**
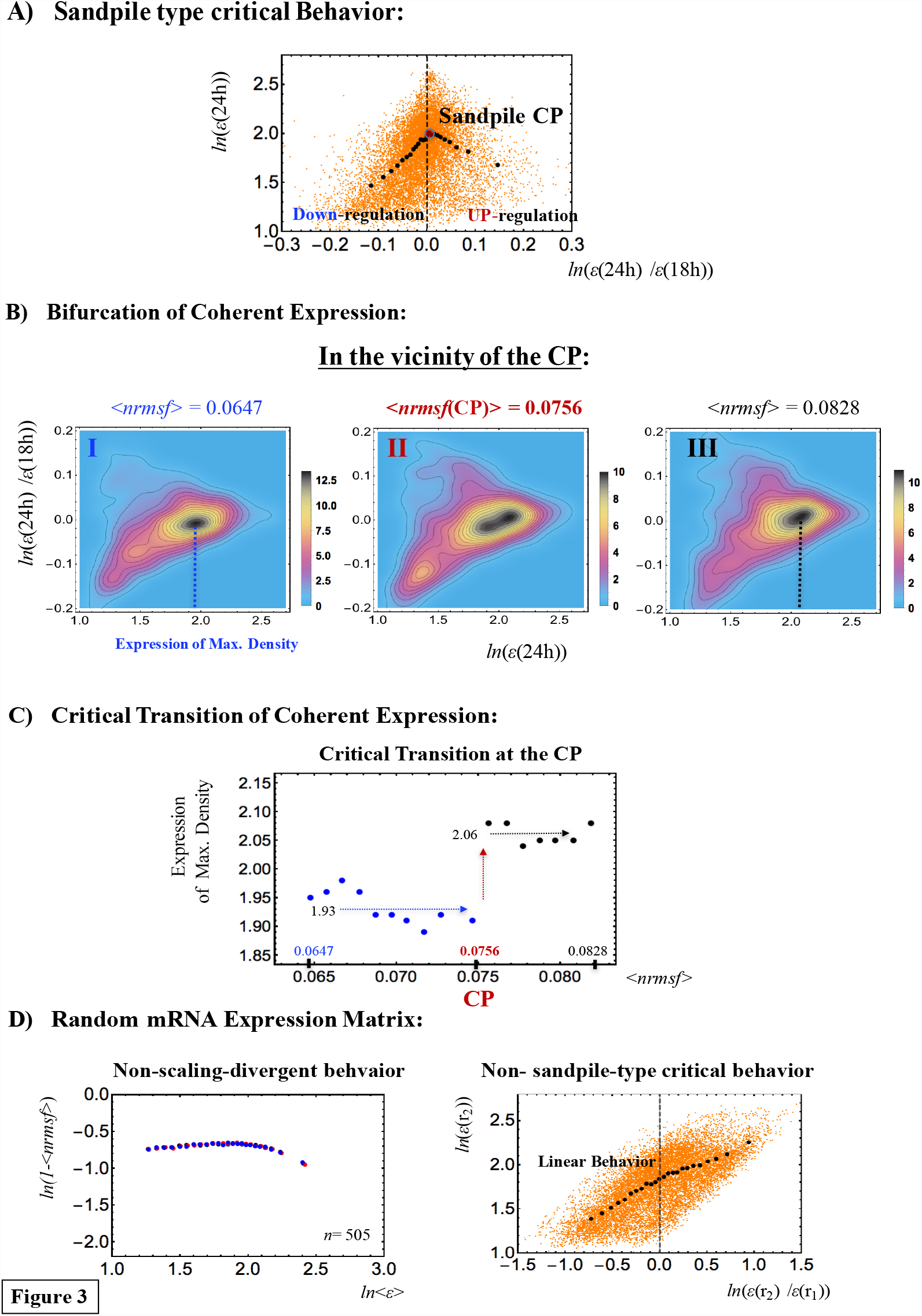
Self-organized criticality (SOC) in the DMSO-stimulated HL-60 cell fate: *A) The grouping of whole expression at 18-24h generates 25 groups with an equal number of 505 elements (mRNAs) according to the fold change in expression. A plot of the average value for each group in log-log space (x: fold change at 18-24h vs. y: expression at 24h) reveals a sandpile-type critical behavior at the critical point (CP), where the CP in terms of the ensemble (group) average (< >) occurs at the near-zero-fold change (x = 0; null expression change; x-axis) with <nrmsf(CP)> = 0.0756. Orange dots represent single mRNA expression in the background*. *B) The probability density function for the ensemble of expression (coherent expression state: CES) is shown in the regulatory space (x: expression at 24h vs. y: fold change in expression at18h-24h). Plots show that around <nrmsf(CP)> = 0.0756, a CES (highest density: x< 2.0) is annihilated and a new CES (highest density: x> 2.0) is bifurcated. The left to right panels show the sequence of the bifurcation-annihilation event: before (**I**: 0.060 <nrmsf< 0.070: <nrmsf> = 0.0647), onset (**II**: 0.071 <nrmsf< 0.081: <nrmsf> = 0.0756)) and after the event (**III**: 0.078 <nrmsf< 0.088: (<nrmsf> = 0.0828). Colored bars represent the probability density.* *C) The corresponding plot (B) reveals that a step function-like critical transition occurs at <nrmsf(CP)> = 0.0756 in the space (x: <nrmsf> of coherent state vs. y: its expression of the highest density at 18h). Blue and black arrows represent average values of the density trends before and after the transition, respectively. The plot shows, in the vicinity of the CP, the occurrence of a self-similar bifurcation (symmetry breaking) in the expression profile to that of the overall expression (see **section III**).* *D) Random mRNA DMSO expression matrix reveals that, in this case, a CP does not exist. This is confirmed by anomalous features of the corresponding SOC (**Methods**): non-scaling-divergent (3 different time points are shown by colors) and non-sandpile critical behavior of random expression between two different time points (orange dots: single random expression). The random matrix is made by randomly selecting each matrix component (i,j) from the original DMSO expression matrix (12625 expression (i) times 13 time points (j)). We observed similar linear correlative behaviors for other cells in both microarray and RNA-Seq data.*

Around the critical behavior in terms of *nrmsf* (normalized root mean square fluctuation, see **Methods**), symmetry breaking corresponds to the annihilation of a CES at a lower expression level and the bifurcation of another CES at a higher expression level through a flattened profile in the regulatory space (Figure 3B). The maximum density of the coherent state follows a step functional like the critical transition at the CP (Figure 3C). The critical transition of coherent expression in the vicinity of the CP shows self-similar behavior to that of the overall expression (see **section III**) in DMSO-stimulated HL-60 cells, i.e., the sandpile-type critical point exhibits a critical transition (called sandpile type critical transition). We also observed a sandpile-type critical transition at a CP in HRG-stimulated cell differentiation [11].

Next, when the DMSO-induced expression matrix is randomly shuffled, no sandpile-type CP is present (Figure 3D: right panel): the fold change scales almost linearly with the logarithm of the level of expression. The corresponding frequency distribution changes from non-Gaussian (Figure 9B) to Gaussian (Figure 9C) according to *nrmsf*. This shows that (i) gene expression becomes random due to the random shuffling of gene expression, and (ii) such *randomized expression destroys the sandpile-type critical behavior* seen in Figure 3A.

Hence, we arrive to the conclusion that it is crucial to investigate the existence of a critical point or criticality for SOC control in overall expression.

## I. Perturbation of SOC control and the Genome-State Change

We investigated the occurrence of critical transitions in distinct cell types. First, we examined whether the sandpile-type critical transition (see **Methods**) occurs around the zero-fold change (i.e., null change in expression) between different time points. The critical point (CP), at the top of the sandpile, corresponds to a group of genes that show almost no (average) change in expression. Next, we assessed whether or not the CP is a fixed point in time by evaluating if the average *nrmsf* value of the CP group changes over time. The basic hypothesis that justifies the choice of *nrmsf* as a metric for evaluating gene-expression dynamics is that gene expression groups scale with topology-associated chromatin domains (TADs) [21-25]. The degree of gene expression normalized fluctuation is presented with respect to chromatin remodeling, i.e., *nrmsf* is expected to be associated with the physical plasticity of genomic DNA and the high-order chromatin structure [11]. Thus, we can expect that, as the flexibility of a given genome patch increases, so should the *nrmsf* of the corresponding genes. To confirm this further, the scaling-divergent behaviors (Figure 4 and **Methods**) of *nrmsf* and average expression, another important feature of SOC, may show a quantitative relation with the aggregation state of chromatin. In **Supplementary File S1**, we show that gene expression exhibits collective behavior (as coherent-stochastic behavior: CSB) in the power-law scaling through interactions among genes.

**Figure 4:**
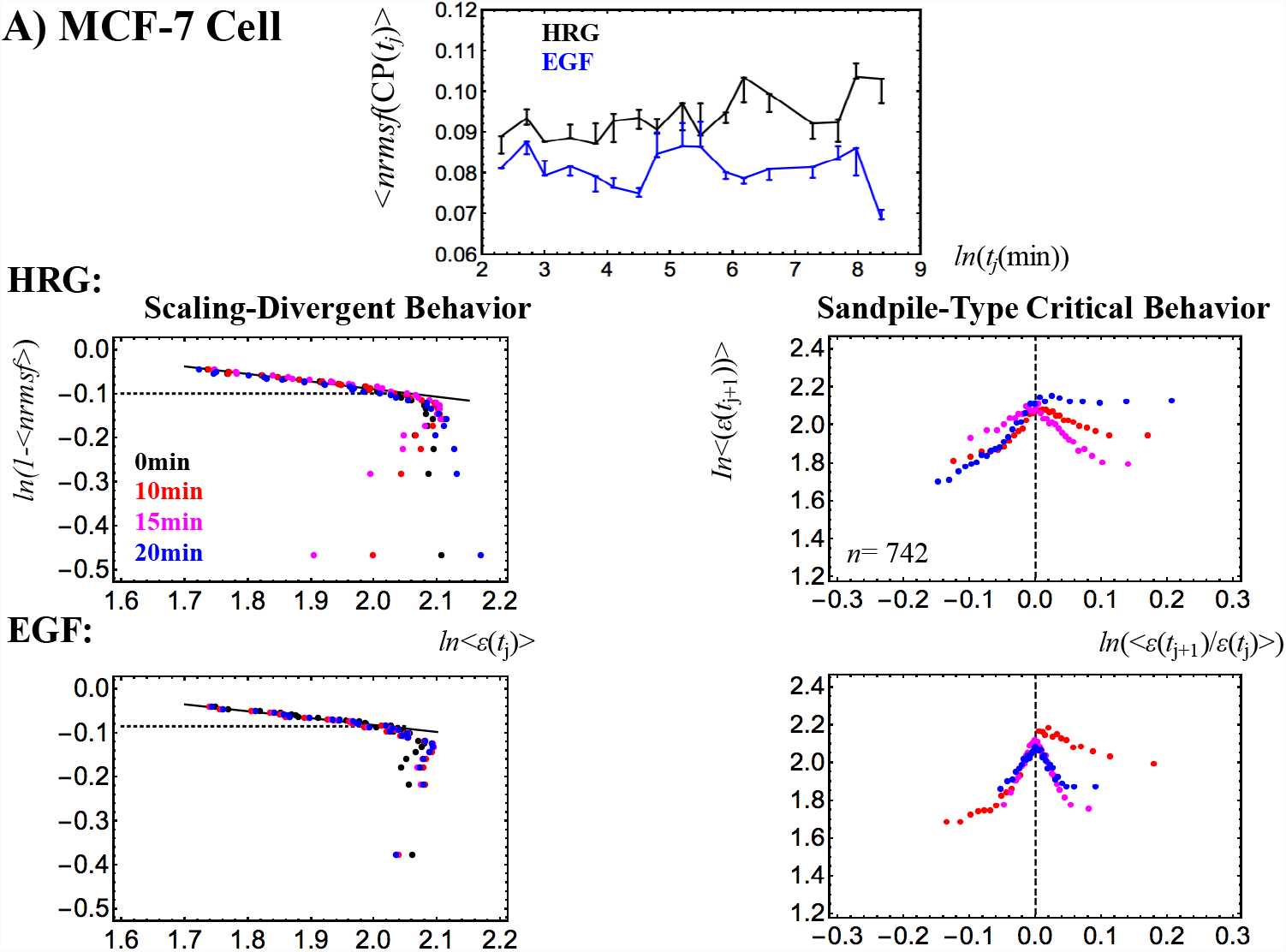
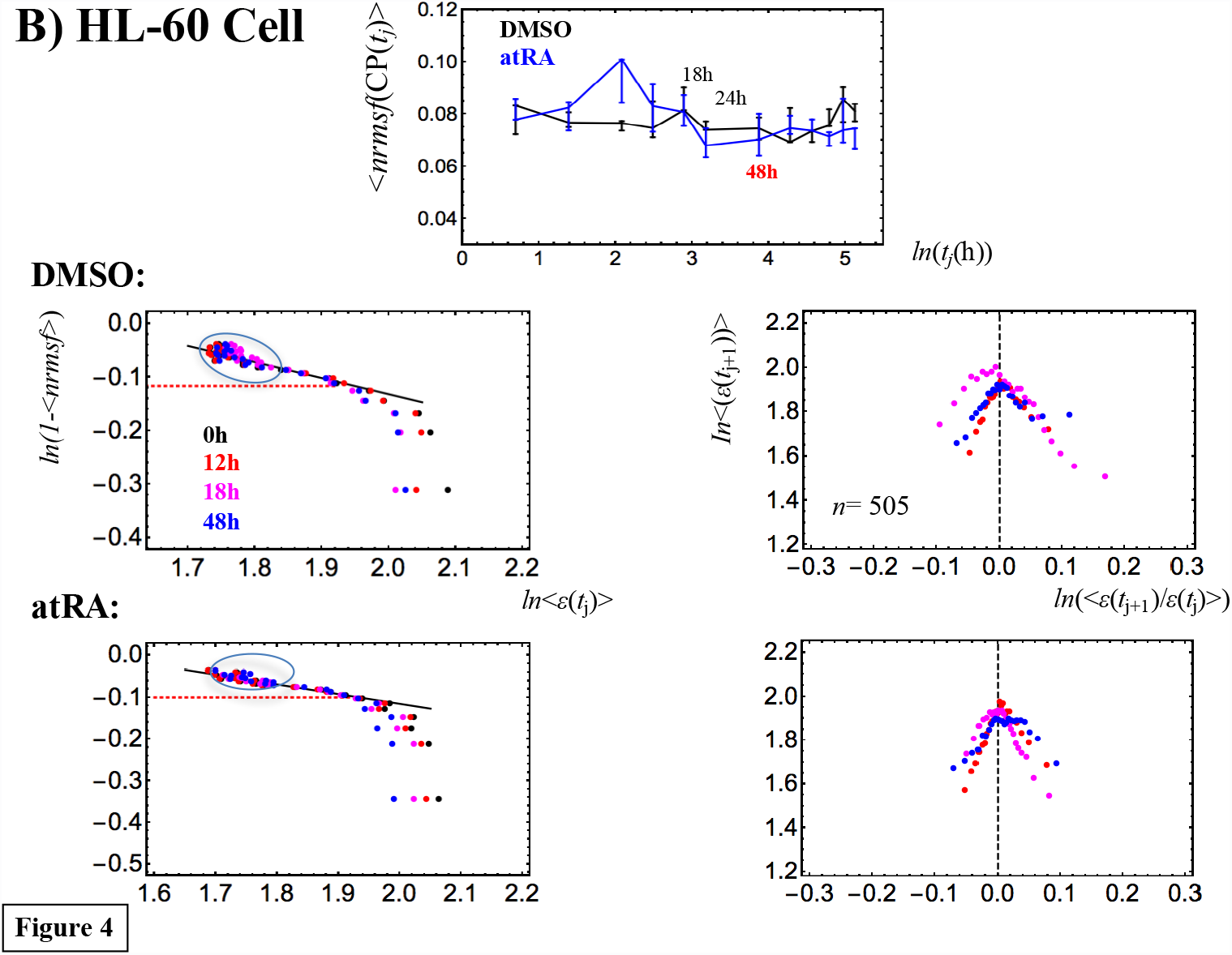
Time-development of the characteristic behaviors of SOC: *A) MCF-7 cells and B) HL-60 cells. At each experimental time point, t*_*j*_ *(A: 18 time points and B: 13 points; **Methods**), the (ensemble) average nrmsf value of the CP at t = t*_*j*_, *<nrmsf(CP(t*_*j*_*))> is evaluated at the sandpile-type critical point (top of the sandpile: right panels). In the top center panels, <nrmsf(CP(t_j_))>, is plotted against the natural log of t*_*j*_ *(A: min and B: hr). Error bar represents the sensitivity of <nrmsf(CP(t*_*j*_*))> around the CP(t*_*j*_*), where the bar length corresponds to the change in nrmsf in the x-coordinate (fold-change in expression) from x(CP(t*_*j*_*)) − d to x(CP(t*_*j*_*)) + d for a given d (A: d = 0.005; B: d =0.01; due to much more mRNAs in MCF-7 cells). Temporal averages of <nrmsf(CP(t*_*j*_*))> are A)* 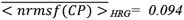 *and* 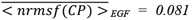 *and B)* 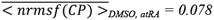, *where an overbar represents temporal average. Note: An overbar for a temporal average is used when ensemble and temporal averages are needed to distinguish*. *A) MCF-7 cells: The temporal trends of <nrmsf(CP(t*_*j*_*))> are different for HRG and EGF. The onset of scaling divergence (left panels: second and third rows) occurs at around* 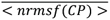 *(black dashed line), and reflect the onset of a ‘genome avalanche’ (**Methods**)*. *B) HL-60 cells: The trends of <nrmsf(CP(t*_*j*_*))> for the responses to both DMSO (black line) and atRA (blue) seem to be similar after 18h (i.e., global perturbation; see **section VI**). The scaling-divergent behaviors for both DMSO and atRA reveal the collapse of autonomous bistable switch (ABS [11]) exhibited by the mass of groups in the scaling region (black solid cycles) for both DMSO and atRA. The onset of divergent behavior does not occur around the CP* 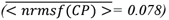, *but rather is extended from the CP (see **section III**).* *The power law of scaling behavior in the form of 1- <nrmsf> = α<ε>^-β^ is:* *A) α = 1.29 & β = 0.172 (p< 10^−10^) for HRG, and α = 1.26 & β = 0.157 (p<10^−9^) for EGF;* *B) α = 1.60 & β = 0.301 (p< 10^−6^) for DMSO, and α = 1.42 & β = 0.232 (p< 10^−6^) for atRA.* *Each dot (different time points are shown by colors) represents an average value of A) n = 742 mRNAs for MCF-7 cells, and B) n = 505 mRNAs for HL-60 cells.*

A transcriptome analysis based on a mean-field (grouping) approach (**Methods**) reveals that sandpile transitions occur and the position of the CP exhibits time-dependence in terms of *nrmsf*, which reflects the temporal development of SOC, i.e., the CP is not a fixed point (Figure 4). Interestingly, the CP of the initial state disappears over time, suggesting the occurrence of a crucial change in the genome state (Figure 5). Regarding critical transitions around the CP (Figure 6), different types of dynamical bifurcation or annihilation of a characteristic coherent expression state occur around the CP in different cell models:

1. Unimodal-flattened-bimodal transition for MCF-7 cell-HRG and -EGF models, and
2. Unimodal-flattened-unimodal transition for HL-60 cell-atRA and -DMSO;

both models point to symmetry breaking in the expression profile.

**Figure 5:**
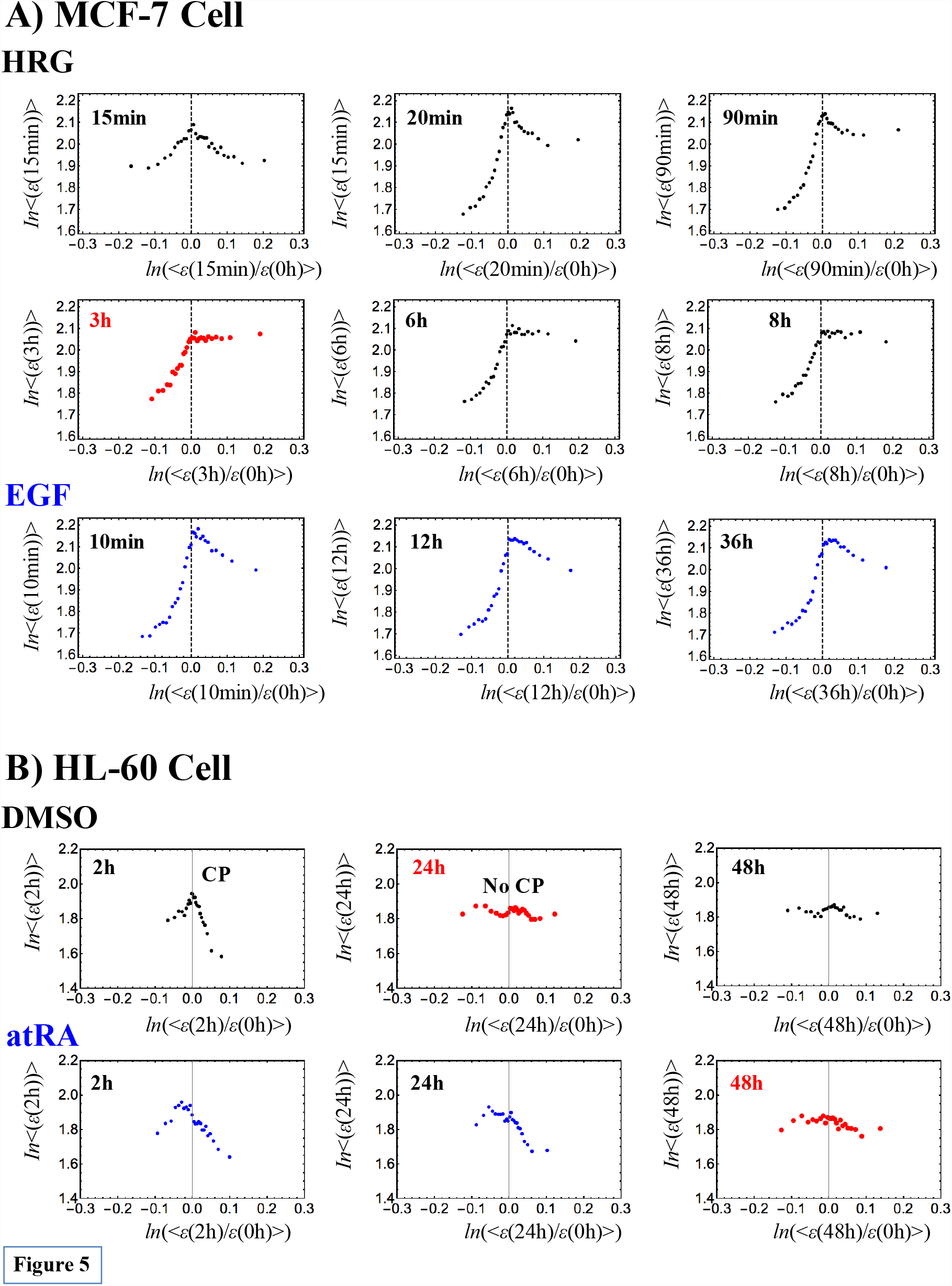
Genome-state change revealed through erasure of the initial-state criticality in overall expression (cell population level): *The grouping of overall expression at t= t*_*j*_ *(j ≠ 0) according to the fold change in expression from the initial overall expression (t= 0) shows how the initial-state critical behavior is erased over time, i.e., a sandpile profile in overall expression is destroyed at t = t*_*j*_ *from t= 0. This event points to the time when the genome-state change occurs. On the x-axis, ln(<ε(t)/ε(0h)>) (t: min or hr) represents the natural log of the ensemble average (< >) of the fold change in expression, ε(t)/ε(t= 0h), and on the y-axis, ln(<ε(t)>) represents the natural log of the ensemble average of expression, <ε(t)>.* *A) MCF-7 cells: In HRG-stimulated cells (black), the divergent behavior in up-regulation is no longer observed, and the same shape is apparent after 3h. This suggests that the genome-state change occurs at 3h (red) through erasure of the initial-state up-regulation process (partial erasure). In contrast, in EGF-stimulated cells (blue), almost the same sandpile profile remains for up to 36h, which suggests that no genome-state change occurs (see the local perturbation in Figure 14).* *B) HL-60 cells: A CP is erased at 24h (red) and 48h (red) in DMSO- (black) and atRA-stimulated (blue) cells through the disappearance of divergent behaviors in both up- and down-regulation of the initial state (full erasure), respectively. Furthermore, these plots suggest that the epigenomic states in DMSO- and atRA-stimulated cells become the same after 48h. Note: The Pearson correlations of overall expression between different time points are near-unity (Figure 1A).* *Each dot represents the average value of A) n = 742 mRNAs for MCF-7 cells, and B) n = 505 mRNAs for HL-60 cells.*

**Figure 6:**
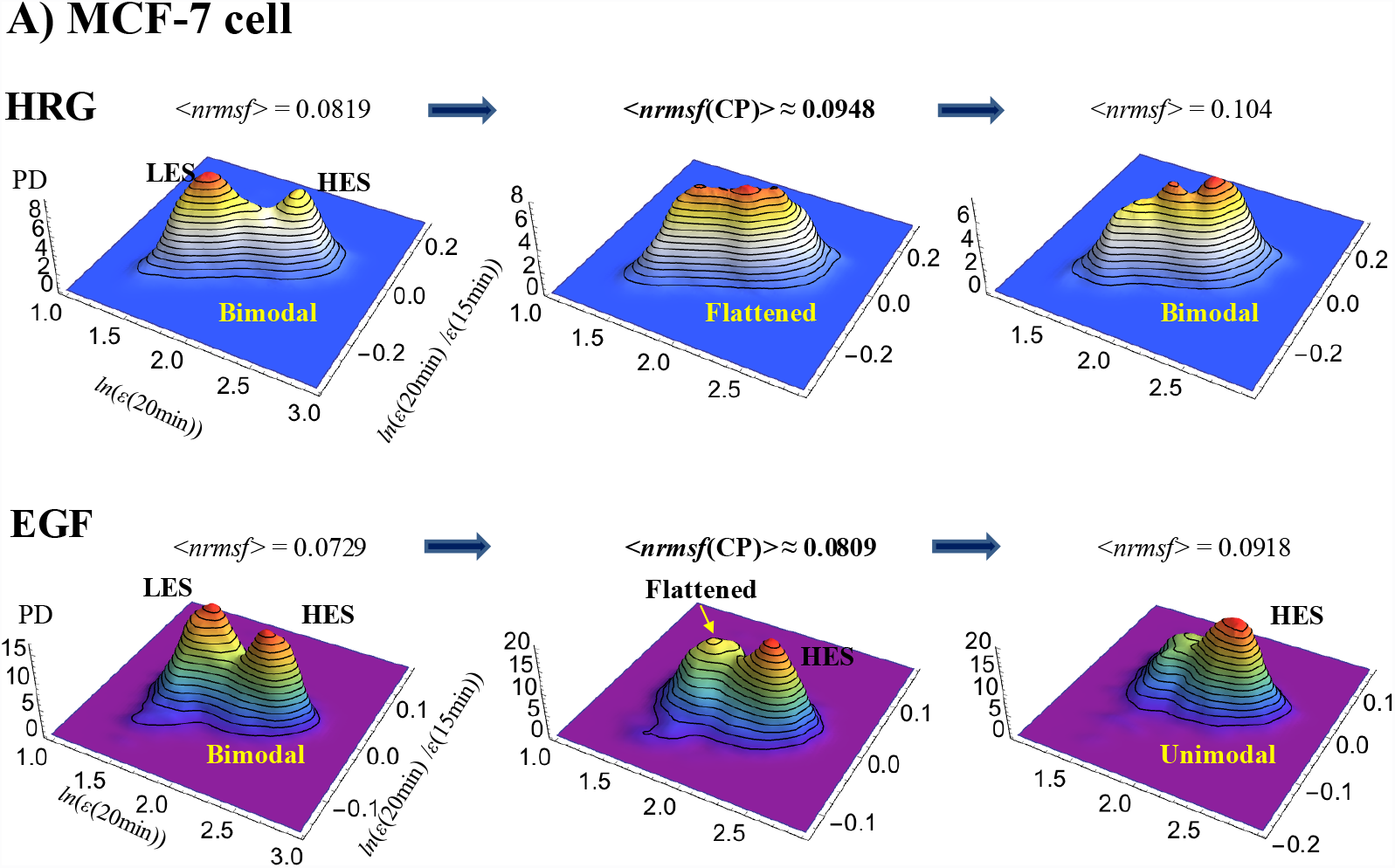
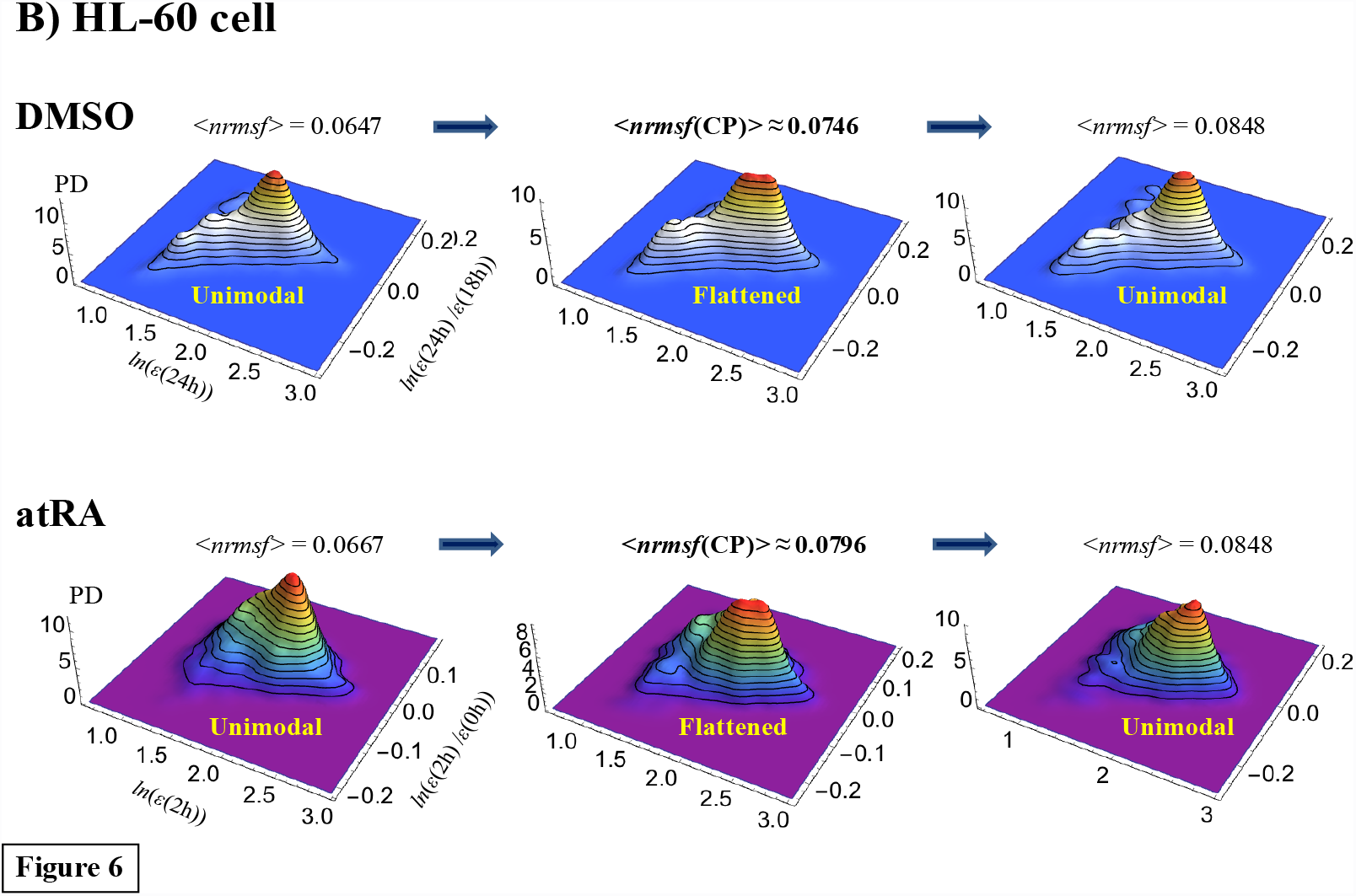
The self-similar bifurcation or annihilation of a characteristic coherent expression state in the vicinity of a critical point in different cell types. *A coherent expression state (CES), which contains approximately 1000 expression points, is bifurcated or annihilated around the CP through nrmsf grouping. The average nrmsf of an expression group, <nrmsf>, is evaluated for a variable range: x + 0.01(m-1) <nrmsf< x + 0.01m (integer, m≤10 for each x = 0.7, 0.8 and 0.9). The probability density function (PDF) in the regulatory space (z-axis: probability density) around <nrmsf(CP)> shows:* *A) MCF-7 cells: In the HRG response at 15-20min, around <nrmsf(CP)> (<nrmsf(CP)>*_*HRG*_*= 0.094 and <nrmsf(CP)>*_*EGF*_ *= 0.081;* Figure 4A), the PDF exhibits a bimodal-flattened-bimodal transition, in which a bimodal profile points to the existence of two CESs: one represents a low-expression state (LES) and the other represents a high-expression state (HES); the valley defines the boundary between low and high expression [10]. In the EGF response (second row), above <nrmsf(CP)>, a low-expression state (LES) is annihilated and only a high-expression state (HES) exists, i.e., a unimodal-bimodal transition is present. In the HRG response, at a time period other than 15-20min, a unimodal-bimodal transition occurs [10]. This result shows the self-similar symmetry-breaking event to the overall expression for MCF-7 cells, even at a transient change in the HRG response: a bimodal-flattened-bimodal transition at 15-20 min. *B) HL-60 cells: A unimodal-flattened-unimodal transition occurs at 18-24h (pseudo-3-dimensional PDF plots of Figure 3B) for DMSO and at 0-2h for atRA, which again reveals the self-similarity to the overall expression for HL-60 cells (**section III**).*

These results offer the following insights:

1. **SOC control in different cell models:** The self-similarity of symmetry breaking around the CP suggests the existence of critical states (distinct transcription response domains) in different cell models (see **section III**). Furthermore, the self-similarity in the overall expression suggests the occurrence of i) a unimodal-flattened-bimodal transition for MCF-7 cell fates, and ii) a unimodal-flattened-unimodal transition for HL-60 cell fates, which will be shown to be robust over time (**section III**), except for a transient change (see Figure 6A, an example in the HRG response: a bimodal-bimodal transition at 15-20 min due to a bifurcation in the unimodal profile; refer also to Figure 7B in [10]). The flattened distribution shows that the degree of cooperation in expression regulation, i.e., the strength of the correlation around the CP, tends to increase with the size of the system ensemble, as expected in the critical dynamics of biological systems (see more in subsection (iv) of **Discussion**). The unimodal-unimodal transition in HL-60 cell fates stems from the fact that a pair of two coherent expression states (a bimodal expression profile as an autonomous bistable switch [11]) collapses to a single coherent state in the scaling region (low *nrmsf* region: Figure 4B).
2. **Perturbation of SOC control**: The temporal development of SOC control reflects the presence of dynamic changes in critical states in terms of both exchanging of genes between critical states and changes in the expression profile. This implies the perturbation of SOC control through the interaction between critical states (see **sections V and VI**). Interestingly, regarding HL-60 cell fates, at 12-18h, a pulse-like global perturbations involving the regulation of critical states occur for the responses to both DMSO and atRA (**section VI**). After 18h, the temporal trends of the CP(*t*_j_) for DMSO and atRA in terms of *nrmsf* become similar (Figure 4B). Note that the bifurcation-annihilation events of CES around the CP for both DMSO and atRA at 24-48h become almost identical (data not shown). This shows that the dissipation of the stressor-specific perturbation in HL-60 cells drives the cell population toward the same attractor state [26,27].
3. **Genome-state change:** Critical dynamics appear in the change in expression between different time points. Thus, in the change in expression at *t*_0_-*t*_j_ (*t*_0_<*t*_j_), the erasure of the criticality of the initial state (Figure 5) at *t* = *t*_j_ indicates that a genome-state change occurs: at 3h in HRG-stimulated MCF-7 cells, and at 24h and 48h in DMSO- and atRA-stimulated HL-60 cells. These HL-60 genome-state changes further confirm that both DMSO- and atRA-stimulated HL-60 cells converge toward the same global gene-expression profile at 48h. Erasure of the initial-state critical behavior occurs in different cell types; divergent behavior in up-regulation (a partial erasure of criticality) disappears in HRG-stimulated MCF-7 cells, whereas the full erasure of criticality occurs in both DMSO- and atRA-stimulated HL-60 cells. As demonstrated in **section VI**, these genome-state changes occur after dissipative pulse-like global perturbations in SOC control (at 12-18h for HL-60 cells and at 15-20min for HRG-stimulated MCF-7 cells). In contrast, as a proof of concept, the genome-state change does not occur in EGF-stimulated MCF-7 cell (Figure 5; refer also to local perturbation; **section VI**), which is consistent with cell proliferation (and the absence of differentiation) in the EGF response [28,29]. Note: The independence of the choice of the initial state at *t* = *t*_0_ for the breakdown of sandpile type criticality at *t* = *t*_b_ (condition: *t*_0_<*t*_b_) further confirms the timing of the genome-state change. Furthermore, we observed that the time point (*t*_0_) of the initial state is earlier or equal to the onset of the pulse-like global perturbation (12-18h) for DMSO-stimulated HL-60 cells (*t*_0_ ≤12h; see Supplementary Figure S1); after the global perturbation, this independence does not hold, which suggests that a pulse-like global perturbation may be related to the first stage of cell-fate determination (process for autonomous terminal differentiation; see subsection (ii) in **Discussion**). The changes in the genome state coincide with real biological critical events that determines the cell-fate change (see subsections (i)-(iii) in **Discussion**).

**Figure 7:**
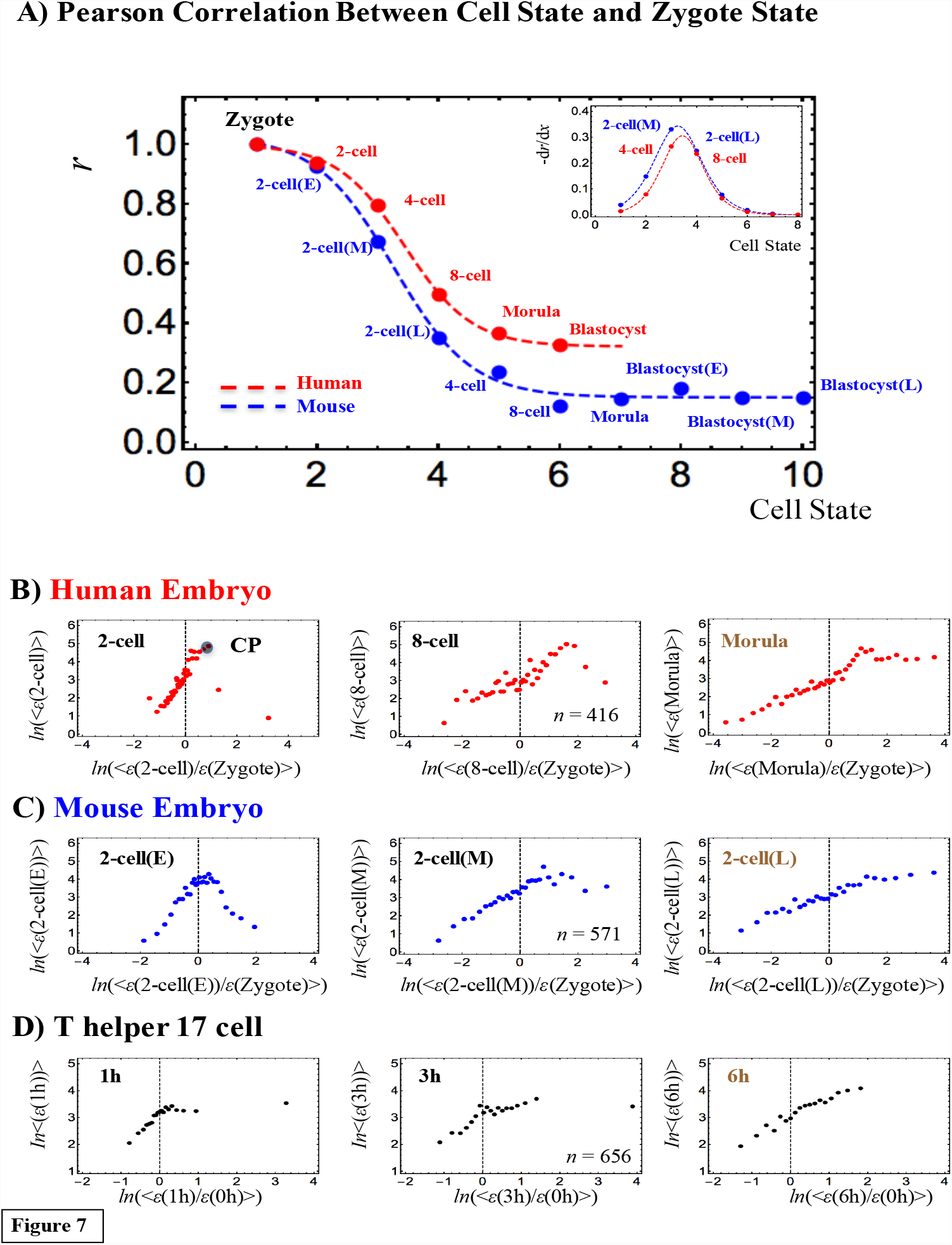
Genome-state change revealed through erasure of the initial-state critical point in overall expression (single cell level): *Transcriptome RNA-Seq data (RPKM) analysis for A-C) human and mouse embryo development, and D) T helper 17 cell differentiation* *A) In both human and mouse embryos (red: human; blue; mouse; refer to development stages in **Methods**), a critical transition is seen in the development of the overall expression correlation between the cell state and the zygote single-cell stage, with a change from perfect to low (stochastic) correlation: the Pearson correlation for the cell state with the zygote follows a tangent hyperbolic function: a-b· tanh(c·x-d), where x represents the cell state with a= 0.59, b= 0.44 c= 0.78 and d = 2.5 (p< 10*^*−3*^*) for human (red dashed line), and a= 0.66, b= 0.34 c= 0.90 and d = 3.1 (p< 10*^*−2*^*) for mouse (blue). The (negative) first derivative of the tangent hyperbolic function, -dr/dx, exhibits an inflection point (zero second derivative), indicating that there is a phase difference between the 4-cell and 8-cell states for human, and between the middle stage and late 2-cell states for mouse (inset); a phase transition occurs at the inflection point. Notably, the development of the SOC control in the development of the sandpile-type transitional behaviors from the zygote stage (30 groups; n: number of RNAs in a group: **Methods**) is consistent with this correlation transition:* *B) In human, a sandpile-type CP (at the top of the sandpile) disappears after the 8-cell state, and thereafter there are no critical points. This shows that the zygote SOC control in overall expression (i.e., zygote self-organization through criticality) is destroyed after the 8-cell state, which indicates that the memory of the initial stage of embryogenesis is lost through a stochastic pattern as in the linear correlation trend (refer to the random expression matrix in Supplementary Figure S1 or to Figure 3D). The results suggest that reprogramming (massive change in expression) of the genome occurs after the 8-cell state.* *C) In mouse, a sandpile-type CP disappears right after the middle stage of the 2-cell state and thereafter a stochastic linear pattern occurs, which suggests that reprogramming of the genome after the middle stage of the 2-cell state destroys the SOC zygote control.* *D) In Th17 cell differentiation, a sandpile-type CP disappears after 3h through a stochastic linear pattern. Therefore, the plot suggests that the genome-state change occurs at around 3-6h in a single Th17 cell.*

**Supplementary Figure S1:**
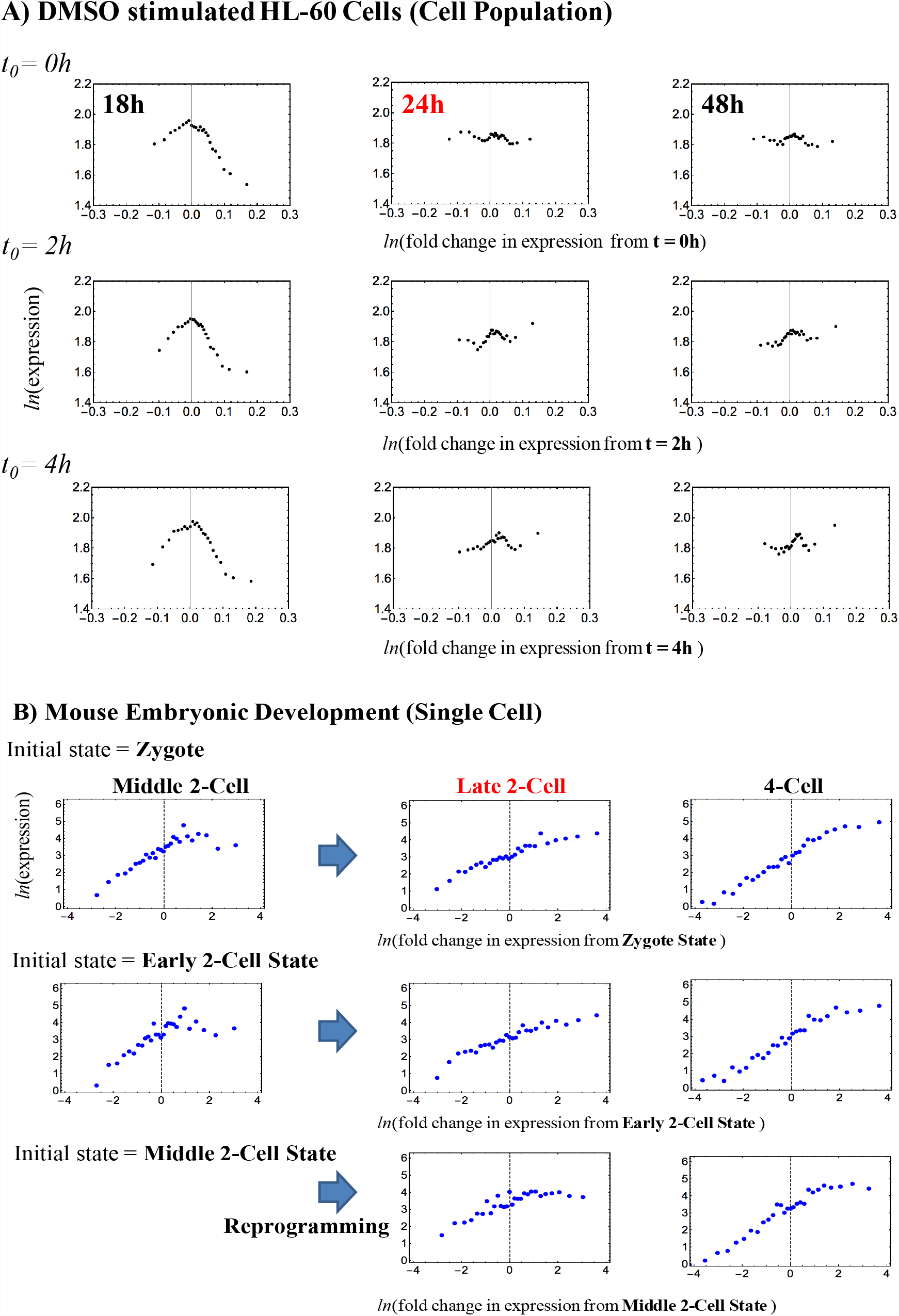
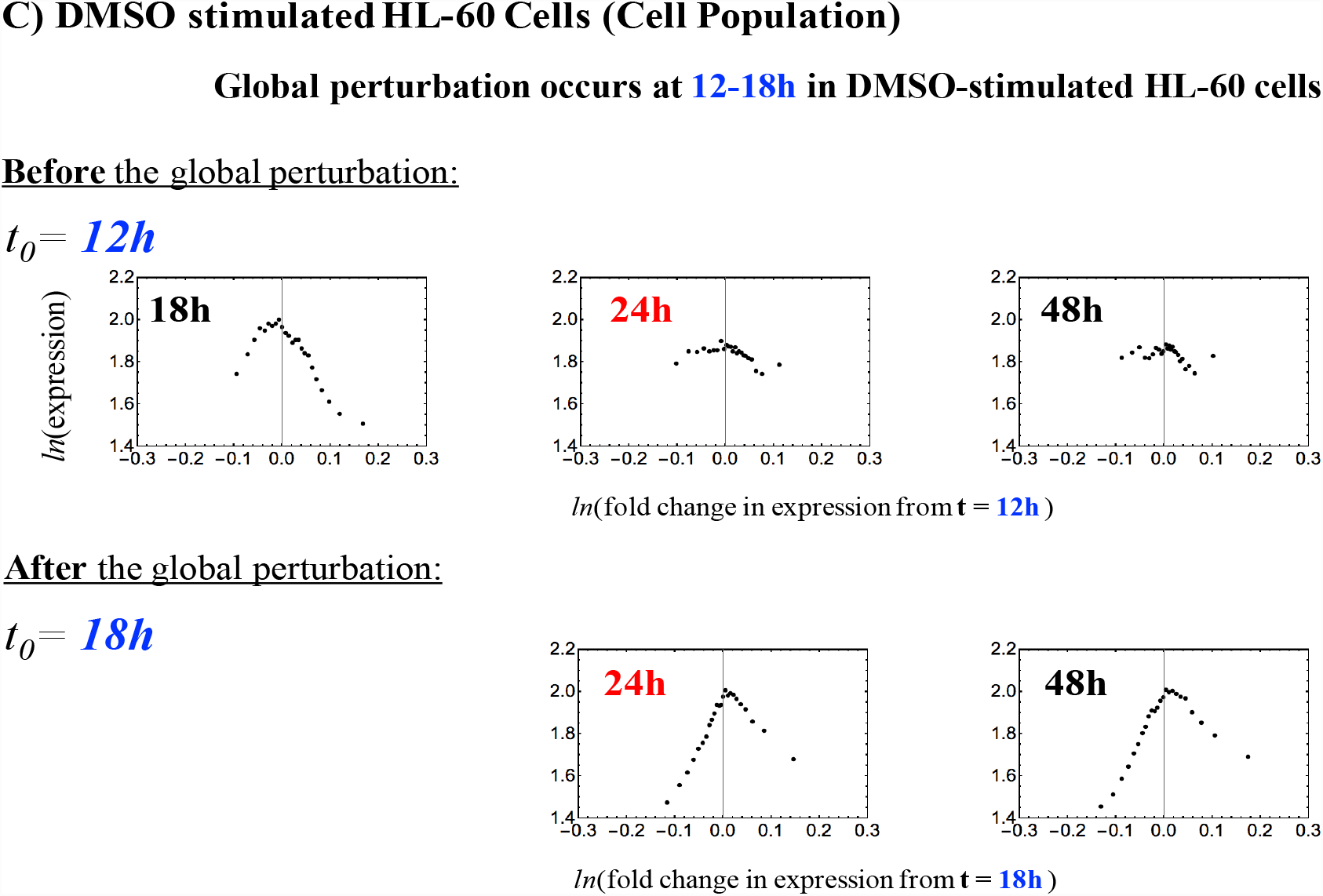
*The timing of the genome-state change occurs at the erasure of an initial-state criticality. This figure demonstrates that the timing of the genome-state change does not vary with the choice of an initial state (t*_*0*_*) for A) DMSO-stimulated HL-60 cells (cell population; microarray data) or B) mouse embryonic development (single cell; RNA-Seq data). This result confirms the occurrence of the genome-state change at 24h for DMSO-stimulated HL-60 cells (see more section I), and of reprogramming after the middle 2-cell state for mouse embryonic development (see more **section II**). C) A pulse-like global perturbation in self-organization occurs at 12-18h (Figure 14: DSMO) in DMSO-stimulated HL-60 cells. The breakdown of criticality occurs at 24h for the initial state at t*_*0*_ *= 12h, before the perturbation, but does not occur for the initial state at t*_*0*_ *= 18h, after the perturbation. This suggests that the global perturbation can be related to the first stage of cell-fate determination (process for autonomous terminal differentiation; see subsection (ii) in **Discussion**).*

To dig deeper into these findings, in the following sections we investigate the transcriptome SOC control in embryonic development at a single-cell level based on next-generation RNA sequencing data, and address the mechanism of the genome-state change in terminal cell fates revealed through the perturbation of genome-wide self-organization.

## II. Control of the Embryonic Development of a Single Zygotic Cell and T helper 17 Cell Differentiation through Self-Organized Criticality

The RNA-Seq approach allows us to check the tenability of SOC control at a single-cell level. Here we analyze RNA-Seq data related to immune cell differentiation, and human and mouse embryonic development.

Figure 7A shows Pearson correlations between gene expression profiles related to the zygote and early embryo single-cell states. We previously demonstrated (see Figure 1) that the between-profiles Pearson correlation follows a tangent hyperbolic function with an increase in the number of genes. A similar transitional behavior (the presence of a critical transition) is observed in both human and mouse early embryo development. This transition takes place between the 4-cell and 8-cell states in human and between the middle and late 2-cell states in mouse.

Notably, the correlation transition corresponds to the onset of the breakdown of SOC control in early embryo development according to the number of cell states starting from the zygote single-cell state. In both human and mouse embryos, the maternal SOC zygote controls break down through cell-state development, and the overall expression becomes stochastic. In human, the maternal SOC zygote control survives until the 8-cell state stage. After the morula state (Figure 7B), no sandpile-type critical point exists, and the overall gene-expression profile is fully stochastic compared with the overall expression in the zygote (see Supplementary Figure S2), which indicates that the memory of genome expression in the zygote is lost in the morula state. In contrast, in the mouse embryo (Figure 7C), the maternal SOC controls survive from the zygote to the 2-cell state (middle stage).

The breakdown of early SOC zygote control in overall expression indicates that significant global perturbation (refer to **section VI**) occurs to destroy the SOC zygote control in early embryo development. The human scenario may be explained by the known fact that the genome of the human embryo is not expressed until the 4-8 cell stage, which suggests that there is no apparent significant perturbation as reprogramming in early human embryo development (see more in **Discussion**). The timing of reprogramming is further confirmed by the independence of the breakdown of criticality from the choice of the initial cell-state (see Supplementary Figure S1).

Along similar lines of reasoning, T helper 17 (Th17) cell differentiation shows that initial SOC control (*t* =0) is destroyed after 3h (Figure 7D), which reveals that the Th17 genome-state changes at around 3-6h after the induction. Sandpile criticality emerges again after 6h in Th17 cell differentiation (see Supplementary Figure S2). The embryo and immune cell results confirm the presence of specific SOC controls, not only in large cell populations, but also at a single-cell level.

Next, we examine the development of SOC control between sequential cell states. As shown in Figure 8, from the zygote to the morula state in mouse, one sandpile-type critical regulation transitions to another through a non-critical transition of regulation (absence of critical behaviors: non-SOC control) in the middle-late 2-cell state. This confirms that reprogramming of the early mouse embryo cells from the zygote destroys SOC control to initiate self-organization in the new embryonal genome at the late 2-cell state, which exhibits a stochastic overall expression pattern (Figure 7C). Thereafter, the SOC control again takes over embryo development. Non-SOC control exhibits almost a linear behavior, which is a characteristic of randomized expression (see Supplementary Figure S2 and refer to the similar linear behavior in Figure 3D in a different cell type).

**Figure 8:**
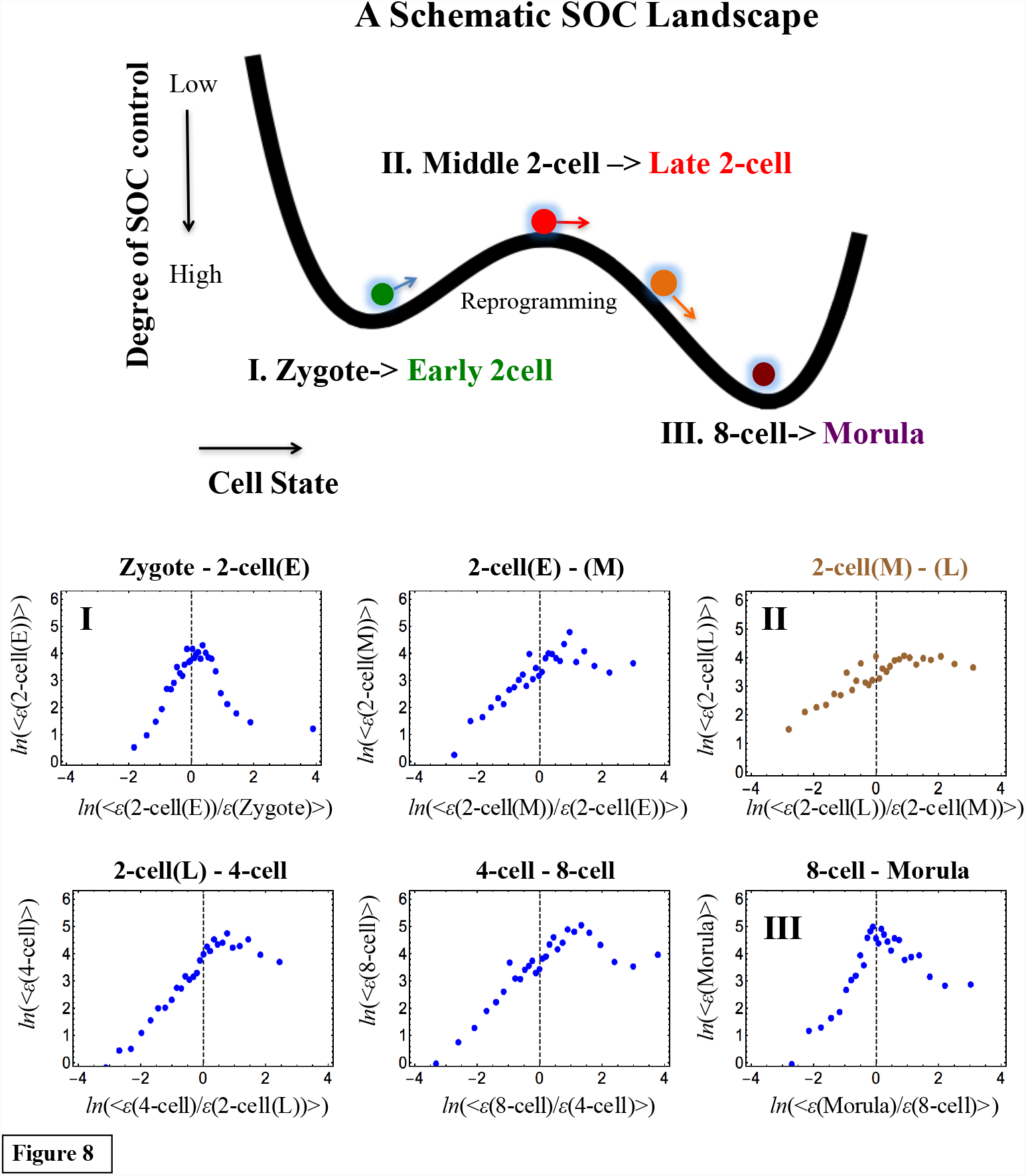
The SOC control landscape as revealed by a sandpile-to-sandpile transition in mouse embryo development. *The development of a sandpile transitional behavior between sequential cell states suggests the existence of an SOC control landscape (first row: schematic picture of a valley-ridge-valley transition; x-axis: cell state; y-axis: degree of SOC control). The second and third rows show that a sandpile-to-sandpile transition occurs in mouse embryo development from the zygote single-cell stage to the morula cell state: **I**: a sandpile (i.e., critical transition) develops from the zygote single-cell stage to the early 2-cell state => **II:** a sandpile is destroyed from the middle to the late 2-cell state, which exhibits stochastic expression (i.e., no critical transition; refer to the random mouse expression matrix in Supplementary Figure S2) => **III:** a sandpile again develops from the 8-cell state to the morula state. These results show that a significant perturbation (reprogramming) in self-organization occurs from the middle stage to the late 2-cell stage through a stochastic overall expression (refer to Figure 7). Note: Qualitatively, there is a high degree of SOC control for a well-developed shape of the sandpile-type transition (SOC control), an intermediate degree of SOC control for a weakened (broken) sandpile, and a low degree for non-SOC control. The latter is due to stochastic expression. The linear behavior (absence of a critical point) in mouse embryo development is also reflected in the low Pearson correlation (r ~0.21 after the 8-cell state) (Figures 7A, C).*

The transition of SOC control through non-SOC control suggests that an SOC-control “landscape”, i.e., a valley (SOC control) - ridge (non SOC control) - valley (SOC control), is seen in early mouse embryo development. The genome expression dynamics in the early embryo, through the development of SOC control, are consistent with the ‘epigenetic landscape’ frame, in the broad terms of the global activation-deactivation dynamics of the genome generally consistent with the DNA de-methylation-methylation landscape [30].

The onset of the genome-state change (at the breakdown of initial-state SOC control) exhibits a clear difference between single cells and a cell population (Figure 5): cell populations do not exhibit a stochastic pattern, in contrast to single cells in early human and mouse embryonic development (Figure 7 and Supplementary Figure S2). The stochastic pattern is confirmed by low Pearson correlation between the zygote and early embryo single-cell states at the onset (*r*< 0.5; Figure 7A), whereas in cell populations, the Pearson correlation for overall expression at any different time points is close to unity (Figure 1A and Figure 2B).

**Supplementary Figure S2:**
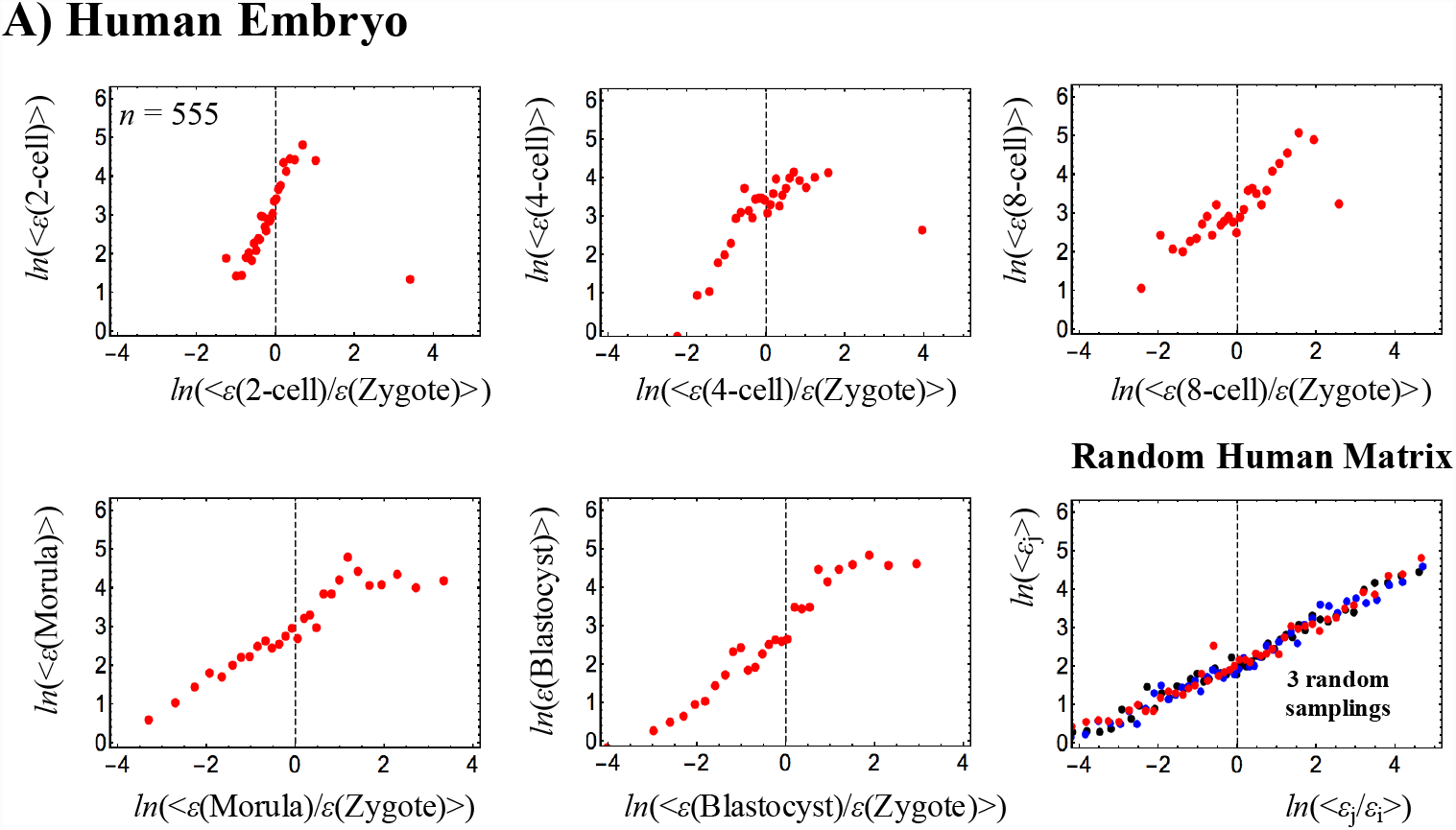
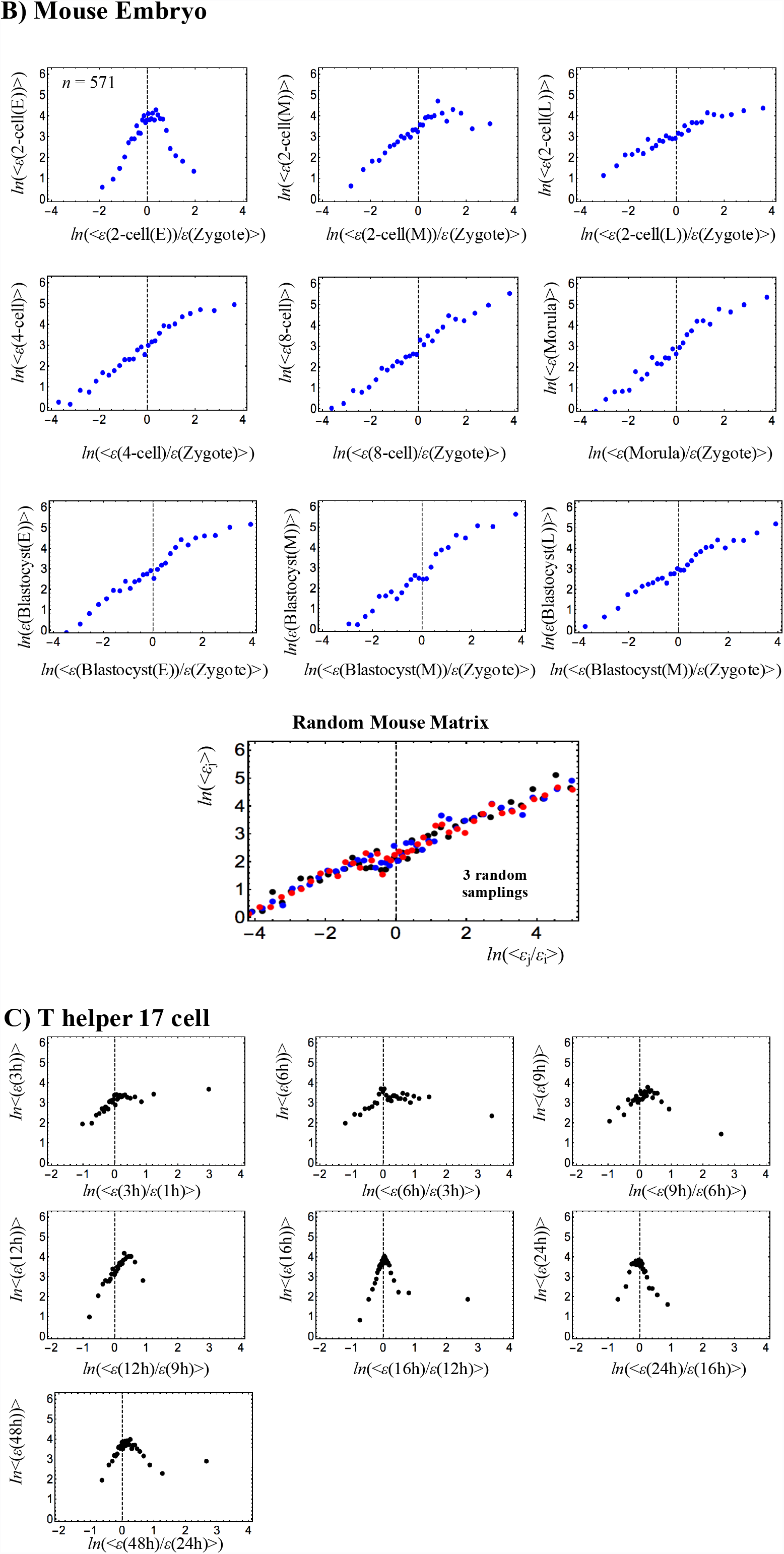
*Complete sandpile transitional analyses of RNA-Seq data (RPKM) for A) human and B) mouse embryo development from the zygote stage, and C) T helper 17 cell differentiation from naïve CD4+ T cells. Random human and mouse expression matrixes (A and B: last panels), (i,j) (i: number of time points; j: number of RNAs; right panels in the second row) reflect stochastic overall expression (i.e., zero correlation between any time points); random matrixes are generated by random shuffling of the corresponding original expression matrixes to show linear correlation behavior (3 random samplings are shown in different colors), similar to that of the DMSO random expression matrix (microarray data: Figure 3D). The embryo results suggest that reprogramming of the genome destroys the SOC zygote control in early embryo development. In T helper 17 cell (Th17) differentiation, the development of a sandpile-type critical transition is observed between sequential cell states.*

This result indicates that there is a critical transition at the genome-state change in the ensemble of cells from single-cell stochastic to highly correlated cell-population behavior (i.e., emergent criticality-induced complexity matching [12]) in overall expression - the emergent layer of a relevant collective regulation starting from a given minimal threshold number of cells [11]. The near-stochastic pattern (Figure 7D) of helper Tell 17 cell differentiation at the onset (single cell) confirms such a transition. The elucidation of the statistical mechanism [31] of the emergent layer of collective regulation in a cell population may explain how coherent oscillation of a critical state in the ensemble of stochastic expression (coherent-stochastic oscillation) [10,11] emerges in interacting cell ensembles.

## III. Distinct Time-Averaged Critical States in Terminal Cell Fates

The self-similarity around the CP (Figure 6) to overall expression highlights three distinct distribution patterns of gene expression relative to different critical states:

i. A unimodal profile corresponding to high-variance expression for a super-critical state, which belongs to a flexible genomic compartment for dominant molecular transcriptional activity.
ii. A flattened unimodal profile (intermediate-variance expression) for a near-critical state, corresponding to an equilibrated genomic compartment, where the critical transition emerges.
iii. A bimodal profile for HRG and EGF responses in MCF-7 cells or a unimodal profile for the DMSO and atRA responses in HL-60 cells for a sub-critical state (low-variance expression). The sub-critical state is the compartment where the ensemble behavior of the genomic DNA structural phase transitions is expected to play a dominant role in the expression dynamics. In the HRG response, a subset of consecutive genes pertaining to the same critical state (called barcode genes) on chromosomes, spanning from kbp to Mbp, has been shown to be a suitable material basis for the coordination of phase-transitional behaviors (refer to Figure 8A in [11]).

The presence of different distributions of a sub-critical state points to different forms of SOC in biological processes with a varying sub-critical state, with regard to the bimodal character of the corresponding expression profile.

In the next section, we will show the existence of the expression flux flow between critical states, which induces temporal fluctuation of the critical point as shown in Figure 4. For evidence of such flow, we need to focus on the average critical state, and then show how perturbation from this average generates activation/inactivation fluxes across the critical states.

The self-similarity of the symmetry break around the CP suggests that critical states have distinct profiles. The degree of *nrmsf* acts as the order parameter for distinct critical states in mRNA expression [10,11]. Thus, to develop a sensible mean-field approach, we estimate bimodality coefficients along *nrmsf* by the following steps (Figure 9):

i. Sort and group the whole mRNA expression according to the degree of *nrmsf.* The *nrmsf* grouping is made at a given sequence of discrete values of *nrmsf*, and
ii. Evaluate the corresponding temporal average of the bimodality coefficient over time to examine if the mean field (behavior of averages of groups) shows any distinctive behavior to distinguish critical states.

**Figure 9:**
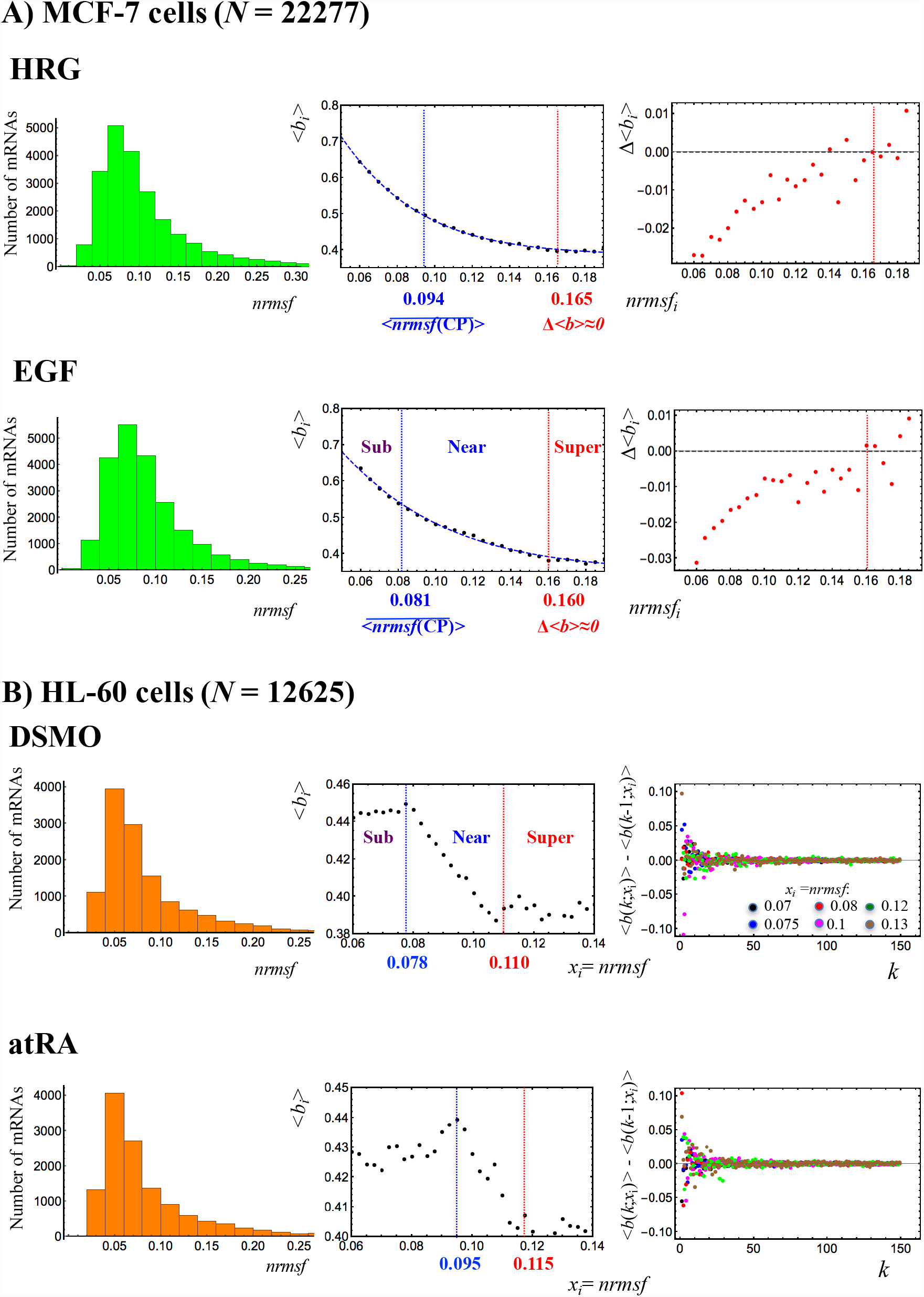
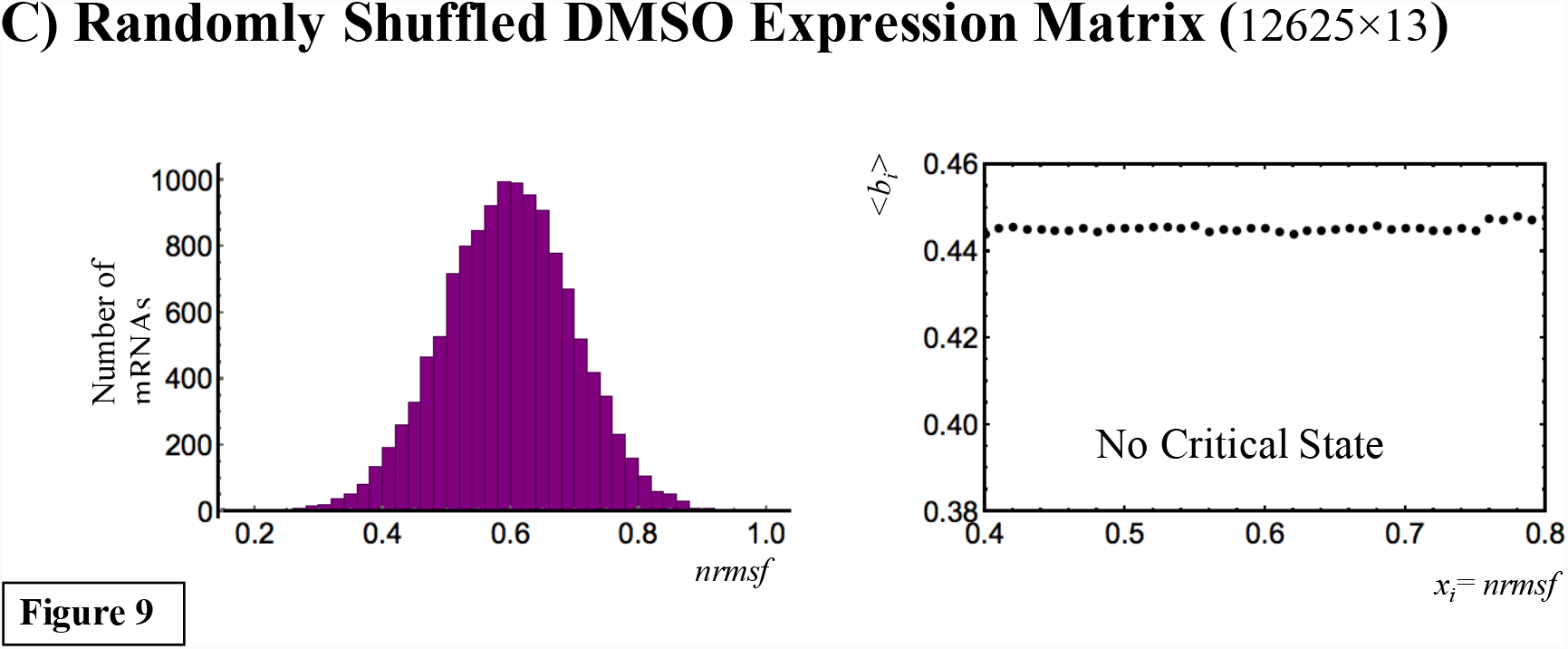
Critical states revealed through distinct functional behaviors of the bimodality coefficient: *A) MCF-7 cells, B) HL-60 cells and C) Random DMSO expression matrix (see Figure 3D). The left panels show the frequency distribution of expression according to the degree of nrmsf. The center panels show the temporal average of Sarle’s bimodality coefficient, <b*_*i*_*>, of the i*^*th*^ *group over time; the value 5/9 represents the threshold between the unimodal (below 5/9) and bimodal or multimodal distributions (above 5/9). The grouping of expression is made at a specific sequence of discrete values of nrmsf (x*_*i*_: *nrmsf*_*i*_ *= i/100; i: integers) with a fixed range: x*_*i*_ - *kd < x*_*i*_ *< x*_*i*_ *+ kd. The values of k and d are set to be k= 150 and d = 0.0001 for MCF-7 and HL-60, and k = 100 and d = 0.001 for a random matrix based on the convergence of the bimodality coefficient. The convergence of the difference in bimodality coefficients at x_i_ with an increase in k (i.e., as the number of elements in a group increases) is shown between the next neighbors, <b*_*i*_*(k;x*_*i*_*)> - <b*_*i*_*(k-1;x*_*i*_*)> for HL- 60 cells in the right panels of B). The 6 colored dots represent the convergent behaviors of different nrmsf points. The behavior of the time average of the bimodality coefficient exhibits* *A) Tangent hyperbolic functions, b*_*i*_ *= a-tanh(b + c*〈*nrmsf*〉_*i*_*); a = 1.38 and 1.35; b = 0.123 and 0.301; c = 13.8 and 10.2 for HRG (p< 10*^*−4*^*) and EGF (p< 10*^*−10*^*), respectively,* *B) Heaviside step function-like transitions for HL-60 cell fates, and* *C) No transition for a random DMSO expression matrix, which importantly reveals that random noises through the formation of a Gaussian distribution destroy a sandpile critical behavior.* *Based on these distinct behaviors, we can determine the boundaries of averaged critical states (Table 1; see **section III**):* *A) Critical states are defined by two points: the average CP,* 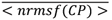, *the onset of a genome avalanche (Figure 4A), for the upper boundary of the sub-critical state, and the point where the change in the bimodality coefficient, Δ<b*_*i*_*>, reaches zero for the lower boundary of the super-critical state; the near-critical state is between them, and* *B) Step function-like transitions reveal the boundaries of averaged critical states, where the near-critical state corresponding to the transitional region separates the other states.*

Figure 9 clearly reveals that, in the overall expression, the mean-field behavior of the average bimodality coefficient confirms the unimodal-bimodal transition of MCF-7 cells and the unimodal-unimodal transition of HL-60 cells. Notably, the mean-field behaviors of bimodality coefficients follow tangent hyperbolic functions (Figure 9A) for MCF-7 cells and Heaviside-like step functions for HL-60 cells (Figure 9B). The distinct time-average behaviors between different cell types further support cell type-specific SOC control.

In contrast, a randomly shuffled expression matrix (DMSO: HL-60 cells) does not exhibit any apparent transitional behavior (Figure 9C), which further confirms the existence of distinctive averaged critical states in both MCF-7 and HL-60 cells.

Two different behaviors in terms of the bimodality coefficient (see Table 1) are evident:

a) In MCF-7 cells, the averaged CP corresponds to the onset of scaling-divergent behavior and the unimodal-bimodal symmetry breaking of the expression profile (Figure 6A), so that the averaged sub-critical state (bimodal profile) is below the averaged CP (*nrmsf*< 0.094 for HRG and *nrmsf*< 0.081 for EGF; Figure 4). The super-critical state (unimodal profile) is above the point where the change in the bimodality coefficient reaches zero (*nrmsf*> 0.165 for HRG and *nrmsf*> 0.160 for EGF), and the near-critical state (flattened unimodal) is between them (Figure 9A).

b) In HL-60 cells, the transition of bimodality coefficients clearly distinguishes average critical states (Figure 9B). Notably, the averaged CP does not correspond to the onset of scaling-divergent behavior for either response (Figure 4B). In fact, the onset is extended: in the DMSO response, the scaling region extends to the upper boundary of the near-critical state, while in the atRA response, the scaling region extends to the upper boundary of the sub-critical state, where the averaged CP exists in the (averaged) sub-critical state. This is attributable to the collapse of autonomous bistable switch (ABS) of the sub-critical state, as discussed in the previous section. In summary, the mean-field behavior of bimodality coefficients exhibits markedly different behaviors that can be used to distinguish averaged critical states.

**Table 1:** Averaged Critical States

| Averaged Critical States | MCF-7 cells |  | HL-60 cells |  |
| --- | --- | --- | --- | --- |
|  | N = 22277 |  | N = 12625 |  |
|  | HRG | EGF | DMSO | atRA |
| **Super-critical** | $0.165 < nrmsf$ | $0.160 < nrmsf$ | $0.110 < nrmsf$ | $0.115 < nrmsf$ |
|  | 3051 mRNAs | 1969 mRNAs | 2582 mRNAs | 2465 mRNAs |
| **Near-critical** | $0.094 < nrmsf < 0.165$ | $0.081 < nrmsf < 0.160$ | $0.078 < nrmsf < 0.110$ | $0.095 < nrmsf < 0.115$ |
|  | 6814 mRNAs | 9119 mRNAs | 2226 mRNAs | 995 mRNAs |
| **Sub-critical** | $nrmsf < 0.094$ | $nrmsf < 0.081$ | $nrmsf < 0.078$ | $nrmsf < 0.095$ |
|  | 12412 mRNAs | 11189 mRNAs | 7817 mRNAs | 9205 mRNAs |

## IV. Coherent-Stochastic Behavior (CSB) in Critical States

Critical states display coherent-stochastic behavior (CSB), where coherent behavior emerges in ensembles of stochastic expression [11]. In Figures 10 A,B, random sampling of the averaged critical states for both MCF-7 and HL-60 cells clearly shows that

1) The near-zero Pearson correlation between different randomly selected gene ensembles in the critical states reveals stochastic expression, and

2) There is a sharp damping in variability (Euclidean distance of single time points from the center of mass CM(*t*_j_) of the critical states). This is a further confirmation that the CM(*t*_j_) of the critical states represents their coherent dynamics.

**Figure 10:**
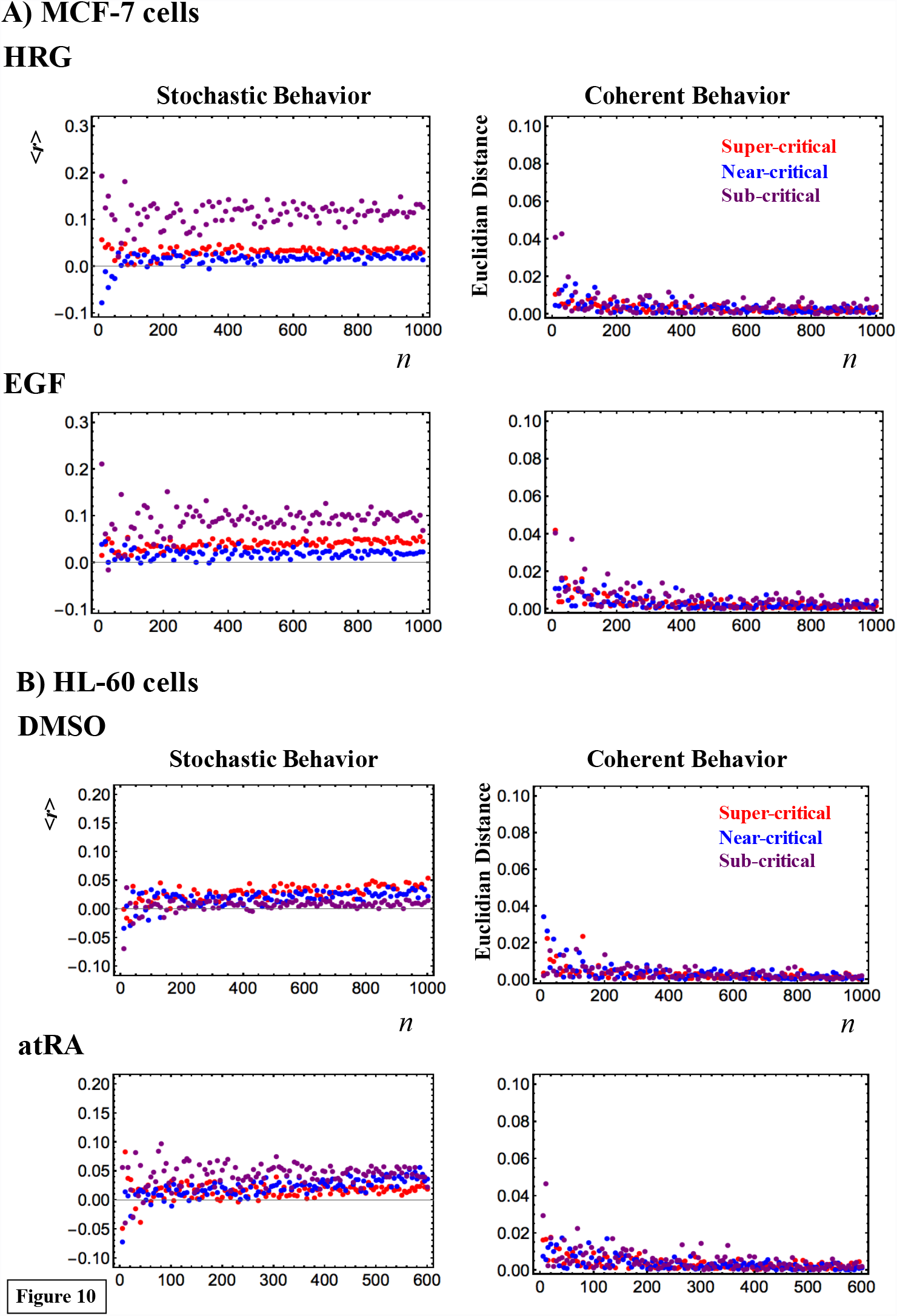
Coherent-stochastic behaviors in critical states: *A) MCF-7 cells and B) HL-60 cells. 200 random-number ensemble sets are created, where each set has n (variable) sorted numbers, which are randomly selected from an integer series {1,2,..,N} (N: total number of mRNAs in a critical state: Table 2). These random sets are used to create random gene ensembles from each critical state.* *1) Left panels (A and B): For each n, Pearson correlations are evaluated between different random gene ensembles in the critical states by averaging over the 200 ensemble sets and experimental time points (**Methods**). This gives a near-zero Pearson correlation, consistent with the global stochastic character of microscopic transcriptional expression regulation in critical states.* *2) Right panels (A and B): For each random gene ensemble, the Euclidean distance of single time points from the center of mass CM(t*_*j*_*) of the critical states is evaluated by averaging over the 200 ensemble sets. The sharp damping of variability confirms that the emergent coherent dynamics of the critical states correspond to the dynamics of the CM(t*_*j*_).

The emergent CSB of critical states through SOC control of the entire expression shows how a population of cells can overcome the problem of stochastic fluctuation in local gene-by-gene regulation. Moreover, the fact that CM represents the coherent dynamics of CSB tells us how macroscopic control can be tuned by just a few hidden parameters through SOC. This collective behavior emerges from intermingled processes involving the expression of more than 20,000 genes; this corresponds to the notion that, despite their apparent bewildering complexity, cell state/fate changes collapse to a few control parameters, and this ‘sloppiness’ [9] derives from a low effective dimensionality in the control parameter-space emerging from the coherent behavior of microscopic-level elements.

Furthermore, mRNA expression in a microarray reflects populations of millions of cells, so that the ensemble of expression at *t = t*_j_ represents a snapshot of time-dependent thermodynamic processes far from equilibrium, where the usual pillars of equilibrium thermodynamics such as ‘time-reversibility’ and ‘detailed balance’ break down, to reveal the characteristics of a dissipative or far-from equilibrium system.

## V. Sub-Critical State as a Generator of Self-Organizing Global Gene Regulation

The temporal development of critical transitions in overall expression suggests that molecular stressors on MCF-7 and HL-60 cells induce the perturbation of self-organization through interactions between critical states. The emergent coherent-stochastic behavior (CSB) corresponds to the dynamics of the center of mass (CM) of critical states (refer to Figure 6B in [11], which shows ON-OFF coherent oscillation of the sub-critical state with its CM). Hence, an understanding of the dynamics of the CM of critical states and their mutual interactions should provide insight into how the perturbation of self-organization in whole-mRNA expression evolves dynamically through perturbation.

Here, it would be useful to abstract the essence of the dynamics of critical states and their mutual interactions into a simple one-dimensional CM dynamical system: the CM of a critical state, *X*(*t*_j_), is a scalar point, and thus, the dynamics of *X*(*t*_j_) can be described in terms of the change in the one-dimensional effective force acting on the CM. From a thermodynamic point of view, this force produces work, and thus causes a change in the internal energy of critical states. Hence, we investigate the genome as an open thermodynamic system. The genome is considered to be surrounded by the intranuclear environment, where the expression flux represents the exchange of genetic energy or activity. This picture shows self-organized overall expression under environmental dynamic perturbations; the regulation of mRNA expression is managed through the mutual interaction between critical states and the external connection with the cell nucleus milieu.

To quantitatively designate such flux flow, we set up the effective force acting on the CM, *f*(*X*(*t*_j_)) at *t* = *t*_j_, where the expression of each gene is assigned to have an equal constant mass (set to unity). The impulse, *F*Δ*t*, corresponds to the change in momentum Δ*P* and is proportional to the change in average velocity: *v*(*t*_j+1_) - *v*(*t*_j_). Since a consideration of the center of mass normalizes the number of genes being expressed in a critical state, we set the proportionality constant, i.e., the mass of the CM, to be unity. Thus, *f*(*X*(*t*_j_)) = –*F =* (*v*(*t*_j_) - *v*(*t*_j+1_))/Δ*t*, where Δ*t* = *t*_j+1_ - *t*_j-1_, and the force is given a negative sign, such that a linear term in the nonlinear dynamics of *X*(*t*_j_) corresponds to a force under a harmonic potential energy. The effective force, *f*(*X*(*t*_j_)) can be decomposed into IN flux: incoming expression flux from the past *t* = *t*_j-1_ to the present *t* = *t*_j_, and OUT flux: outgoing expression flux from the present *t* = *t*_j_ to the future *t* = *t*_j+1_ (Δ*t*_j_ = *t*_j_ *- t*_j-1_):

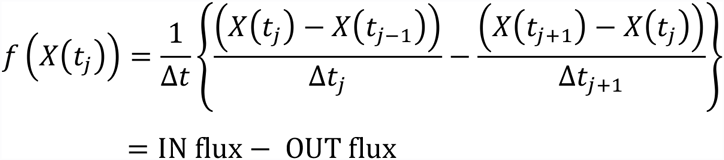

We call the force, *f*(*X*(*t*_j_)), the *net self-flux of a critical state*. The net self-flux, IN flux - OUT flux, has a positive sign for incoming force (net IN self-flux) and a negative sign for outgoing force (net OUT self-flux).

When we adopt this concept of expression flux, it becomes straightforward to define the *interaction flux* of a critical state *X*(*t*_j_) with respect to another critical state or the environment *Y*:

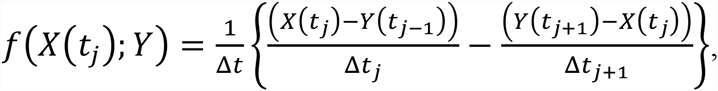

where, again, the first and second terms represent IN flux and OUT flux, respectively, and the net, IN flux- OUT flux, represents incoming (IN) interaction flux from *Y* for a positive sign and outgoing (OUT) interaction flux to *Y* for a negative sign. As noted, the interaction flux between critical states can be defined as when the number of gene expressions in a critical state is normalized, i.e., when we consider the CM. Due to the law of force, the net self-flux of a critical state is the summation of the interaction fluxes with other critical states and the environment (see **Methods**).

Next, we consider how the net IN (OUT) flux of a critical state, the effective force acting on the CM of a critical state, corresponds to the dynamics of its CM. Figure 11 clearly shows that the trend of the dynamics of the CM of a critical state follows its net self-flux dynamics, in that the CM is up- (down-) regulated for net IN (OUT) flux, where the CM is measured from its temporal average value. This implies that the respective temporal average values are the baselines for both flux and CM; this is further confirmed by the existence of an average flux balance in critical states, where the net average fluxes coming in and going out at each critical state are balanced (near-zero) (**Methods**).

**Figure 11:**
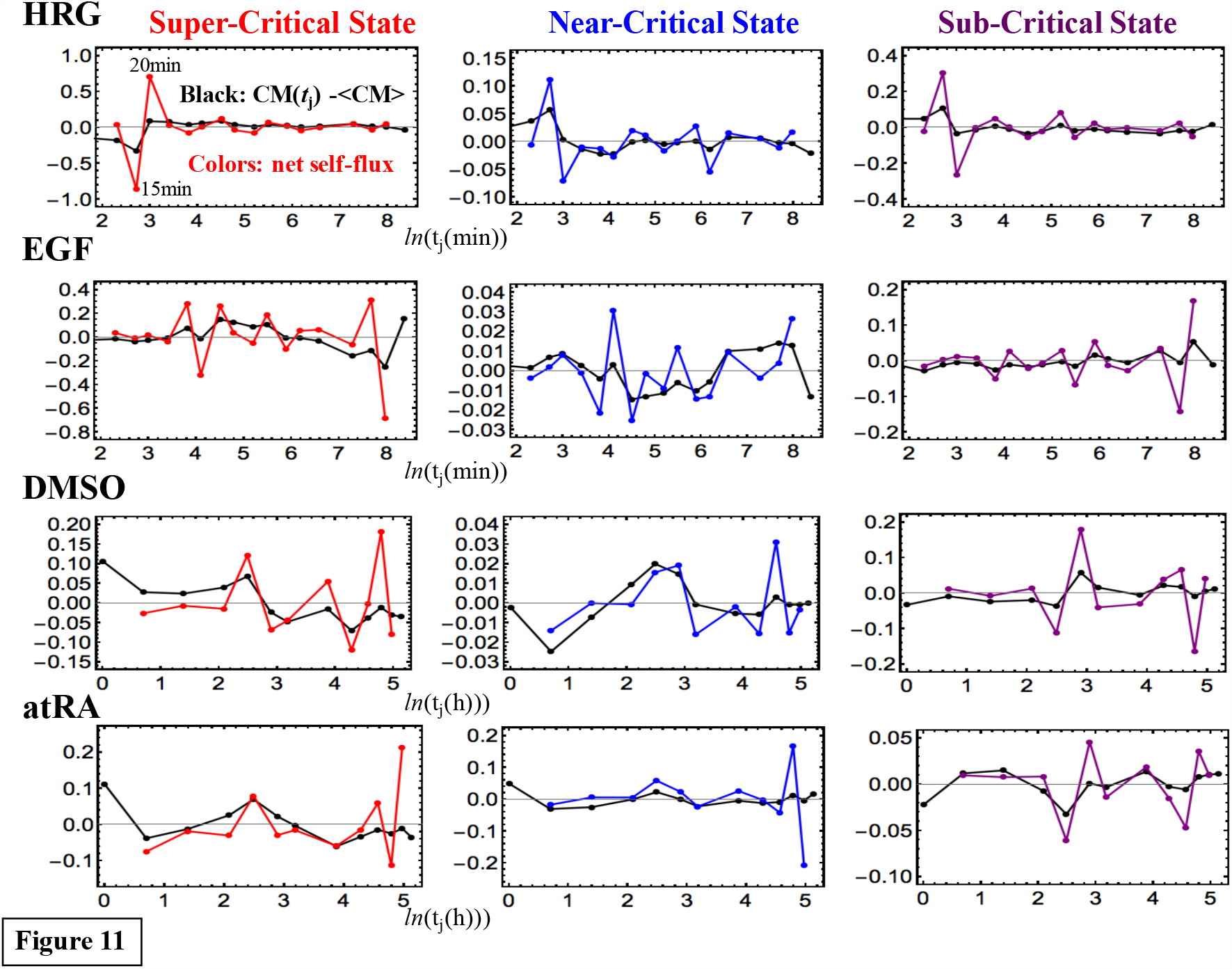
Dynamics of the center of mass (CM) of critical states compared with the net self-flux dynamics: *Colored lines (red: super-critical; blue: near-critical; purple: sub-critical state) represent net self-fluxes of critical states from their temporal averages (effective force acting on the CM: **Methods**). Black lines represent the dynamics of the CM of critical states from their temporal averages, which are increased three-fold for comparison to the corresponding net self-fluxes. The plots show that the net self-flux dynamics follow up- (down-) regulated CM dynamics, such that the sign of the net self-flux (i.e., IN and OUT) corresponds to activation (up-regulated flux) for positive responses and inactivation (down-regulated flux) for negative responses. The natural log of the experimental time points (MCF-7: minutes and HL-60: hours) is shown.*

Thus, we consider both temporal averaged expression flux among critical states and the fluctuation of expression flux from the average (*flux dynamics*; **Methods**), so that

*- Averaged expression flux shows a temporal average expression flow among critical states through the environment, i.e., the characteristics of an open thermodynamic genomic system* (“genome engine” in **Discussion**), and

*- The flux dynamics represent fluctuation of the expression flux flow that is markedly different from the basic properties in equilibrium Brownian behavior under a detailed balance*. Furthermore, the sign of the net self-flux (i.e., IN or OUT) corresponds to the activation (up-regulation) of flux for positive responses and inactivation (down-regulation) for negative responses.

We examined the average flux network for the processes of MCF-7 and HL-60 cells. Figure 12A and Table 2 intriguingly reveal four distinct processes that share common features in their flux networks:

**Figure 12:**
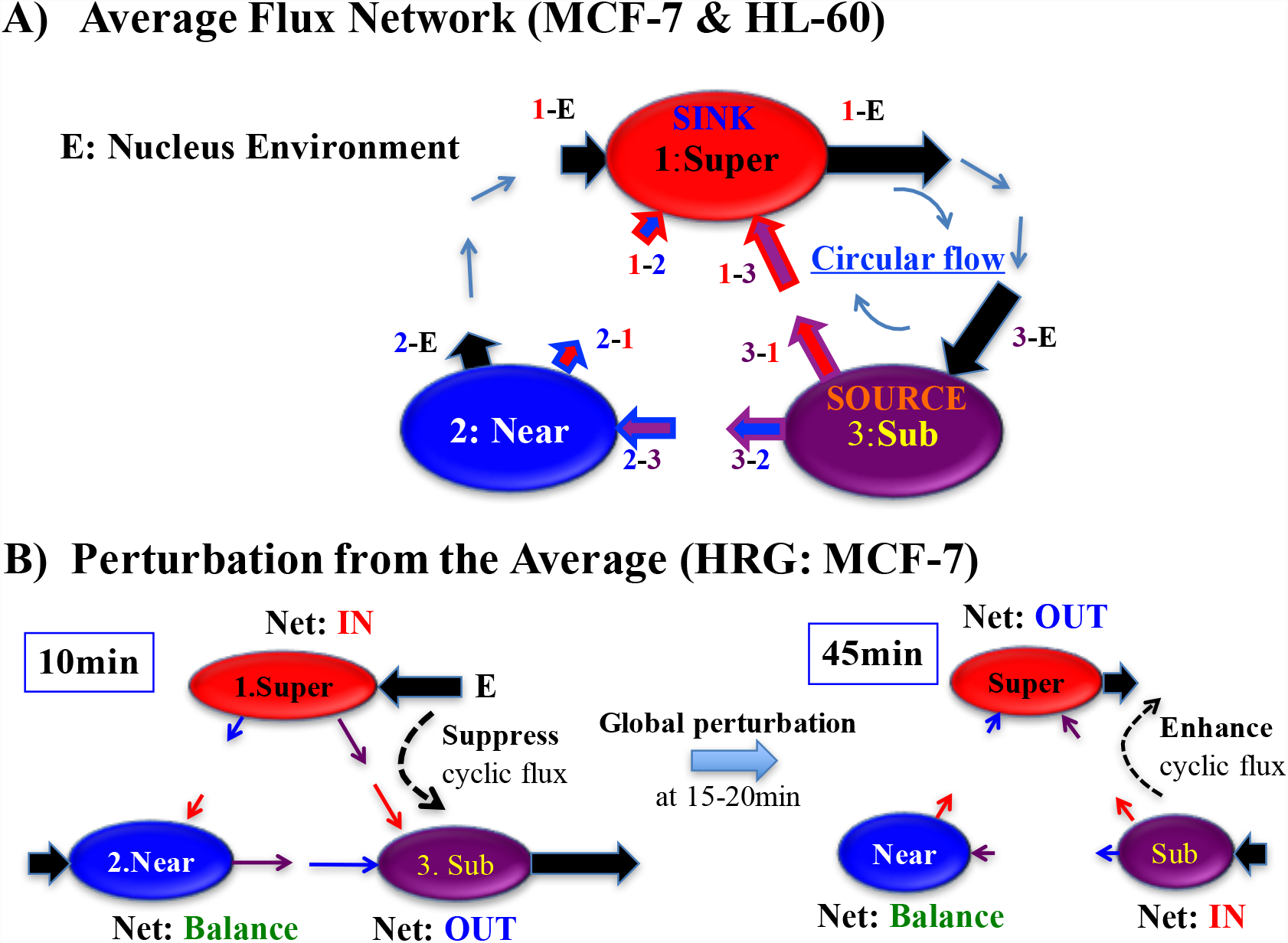
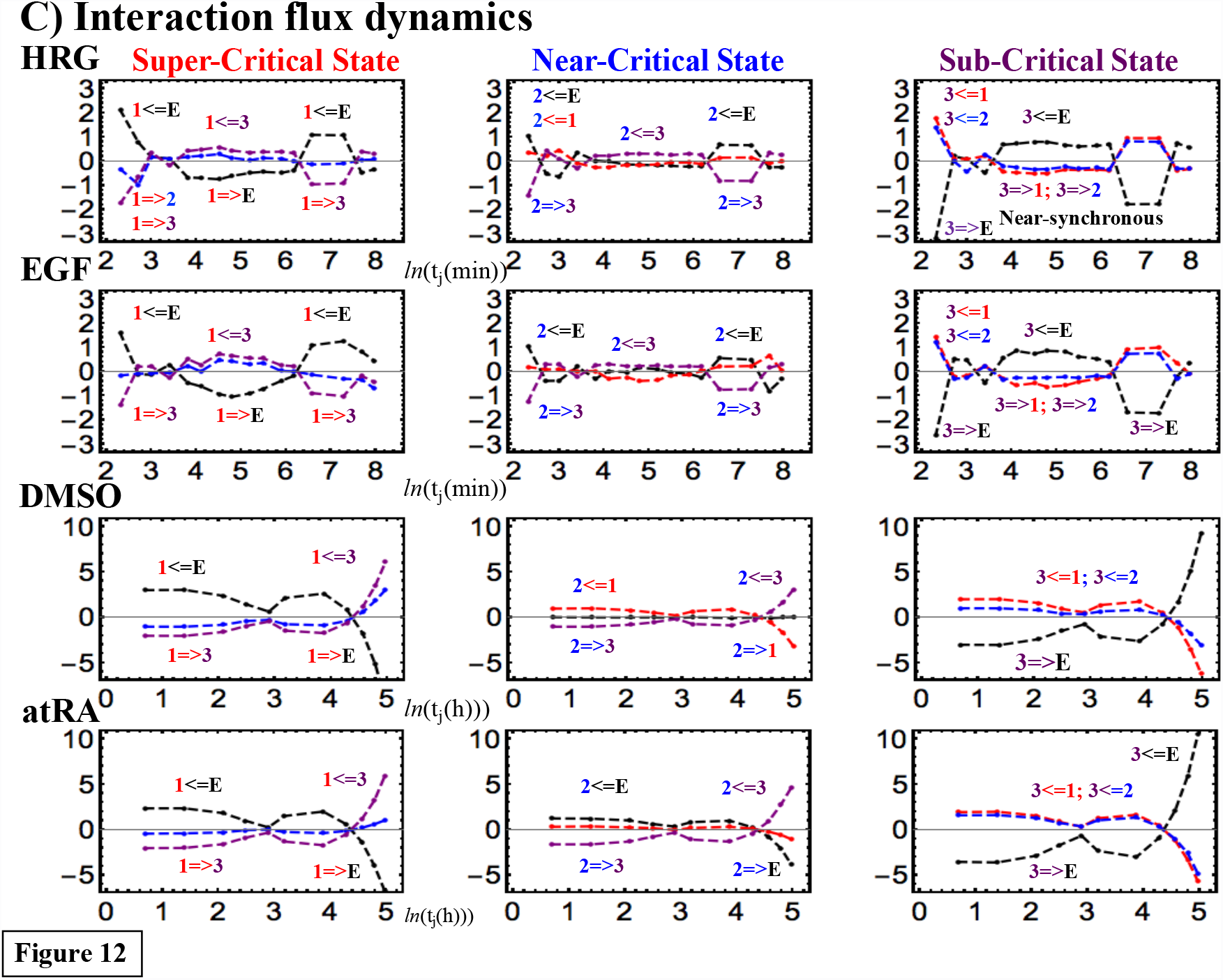
Genomic expression dynamics revealed through a flux analysis that includes crosstalk with the environment: *A) Average values of interaction flux (colored arrows) for MCF-7 and HL-60 cells show that a sub-critical state acts as an internal ‘source’, where IN flux from the environment is distributed to other critical states. In contrast, a super-critical state acts as an internal ‘sink’ that receives IN fluxes from other critical states, and the same amount of expression flux is sent to the environment, due to the average flux balance (**Methods**). Furthermore, the formation of a dominant cyclic state flux is revealed between super- and sub-critical states through the environment. The average interaction flux is represented as i-j: interaction flux of the i*^*th*^ *critical state with the j*^*th*^ *critical state (i, j= 1: super- (Super; red), 2: near- (Near; blue), 3: sub-critical state (Sub; purple)), and a colored arrow for an internal i-j interaction, where outline and base colors are based on the i*^*th*^ *critical and j*^*th*^ *critical states, respectively, points in the direction of interaction with the relative amount of flux (see details in Table 2; positive and negative values represent incoming and outgoing flux, respectively, at a critical state). E represents the internal nucleus environment.* *B) An early flux dynamics event in the HRG response resulting from the HRG interaction flux dynamics (see C) are shown. Interaction flux dynamics i<=j (or i=>j; color based on j) represent the interaction flux from the j*^*th*^ *critical state to the i*^*th*^ *critical state or vice versa. The interaction fluxes (see HRG in C; the flux direction changes at y =0) align to suppress the cyclic state flux at 10min (the first point in C), where the interaction flux shows 1<=E, 1=>2, and 1=>3 at the super-critical state, 2<=E, 2<= 1, 2=>3 at the near-critical state, and 3=>E, 3<=1, 3<=2 at the sub-critical state. They then align to enhance the cyclic state flux at 45 min (5^th^ point in C). This change in the dynamic flux structure is due to the global perturbation at 15-20min (see Figure 14; **section VI**). At each node, the net flux (Figure 11; 10min: first point; 45min: 5th point) is indicated as - IN for net incoming flux (y >0), OUT for outgoing flux (y <0), or Balance (y~0). Note: The average flux balance at each node is maintained, but not at individual time points.* *C) Notably, for MCF-7 cell fates (HRG and EGF), near-synchronous interaction flux dynamics are seen at sub-critical states. The plots further show that the overall patterns are similar between the same cell types, which again supports cell-type-specific SOC control.*

**Table 2:** **Average Interaction Fluxes:** The Arrow shows the direction of flux.

|  | MCF-7<br>HRG | MCF-7<br>EGF | HL-60<br>DMSO | HL-60<br>atRA |
| --- | --- | --- | --- | --- |
| **Super-Critical**<br>**SINK** |  |  |  |  |
| **1-2** Super<br>←<br>Near | 0.3 | 0.2 | 1.3 | 0.5 |
| **1-3** Super<br>←<br>Sub | 1.9 | 1.6 | 2.6 | 2.5 |
| **1-E** Super<br>→<br>EN | -2.2:<br>1.3(Near)<br>-3.5(Sub) | -1.8:<br>1.2<br>-3.0 | -3.9:<br>0.0<br>3.9 | -3.0:<br>1.5<br>-4.5 |
| **Near-Critical** |  |  |  |  |
| **2-1** Near<br>→<br>Super | -0.3 | -0.2 | -1.3 | -0.5 |
| **2-3** Near<br>←<br>Sub | 1.6 | 1.4 | 1.3 | 2.0 |
| **2-E** Near<br>→<br>EN | -1.3 | -1.2 | 0.0 | -1.5 |
| **Sub-Critical**<br>**SOURCE** |  |  |  |  |
| **3-1** Sub<br>→<br>Super | -1.9 | -1.6 | -2.6 | -2.5 |
| **3-2** Sub<br>→<br>Near | -1.6 | -1.4 | -1.3 | -2.0 |
| **3-E** Sub<br>←<br>EN | 3.5 | 3.0 | 3.9 | 4.5 |

### i A dominant cyclic flux between super- and sub-critical states

Average net IN and OUT flux flows reveal how the internal interaction of critical states and external interaction with the cell nucleus environment interact: notably, the formation of robust average cyclic state-flux between super- and sub-critical states through the environment forms a dominant flux flow in the genomic system. This formation of the cyclic flux causes strong coupling between the super- and sub-critical states as revealed through the correlation analysis [11].

### ii Sub-critical state as a source of internal fluxes through a dominant cyclic flux

The direction of the average flux in super- and sub-critical states reveals that the sub-critical state is the source of average flux, which distributes flux from the nucleus environment to the rest of the critical states, where the super-critical state is the sink to receive fluxes from the near- and sub-critical states. Importantly, this clearly shows that the sub-critical state, an ensemble of low-variance expression acts as a generator of perturbation in genome-wide self-organization.

In summary, the results regarding average flux reveal the roles of critical states in SOC control of overall expression, and provide a statistical mechanics picture of how global self-organization emerges through the interaction among critical states and the cell nucleus microenvironment.

## VI. SOC Control Mechanism of Overall Gene Expression

The flux dynamics at critical states reveal the dynamic mechanism of SOC control in overall expression. We can elucidate how dynamic interaction among critical states and the environment perturbs the average flux flow in the genomic system, as follows:

### 1 Flux dynamics reveal early nucleus activities

In Figures 12B,C, the flux dynamics of CMs in HRG-stimulated MCF-7 cells show how the average flux network (Figure 12A) is substantially perturbed at early time points, which reflects the early occurrence of significant genetic energy dissipation. This can be seen from the net self-flux dynamics (HRG response; Figure 11). At 15-20min, *global perturbation* involving a large change in net self-flux in more than one critical state occurs: the net self-flux of the super-critical state shows a pulse-like change from net OUT (negative value) to IN self-flux (positive), i.e., an increase in internal energy, which explains how the significant net flux into the super-critical state is used to activate the expression of genes in the super-critical state from 15 to 20min. In contrast, for the sub- and near-critical states, the net self-flux significantly changes from the net IN to net OUT self-flux (anti-phase with respect to the dynamics of the super-critical state), i.e., they show a loss of internal energy.

The global perturbation at 15-20 min stems from genetic energy flow in the genomic system, which shows a change from the strong suppression of cyclic flux at 10min before perturbation to the enhancement at 45min after perturbation (Figure 12B): at 10min, the flow of the interaction fluxes between the super- and sub-critical states aligns against the average cyclic flux to suppress the cyclic flux (the strongest inhibition over time), and then, at 45 min, the interaction fluxes change and align to enhance the cyclic flux; the change in the cyclic flux is due to reversal of the genetic energy flow at 10min by a pulse-like global perturbation in self-organization at 15-20 min (see below).

These results suggest the presence of early cell nucleus activities in HRG-stimulated MCF-7 cells. At 15min, genetic information, through signaling activities [28,29] in the cytoplasm from the cell membrane induced by HRG, reaches the nucleus, and at 15-20min it activates the high-variance genes of the super-critical state. In contrast, the near- and sub-critical states (intermediate- and low-variance genes, respectively) are suppressed, so that the internal genetic energy flow into the environment should induce a change in the physical plasticity of chromatin structures of genes in these states, i.e., less pliable structures at the ensemble scale (not at an individual scale).

### 2 SOC control mechanism of the genome-state change - role of the critical gene ensemble

Here, we describe the *SOC control mechanism of the genome-state change* in terminal cell fates at the cell population level. The genome-state change occurs through the breakdown of initial-state criticality as follows:

Expression flux dynamics (Figure 13) show that the net self-flux of the sub- and super-critical states can be well-described in terms of the net interaction between sub- and near-critical states (Sub-Near), and between sub- and super-critical states (Sub-Super), respectively. Namely, the dynamic interactions of Sub-Near and Sub-Super determine the net self-flux of the sub- and super-critical states, respectively, which represents the effective forces acting on their CMs, and thus determines their coherent oscillatory dynamics (Figure 11; see Figure 6 in [11]). This essential role of the interactions explains how the temporal change in criticality at the near-critical state, i.e., in expression of the **critical gene ensemble**, directly perturbs the sub-critical state (the generator of flux dynamics) through their mutual interaction, and the perturbation of this generator can spread over the entire system (Figure 13A).

**Figure 13:**
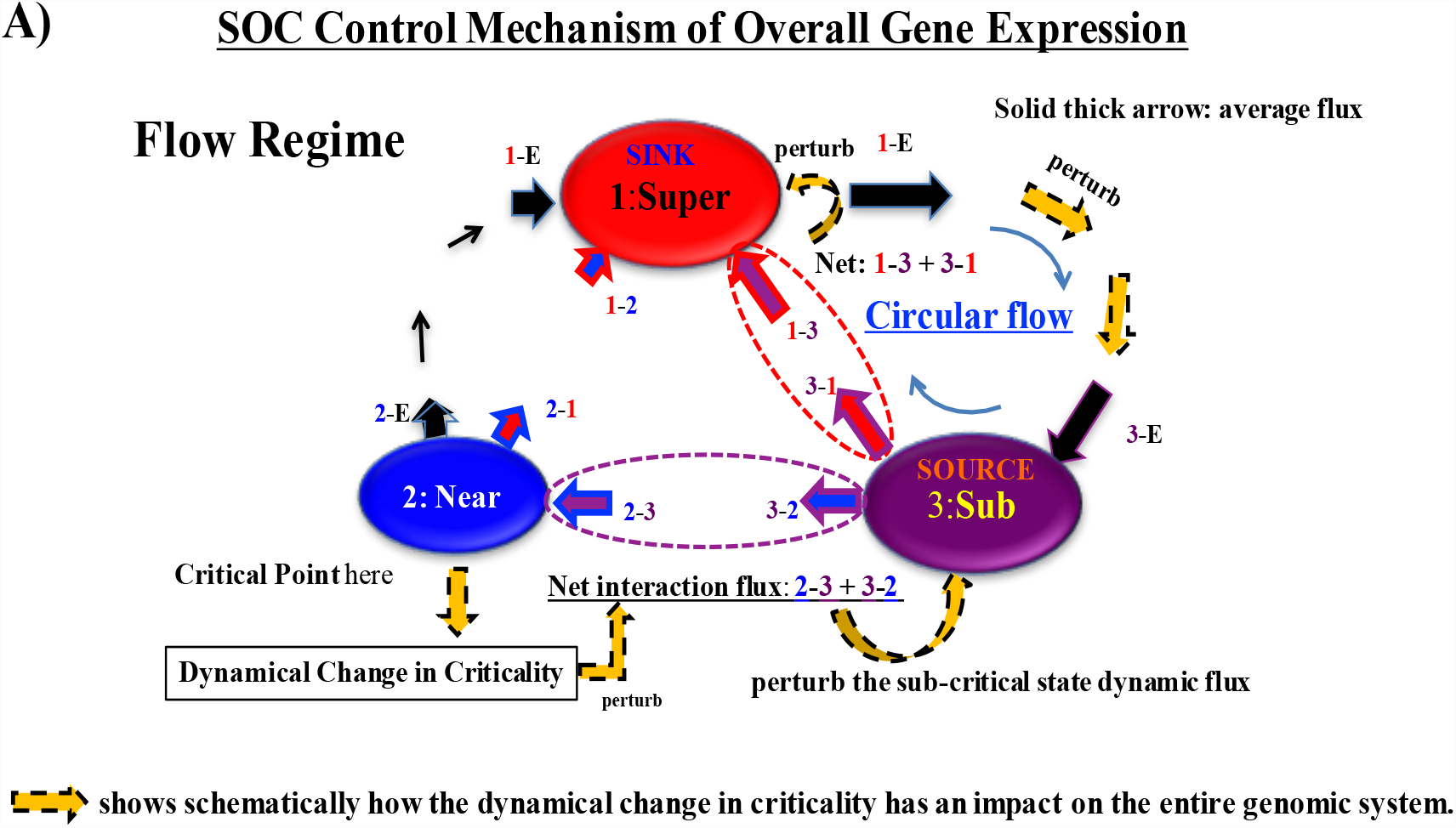
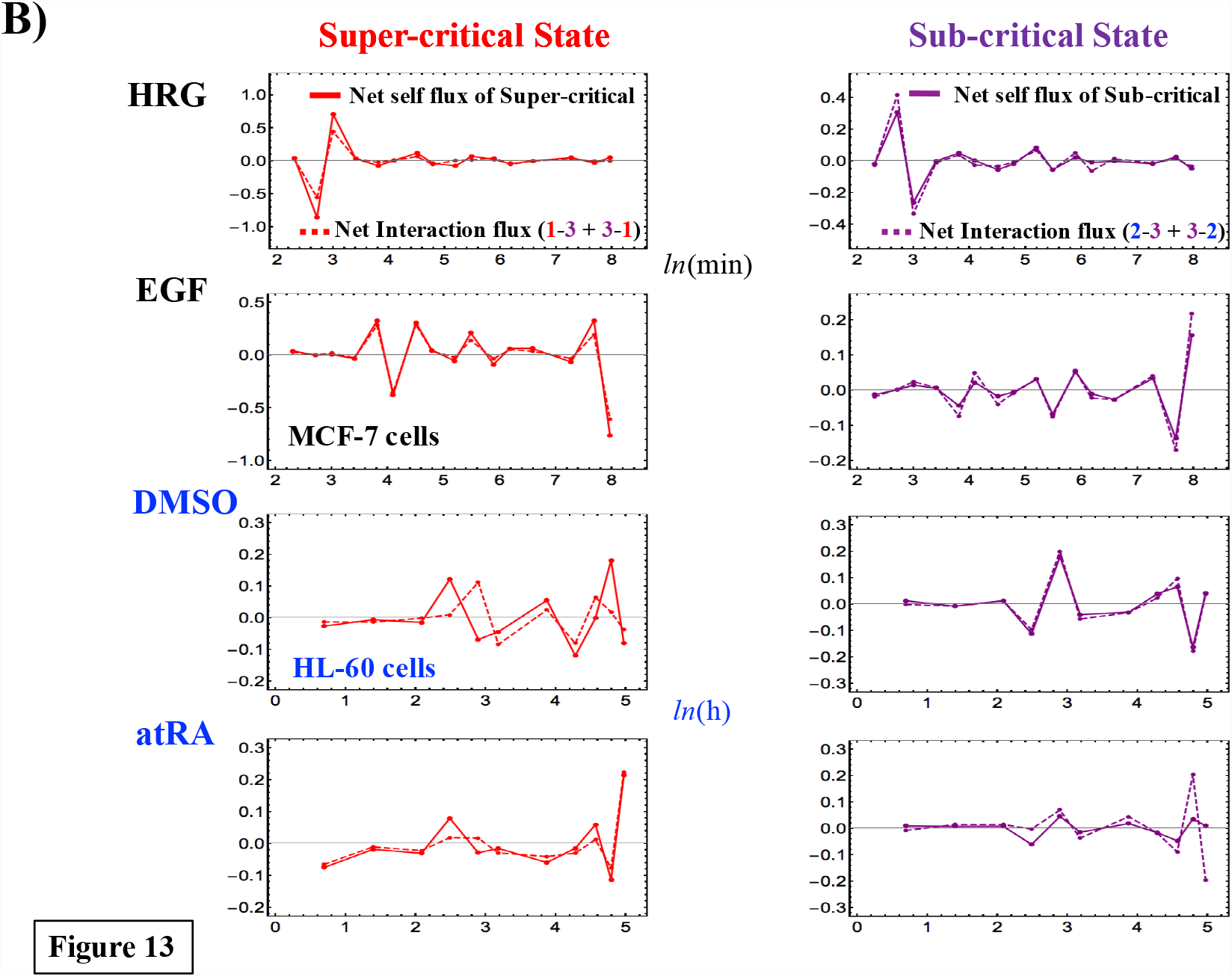
SOC control mechanism of overall gene expression in terminal cell fates (MCF-7 and HL-60 cells): *A) Flow regime shows how a dynamical change in criticality (critical behaviors) at the CP in terminal cell fates affects the entire genome (thick yellow arrow with a black dashed outline) as follows (Super: super-; Near: near-; Sub: sub-critical state):* *The dynamical change in criticality (critical behaviors) that originates from the CP at Near perturbs the net interaction flux (2-3 + 3-2; refer to Figure 12A) at Sub-Near, which determines the net self-flux of Sub (B: right panels), and thus directly perturbs the Sub-response. This induces a perturbation in the net interaction flux of Sub-Super (1-3+3-1), i.e., directly perturbs the Super-response (B: left panels). Furthermore, these perturbations on Sub and Super disturb the dominant cyclic flux between Sub and Super, which in turn has an impact on sustaining the critical dynamics (source: Sub and sink: Super); solid thick arrows represent average fluxes (Table 2).* *Note: the net self-flux of a critical state represents the effective force acting on CM, and thus dynamic interactions among states determine the coherent oscillatory dynamics of Sub and Super (Figure 11). This schematic picture also shows that the erasure of an initial-state criticality at the genome-state change reflects the destruction of the initial-state SOC control mechanism of the dynamical change in the entire genomic system (i.e., pruning of the mechanism for regulating global gene expression at the initial state).*

The genome-state change (cell-fate change in the genome) occurs in such a way that the initial-state SOC control of overall gene expression (i.e., initial-state global gene expression regulation mechanism) is destroyed through the erasure of an initial-state criticality; this shows that the critical gene ensemble of the CP plays a significant role in determining the cell-fate change. In other words, a statistical mechanical layer [31] for the dynamic control of genome-wide expression emerges in cells, where the critical gene ensemble ‘drives’ the fate of cell ensembles.

### 3 Global and local perturbations exist in the SOC control

So far, we have applied the expression flux concept to the effective force acting on the CM of a critical state, *X*(*t*_j_). This concept can also be extended to define kinetic energy self-flux for the CM of a critical state. The kinetic energy of the CM with unit mass at *t* = *t*_*j*_ is defined as 1/2v(*t*_*j*_)^2^, such that the net kinetic energy self-flux from its temporal average can be defined as

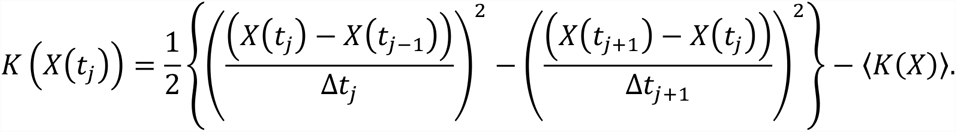

We investigate when a significant kinetic energy self-flux occurs to better understand the global perturbation in the SOC control. Figure 14 shows that global perturbations are more evident in the net kinetic energy self-flux dynamics than in the effective force (Figure 11):

i. MCF-7: Global perturbation involving a change in net kinetic energy flux in critical states occurs only at 10-30min, where a pulse-like transition occurs from outgoing kinetic energy flux to incoming flux at 15-20min. The occurrence of this transition is confirmed by a pulse-like maximal response in Pearson autocorrelation (*P*(*t*_j_;*t*_j+1_) at 15-20min: see Figure 4A in [11]). Dissipation of the kinetic energy is evident in the HRG response, which quickly ends at 30min. In contrast, in the EGF response, only vivid flux activity in the super-critical state is apparent until *t* = 36h, which demonstrates *local perturbation* in the EGF response. Thus, global and local perturbations differentiate the cell fate between the responses to HRG and EGF: the dissipative pulse-like global perturbation in the HRG response at 15-20min leads to the genome-state change at 3h (Figure 5A), whereas the local perturbation in the EGF response does not induce the state change (see **section I**). Furthermore, this global perturbation (see also paragraph 1) above) suggests *the existence of a novel primary signaling transduction mechanism or a biophysical mechanism which can induce the inactivation of gene expression in the near- and sub-critical states (a majority of expression: mostly low-variance gene expression) within a very short time.* This global inactivation mechanism that is associated with more than 10,000 of low-variance mRNAs in a coordinated manner should be causally related to the corresponding changes in the higher-order structure of chromatin (as discussed in the literature from a theoretical perspective [32-34] and a biological perspective [35,36]).
ii. HL-60: In the response to DMSO, a clear pulse-like global perturbation occurs at 12-18h. In the response to atRA, the first significant global perturbation occurs early (2-4h) and a second smaller global perturbation occurs at 12-18h, which can be confirmed by a change in the effective force (Figure 11). They both show distinct dissipative oscillatory behavior of kinetic energy flux dynamics. Again, these global perturbations occur before the genome-state changes (24h for DMSO and 48h for atRA; Figure 5B). As shown in Supplementary Figure S1, a pulse-like global perturbation may be related to process for autonomous terminal differentiation (the first stage of cell-fate determination; see more in subsection (ii) in **Discussion**).

**Figure 14:**
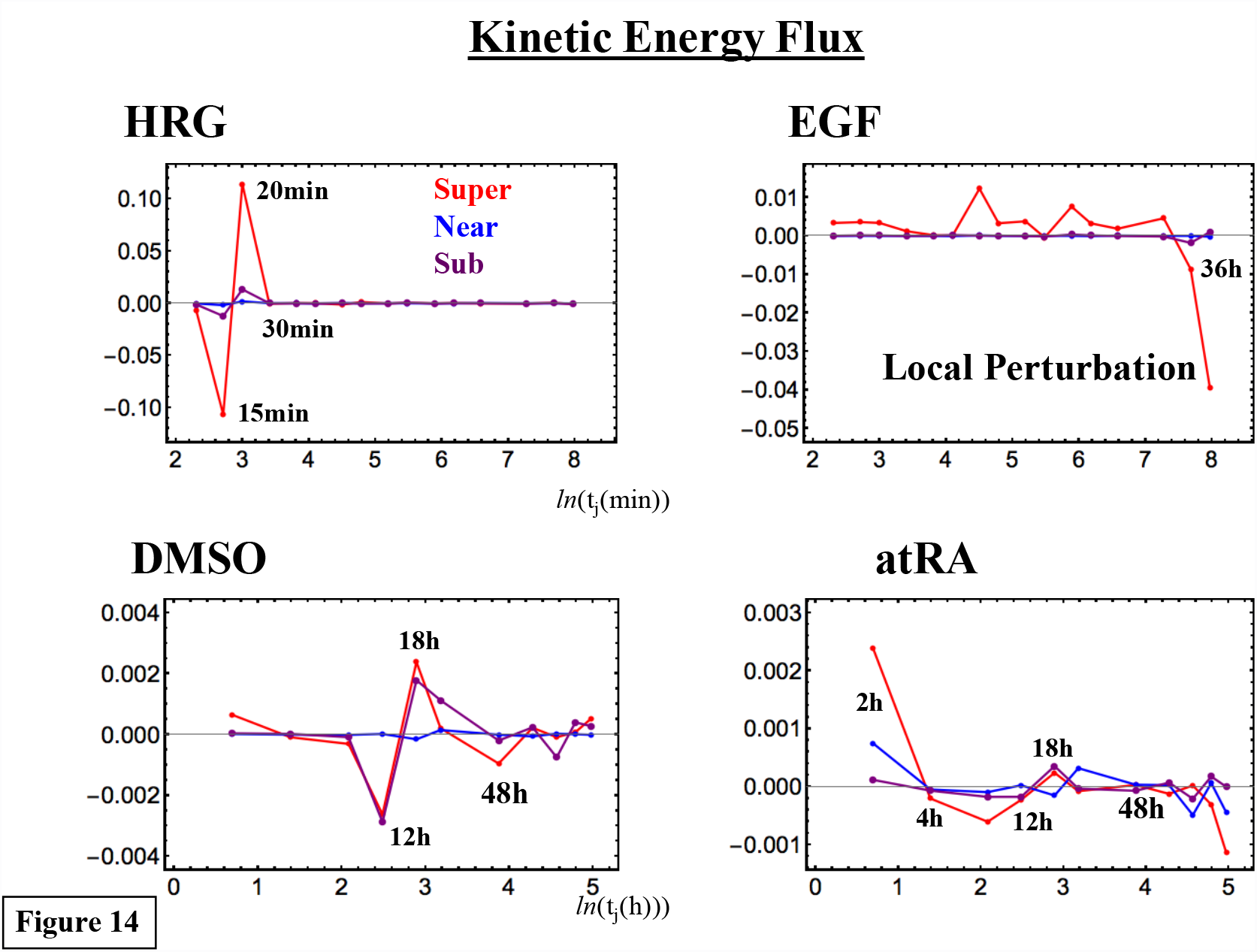
Local and global perturbations in self-organizing genome-wide expression: *The kinetic energy self-flux dynamics (y-axis) for the CM of a critical state (see **section VI**) exhibit clear energy-dissipative behavior. Notably, the results show the occurrence of global and local perturbations of self-organization. Pulse-like global perturbations show a transition from IN to OUT net kinetic energy flux or vice versa (IN-OUT switching) in more than one critical state: at 15-20min, HRG; at 12-18h, DMSO; and at 2-4h (significant) and 12-18h, atRA. In contrast, local perturbation is observed in the EGF response for up to 36h: there is only a marked response in the super-critical state, and almost no response in the other states (i.e., the dynamics of the CM of critical states are localized around their average: Figure 11). The results suggest that the global and local perturbations differentiate MCF-7 cell fates, whereas global perturbations drive the state change in HL-60 cells (see **section I**).*

### 4 Long-term global mRNA oscillation underlies SOC control

The sub-critical state is the source of internal genetic energy flow in SOC control, and therefore, the oscillatory net self-flux of sub-critical states generates a long-term global mRNA oscillation [37] to sustain the self-control of SOC.

## Discussion

We investigated the dynamics of collective gene behavior in several biological processes associated with changes in the cell fate:

1. Early embryo development in human and mouse,
2. Helper T 17 cell differentiation, induction of terminal differentiation in human leukemia HL-60 cells by DMSO and atRA,
3. Activation of ErbB receptors in human breast cancer MCF-7 cells by epidermal growth factor (EGF) and heregulin (HRG).

Our approach builds upon an analysis of transcriptional expression of gene ensembles ordered according to the normalized amount of change in time (*nrmsf*) and the fold change in expression.

In all of the models analyzed, despite temporally different experimental intervals, a self-organized critical transition (SOC) in whole-genome expression was found to play an essential role in the change in the genome state at both the population and single-cell levels. The results suggest that the full or partial erasure of the initial-state sandpile-type criticality (critical behavior) can be an indicator of the genome-state change (see the summary in Table 3). This is due to the fact that the erasure of criticality destroys the SOC control mechanism of the dynamical change in the entire genomic system (Figure 13A).

**Table 3:**
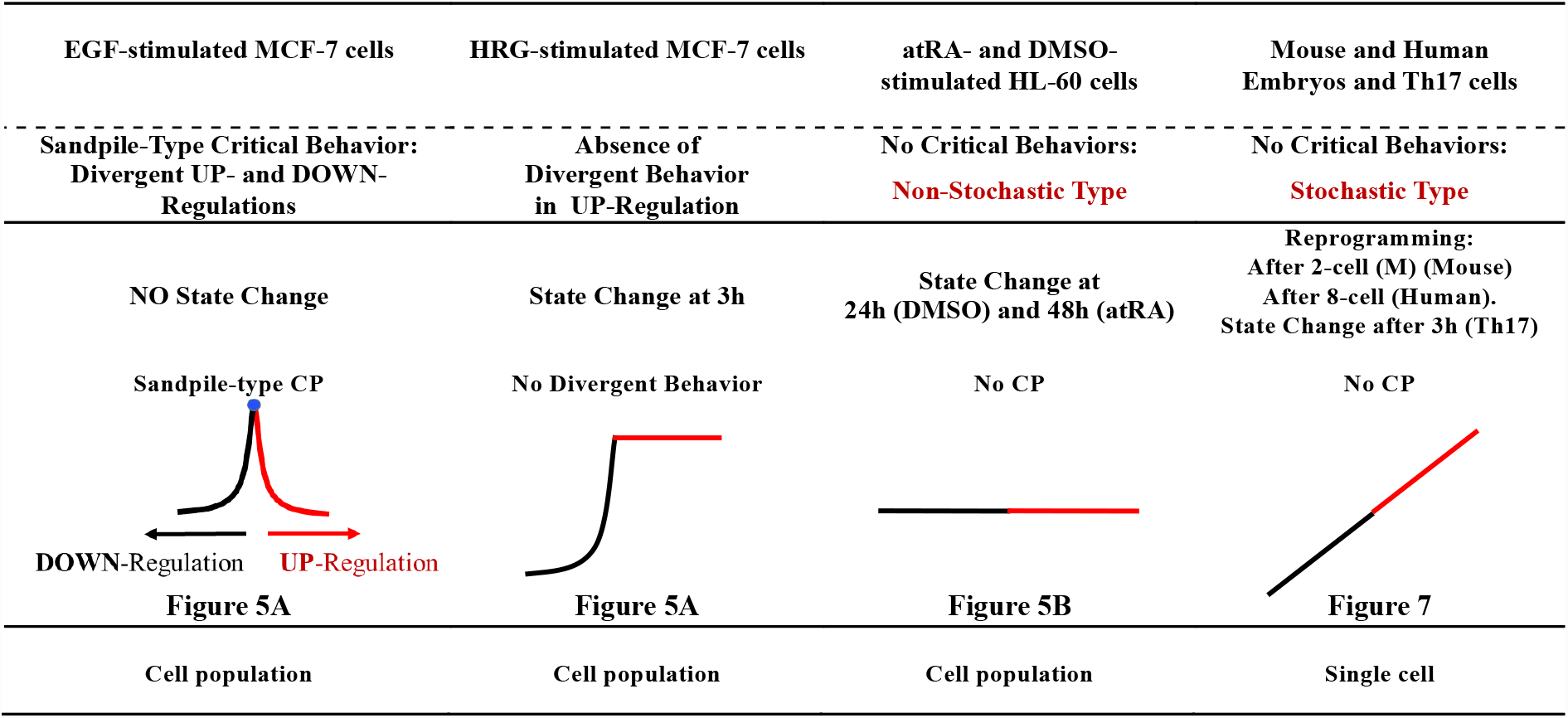
Timing of the Genome-State Change Through Distinct Erasures of Initial-State Critical Behavior: Trends of averaging behavior are shown (represented by solid black and red lines; x-axis: natural log of fold change in expression; y-axis: natural log of expression) (initial-state: state at t= t0 or zygote state for embryos).

Notably, regarding embryo development, the initiation of reprogramming, i.e., whether or not a single cell successfully achieves reprogramming, can be determined by the erasure of the sandpile-type critical point (CP) or criticality stemming from the initial stage of embryogenesis. Thus, the critical gene ensemble of the sandpile-type criticality should exist to affect the entire genome expression in reprogramming. It is important to note that the erasure of criticality is independent of the choice of the initial-state (see more in **section I**), and this independence further confirms the timing of the cell-fate change for both reprogramming in embryonic development and cell differentiation in terminal cells.

The whole-genome expression self-organizes into distinct critical states (expression compartments) through a critical transition. Their coherent dynamics emerge from an ensemble of stochastic expression (coherent-stochastic behaviors) corresponding to the one-dimensional dynamics of the center of mass (CM) of expression, which exhibited the characteristics of an open thermodynamic system and the dynamic perturbation of SOC control for the genome-state change:

i. The average expression flux of critical states (Figure 12A) clearly showed that, for MCF-7 and HL-60 cells, the collective behavior of low-variance gene ensembles acted as a driving force to transmit their potentiality, or energy of coherent transcription fluctuations, to high-variance genes. This can be interpreted in terms of the metaphor of a ‘genome engine’: A sub-critical state, which is an ensemble of low-variance mRNAs, acts as a large piston that moves only slightly. This work propagates through cyclic flux (like a camshaft) to make a super-critical state, an ensemble of high-variance mRNAs, activated as a small piston that moves greatly, while remaining anti-phase to the dynamics of the sub-critical state. A near-critical state, where a critical transition occurs, acts as an ignition switch of the genome engine. The genome engine emerges only when thousands of molecular gene regulations are integrated through SOC control.
ii. Flux dynamics (expression flux dynamics from their averages) revealed that SOC control (i.e., the genome engine) is perturbed locally or globally to determine the genome-state change. In cell differentiation (HRG: MCF-7 cells, and DMSO and atRA: HL-60 cells), the genome-state changes occur at the end of the dissipation of global pulse-like perturbations, which involve the overall expression through the relevant responses of other critical states in addition to the super-critical state. In contrast, in EGF-stimulated MCF-7 cells (cell proliferation and no differentiation), no genome-state change can occur due to the local perturbation: only the super-critical state is affected, while other states show only very weak responses (i.e., they remain near their average expression fluxes). Thus, the global and local perturbations of SOC control differentiate MCF-7 cell fates, whereas global perturbations underlie the state change in HL-60 cells.

These results give us a mechanism of the genome-state change in terminal cell fates: the (partial or full) erasure of an initial-state critical behavior leads to a critical change in the genome state at the end of a dissipative pulse-like global perturbation in self-organization. Regarding early embryo development, further studies will be needed to confirm this mechanism for regulating the cell fate. Note that in single-cell embryonic development, when we consider the number of cell states as different time points, a similar approach to the terminal cell-fate change (**sections III-VI**) can be taken to investigate the mechanism of how genome reprogramming can occur through the breakdown of self-organization.

Notably, an analysis of the literature strikingly confirms that the time intervals of the observed thermodynamic changes revealed by an expression flux approach to transcriptome data are consistent with real biological critical events that determine the change in cell fate. The embryo model is particularly intriguing for our purpose because, in contrast to both HL-60 and MCF-7 cells, it is not based on the average behavior of a huge population, but rather on the behavior of a very few cells at a time.

The above findings lead to some important consequences:

### i Biological Interpretations of the Global and Local Perturbations of SOC Control in MCF-7 Cells

Regarding our present finding of the global perturbation in the SOC control in MCF-7 cell-HRG dynamics, a study of the early gene response [28] established that EGF and HRG induced a transient and sustained dose-dependent phosphorylation of ErbB receptors, respectively, followed by similar transient and sustained activation kinetics of Akt and ERK.

Following ERK and Akt activation from 5-10min, the ligand-oriented biphasic induction of early transcription key proteins of the AP-1 complex (c-FOS, c-MYC, c-JUN, and FRA-1) took place: high for HRG and negligible for ERG. The proteins of the AP-1 complex are non-specific stress-responders supported by the phosphorylation of ERK in a positive feedback loop. In addition, the key reprogramming transcription factor c-MYC (the protein of which peaked at 60min, as confirmed in JE’s laboratory at Latvian Biomedical Research & Study Centre) can amplify the transcription of thousands of active and initiated genes [38] or direct-indirect targets [39] and modify chromatin by recruiting histone acetyltransferase [40]. Further Saeki *et al*. [41] revealed that after HRG the early sustained by ERK activation of AP-1, c-FOS in particular, induces, the sequential activation of late transcription factors EGR4, FOSL-1, FHL2, and DIPA peaking at 3h. In turn, those begin to down-regulate an ERK proliferative pathway by a negative feedback loop. This allowed differentiation to occur (differentiation needs suppression of proliferation).

Thus, the continuity of the biological relay of the HRG-induced early (pre-committing) and late transcription activities leads to a commitment of differentiation from 3h (cell-fate change), which is necessarily coupled to the suppression of proliferation and stops the genome boost by the ERK pathway. Neither a sustained ERK dependent positive feed-back loop, nor the following negative feed-back is achieved in the case of ERG. Subsequently, these cells did not differentiate but continued proliferation.

In the corresponding expression data, we observed, after HRG, a powerful genome engine causing a pulse-like global perturbation (15-20min) as pre-committing and erasure of the initial-state critical transition to induce the genome-state change at 3h, which was not observed after treatment with EGF, in which case only local perturbation (i.e., only vivid activity of the super-critical state) was observed.

It is important to note that the sub-critical state (low-variance expression: a majority of mRNA expressions; Table 1) generates the global perturbation in the HRG response (**section V**). The dynamic control of gene expression in the sub-critical state is expected from the cooperative ensemble behaviors of genomic DNA structural phase transitions (see (v) below) through interaction with environmental small molecules, which has not been considered in previous biological studies. Thus, a true biological picture for MCF-7 cells may be obtained by deciphering the biological functions of genes in the sub-critical state in a coordinated (rather than an individual) manner.

### ii Global Perturbations of SOC Control in HL-60 Cells Committed to Differentiation

The general mechanisms of the commitment to differentiation are not yet well understood. Developmental biologists usually discriminate the two phases into (1) reversible, with the capability of autonomous differentiation; and (2) essentially irreversible [42]. A study in an HL-60-DMSO cell model [43] found that a minimum induction time of 12h was needed for cells to commit to differentiation. In turn, Tsiftsoglou *et al*. [44] found that exposure to differentiation inducers for only 8 to 18h, which is much shorter than the duration of a single generation, is needed to provide commitment for autonomous terminal differentiation.

Consistent with these findings, we revealed that, at 12-18h, global perturbation involving critical states is observed for the responses to both DMSO and atRA. The induction of differentiation for both inducers is different in the sense that, in contrast to DMSO, which induces the development of macrophages/monocytes, treatment with atRA leads to segmented neutrophils [44]. Interestingly, our analysis also shows that the achievement of cell-fate determination at 24h for DMSO and at 48h for atRA (Figure 5B) occurs in these two models in different ways, although they converge at the same final state at 48h.

In particular, we observed early global genome perturbation in the response to atRA (at 0-4h), which was not seen in the DMSO model. This may be due to a difference in Ca^2+^ influx. Calcium release from the ER combined with capacitative calcium influx from the extracellular space leads to markedly increased cytosolic calcium levels and is involved in cell activation [45] to control key cell-fate processes, which include fertilization, early embryogenesis [46], and differentiation [47]. The amplitude and duration of the Ca^2+^ response are decoded by downstream effectors to specify cell fates [48]. Yen *et al*. [49] and others have shown that pre-committed HL-60 cells display early cytosolic Ca^2+^ influx. Moreover, intracellular calcium pump expression is modulated during myeloid differentiation of HL-60 cells in a lineage-specific manner, with higher actual flux in the atRA response [50].

### iii SOC Control in Human and Mouse Related to the Developmental Oocyte-to-Embryo Transition

Fertilized mature oocytes change their state to become developing embryos. This process implies a global restructuring of gene expression. The transition period is dependent on the switch from the use of maternally prepared stable gene transcripts to the initiation of proper embryonic genome transcription activity. It has been firmly established that, in mice, the major embryo genome activation occurs at the two-cell stage (precisely between the mid and late 2-cell states), while in humans this change occurs between the 4- and 8-cell stages [51]. We detected these time intervals, which differ for mouse and human, in the development of SOC control: from sandpile-type critical transitions to stochastic expression distributions (Figures 7A,B). Reprogramming of the genome destroys the SOC control of the initial stage of embryogenesis. Developmental studies by Wennekamp *et al*. in Hiiragi’s group [52] revealed the onset of symmetry breaking between cells in the early embryo and consequently the specification of distinct cell lineages strictly consistent with our model. In addition, what about the physical state of chromatin during these developmental steps?

The erasure of paternal imprinting by DNA 5-methylcytosine de-methylation and hydroxymethylation as part of epigenetic reprogramming occurs in the embryo, which allows significant decompaction of the repressive heterochromatin and an increase in the flexibility of the transcribing part of chromatin [53]. Detailed studies of the DNA methylation landscape in human embryo by Guo and colleagues [30] revealed a large decrease in the level of methylation of gene promoter regions from the zygote to the 2-cell stage, which would erase oocyte imprinting. The strength of this correlation increases gradually until it becomes particularly strong after human embryonic genome activation (full reprogramming) at the 8-cell stage [54]. DNA de-methylation unpacks repressive heterochromatin, which manifests in the dispersal and spatio-temporal reorganization of pericentric heterochromatin as an important step in embryonic genome reprogramming [55]. This synchronization of the methylome to confer maximum physical decompaction and flexibility to chromatin and the reprogramming of transcriptome activity for totipotency support the feasibility of SOC, which was determined here by an independent method.

In addition, transposable elements, which are usually nested and epigenetically silenced in the regions of hypermethylated constitutive heterochromatin, also become temporarily activated during the oocyte-to-embryo transition in early embryogenesis [56]. In turn, in human, a peak in SINE activation coincides with the 8-cell stage [30]. The significance of transient retrotransposon activation in embryogenesis, as has been suggested [53], may be that the thousands of endogenous retro-elements in the mouse genome provide potential scope for large-scale coordinated epigenetic fluctuations (further harnessed by piRNA along with *de novo* DNA methylation). In other words, this should create a necessary critical level of transcriptional noise as a thermodynamic prerequisite for the non-linear genome-expression transition using SOC.

### iv The Extended Concept of Self-Organized Criticality in the Cell-Fate Decision

The recent success with induced pluripotent stem cells (iPSCs) [1] is a remarkable breakthrough not only for possible manipulation of the cell fate through somatic genome reprogramming, but also for understanding the mechanisms of both development and disease.

However, it is still a daunting challenge to elucidate the mechanism of how the transition to a different mode of global gene expression involving the entire genome through a few reprogramming stimuli occurs to achieve the self-control of on/off switching for thousands of functionally unique heterogeneous genes in a small and highly packed cell nucleus.

A fundamental issue is to elucidate the mechanism of the self-organization at the ‘whole genome’ level of gene-expression regulation that is responsible for massive changes in the expression profile through genome reprogramming. To explain this mechanism of massive state changes, self-organized criticality (SOC) has been found to be one of the most important discoveries in statistical mechanics. SOC was proposed as a general theory of complexity to describe self-organization and emergent order in non-equilibrium systems (thermodynamically open systems). However, despite of considerable research over the past several decades, a universal classification has not yet been developed to construct a general mathematical formulation of SOC [20]. Useful background information on SOC is found in [20, 57-63].

Recently, the concept of SOC has been used (and extended) to propose a conceptual model of the cell-fate decision (critical-like self-organization or rapid SOC) through the extension of minimalistic models of cellular behavior [64, 65]. A basic principle for the cell-fate decision-making model (a coarse-grained model, the same as in our approach) is that gene regulatory networks adopt an exploratory process, where diverse cell-fate options are first generated by the priming of various transcriptional programs, and then a cell-fate gene module is selectively amplified as the network system approaches a critical state [66,67]; these review articles also present a useful survey of studies on self-organization in biological systems.

Self-organizing critical dynamics of this type are possible at the edge between order and chaos and are often accompanied by the generation of exotic patterns. Self-organization is considered to occur at the edge of chaos [68-71] through a phase transition from a subcritical domain to a supercritical domain: the stochastic perturbations initially propagate locally (i.e., in a sub-critical state), but due to the particularity of the disturbance, the perturbation can spread over the entire system in an autocatalytic manner (into a super-critical state) and thus, global collective behavior for self-organization develops as the system approaches its critical point.

We checked the above paradigm in different cases of cell-state transitions at both the population and single-cell levels, and interpreted the observed changes in gene expression in terms of critical-like self-organization. A rigorous statistical-mechanical analysis of the time-development of perturbation in self-organization (see **sections IV-VI**) revealed the relevant features, however, these were somewhat different from classical SOC models. We found that:

1) SOC control in overall expression exists at both the population and single-cell levels. In the cell-fate change at the terminal phase (determination of differentiation for the cell population), SOC does not correspond to a phase transition in the overall expression from one critical state to another. Instead, it represents a self-organization of the coexisting critical states through a critical transition.

2) The timing of the genome-state change (i.e., cell-fate change in the genome) is determined through erasure of the initial-state criticality at both the population and single-cell levels. This suggests the existence of specific molecular-physical routes for the erasure of critical dynamics for the cell-fate decision. Furthermore, the cell-fate change (commitment to cell differentiation) in terminal cell fates occurs at the end of dissipative global perturbation in self-organization.

3) The sub-critical state (ensemble of low-variance gene expressions) sustains critical dynamics in SOC control of terminal cell fates. Furthermore, a sub-critical state forms a robust cyclic state-flux with a super-critical state (ensemble of high-variance gene expressions) through the cell nuclear environment. The results show that there is no fine-tuning of an external driving parameter to maintain critical dynamics in SOC control.

It has been pointed out that fine-tuning of a driving parameter for self-organization in classical SOC generates a controversy regarding the real meaning of self-organization (see more in [66]).

However, despite these differences, the occurrence of global and local perturbations in cell-fate decision processes suggests that there may be another layer of a macro-state (genome state) composed of distinct micro-critical states (found by us). After a pulse-like global perturbation occurs in multiple micro-states through the erasure of initial-state criticality, the genome state transitions to be super-critical to guide the cell-fate change (HRG-stimulated MCF-7 cells, and DMSO- and atRA-stimulated HL-60 cells). On the other hand, the genome state of EGF- stimulated MCF-7 cells remains sub-critical (no cell-fate change) because the local perturbation only induces the activation of a micro-critical state. Further studies on this matter are needed to clarify the underlying fundamental mechanism, and the development of a theoretical foundation for the SOC control mechanism as revealed in our findings is needed.

### v Mechanism of the ‘Genome Engine’ in Self-Organization and Genome Computing

In the genome engine, the role of the sub-critical state as a generator leads to a new hypothesis that the genomic compartment, spanning from kbp to Mbp, which produces low-variance gene expression may be the mechanical material basis for the generation of global perturbation, where the coordinated ensemble behavior of genomic DNA structural phase transitions through interactions with environmental molecules plays a dominant role in the expression dynamics [72,73]. In HRG-stimulated MCF-7 cell differentiation, a subset of consecutive genes pertaining to the same critical state (called barcode genes from kbp to Mbp; see Figure 8A in [11]) on chromosomes has been shown to be a suitable material basis for the coordination of phase-transitional behaviors. The critical transition of barcode genes was shown to follow the sandpile model as well as genome avalanche behavior. This indicated that there is a non-trivial similarity through SOC between the coherent-stochastic network of genomic DNA transitions and the on-off nerve-firing in neuronal networks [63]. Thus, a potential function of the genome engine may be asynchronous parallel computing, and coherent-stochastic networks based on the on/off switching of sub-critical barcode genes may act as rewritable self-organized memory in genome computing.

Here we have highlighted how the genome engine, i.e., the global response, is driven by the oscillatory behavior of a sub-critical state, i.e., of genes for which the change in expression is quantitatively minor but strongly coherent. This coherence appears when we consider ensembles of genes with *n* (number of genes) greater than 50 [11]; this behavior is consistent with the particular organization of chromatin into topology-associated domains. The relative flexibility of each domain is probably associated with the changes in expression.

The demonstration of a ‘genetic energy flux dynamics’ across different critical states tells us that chromatin is traversed by coherent waves of condensation/de-condensation, analogous to the allosteric signals in protein molecules. The possibility of controlling such signal transmission through control of the higher-order structure of genomic DNA raises the possibility of very intriguing applications, such as in the much more mature case of allosteric drugs [74,75].

## Methods

### Biological Data Sets

We analyzed mammalian transcriptome experimental data for seven distinct cell fates in different tissues:

i. Cell population: Microarray data of the activation of ErbB receptor ligands in human breast cancer MCF-7 cells by EGF and HRG; Gene Expression Omnibus (GEO) ID: GSE13009 (*N* = 22277 mRNAs; experimental details in [41]), which has 18 time points: *t*_0_ = 0, *t*_1_ = 10, 15, 20, 30, 45, 60, 90min, 2, 3, 4, 6, 8, 12, 24, 36, 48, *t*_*T*=17_ = 72h,
ii. Cell population: Microarray data of the induction of terminal differentiation in human leukemia HL-60 cells by DMSO and atRA; GEO ID: GSE14500 (*N* = 12625 mRNAs; details in [26]), which has 13 time points: *t*_0_ = 0, *t*_1_ = 2, 4, 8, 12, 18, 24, 48, 72, 96, 120, 144, *t*_*T*=12_ = 168h,
iii. Single cell: RNA-Seq data of T helper 17 cell differentiation from mouse naive CD4+ T cells in RPKM values (Reads Per Kilobase Mapped), where Th17 cells are cultured with anti-IL4, anti-IFNγ, IL-6 and TGF-β (details in [76]); GEO ID: GSE40918 (mouse: *N* = 22281 RNAs), which has 9 time points: *t*_0_ = 0, *t*_1_ = 1,3,6,9,12,16,24, *t*_*T*=8_ = 48h,
iv. Single cell: RNA-Seq data of early embryonic development in human and mouse developmental stages in RPKM values; GEO ID: GSE36552 (human: *N* = 20286 RNAs) and GSE45719 (mouse: *N* = 22957 RNAs) (experimental details in [77]) and [78], respectively). We analyzed 7 human and 10 mouse embryonic developmental stages:
Human: oocyte (*m*=3), zygote (*m*=3), 2-cell (*m*=6), 4-cell (*m*=12), 8-cell (*m*=20), morula (*m*=16) and blastocyst (*m*=30),
Mouse: zygote (*m*=4), early 2-cell (*m*=8), middle 2-cell (*m*=12), late 2-cell (*m*=10), 4-cell (*m*=14), 8-cell (*m*=28), morula (*m*=50), early blastocyst (*m*=43), middle blastocyst (*m*=60) and late blastocyst (*m*=30), where *m* is the total number of single cells.

For microarray data, the Robust Multichip Average (RMA) was used to normalize expression data for further background adjustment and to reduce false positives [79-81], whereas for RNASeq data, RNAs that had zero RPKM values over all of the time points were excluded. Random real numbers in the interval [0-1] generated from a uniform distribution were added to all expression values for the natural logarithm. This procedure avoids the divergence of zero values in the logarithm. The robust mean-field behavior through the grouping of expression (see **section II** or Figures 7B-D, 8) was checked by multiplying the random number by a positive constant, *a* (*a*< 10), and thus, we set *a* = 1. Note: The addition of large random noise (*a*>>10) destroys the sandpile CP.

## Emergent Properties of SOC in the Mean-Field Approach

In this report, we examine whether characteristic behavior at a critical point (CP) is present in overall expression to investigate the occurrence of SOC in various cell fates. We briefly summarize below how SOC in overall gene expression was elucidated in our previous studies [10,11].

i. **Global and local genetic responses through mean-field approaches**: Our approach was based on an analysis of transcriptome data by means of the grouping (gene ensembles) of gene expression (mean-field approach) characterized by the amount of change in time to reveal the coexistence of local and global gene regulations in overall gene expression [82,83]. A global expression response emerges in the collective behavior of low- and intermediate-variance gene expression, which, in many expression studies, is cut-off from the whole gene expression by an artificial threshold, whereas a local response represents the genetic activity of high-variance gene expression as elucidated in molecular biology.
ii. **Self-organized criticality as an organizing principle of genome expression**: To understand the fundamental mechanism/principle for the robust coexistence of global and local gene regulation, and further, the role of the global gene expression response, we elucidated a self-organized criticality (SOC) principle of genome expression that could account for global gene regulation [11]. In self-organization, the temporal variance of expression, *nrmsf* (normalized root mean square fluctuation), acts as an order parameter to self-organize whole gene expression into distinct expression domains (distinct expression profiles) defined as *critical states*, where *nrmsf* is defined by dividing *rmsf* by the maximum of overall {*rmsf*_*i*_}:

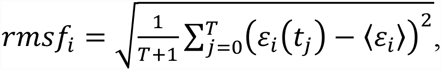

where *rmsf*_*i*_ is the *rmsf* value of the *i*^*th*^ expression (mRNA or RNA), which is expressed as *ε*_*i*_(*t*_*j*_) at *t* = *t*_*j*_ *(j* =0,1,..,T) and 〈*ε*_*i*_〉 is its temporal average (note: we use an overbar for a temporal average when ensemble and temporal averages are needed to distinguish).
iii. **Coherent-stochastic behaviors in critical states**: Coherent expression in each critical state emerges when the number of stochastic expressions is more than 50 [11] (called coherent-stochastic behavior: CSB). The bifurcation - annihilation of the ensemble of coherent gene expression determines the boundary of critical states (Figure 5 in [10]).
iv. (iv) **Biophysical reason for specific groupings**: Coherent dynamics such as coherent oscillation emerge in the change in expression (e.g., fold change) (refer to Figure 7 in [10]; static and dynamic domains correspond to sub- and super-critical states, respectively), but not in expression itself. Furthermore, the change in expression but not expression itself, is amplified in a critical state (Figure 6B in [11]).). This indicates stochastic resonance effect in the change in expression. Thus, regarding self-organization, we investigated averaging behaviors in *nrmsf* (order parameter) and the change in expression.
v. **Characteristic properties of SOC**: Distinctive critical behaviors emerge at a critical point in the averaging of two observables: the fold change in expression and *nrmsf*. Importantly, different critical behaviors (i.e., criticality) that occur at the same CP can originate from different averaging behaviors (MCF-7 cells), which confirms that the mean-fields are not a statistical artifact. We call these *sandpile-type transitional behavior and scaling-divergent behaviors at a critical point* (CP) summarized as follows:
  a. **Sandpile-type transitional behavior** is based on the grouping of expression into *k* groups with an equal number of *n* elements according to the fold change in expression. A sandpile-type transition is the most common in SOC [84]. As *n* increased, the average value of a group (a mean-field) converges, and an ensemble of averages exhibits a sandpile profile. Good convergence in the group is obtained above *n* = 50 (Figure 2 in [11]), which stems from CSB. The top of the sandpile is a critical point (CP), where the CP usually exists at around a zero-fold change (null change in expression), and up- and down-regulated expression is balanced between different time points. This indicates that the critical behavior occurs through a ‘flattened expression energy profile’. This flatness suggests that the strength of the correlation tends to increase with the size of the system ensemble. As noted, a critical point can exist away from a zero-fold change through erasure of the initial-state criticality. A sandpile-type critical behavior shows that, as the distance from the CP (summit of the sandpile) increases, two different regulatory behaviors, which represent up-regulation and down-regulation, respectively, diverge. Furthermore, in the vicinity of the CP according to *nrmsf*, in terms of coherent expression, self-similar bifurcation of overall expression occurs to show transitional behavior (Figures 1, 3A in [11]). Thus, since a critical behavior and a critical transition occur at the CP, we can characterize it as a *sandpile-type transition*.
  b. **b. Scaling-divergent behavior** (**genomic avalanche)** is based on the grouping of expression according to *nrmsf*: a nonlinear correlation trend between the ensemble averages of *nrmsf* and mRNA expression at a fixed time point. Originally, we called this the DEAB (*Dynamic Emergent Averaging Behavior*) of expression [10], which has both linear (scaling) and divergent domains in a log-log plot. In the scaling domain, the quantitative relationship between the ensemble averages of *nrmsf* and mRNA expression, <*nrmsf*> and <*ε*>, is shown in terms of power-law scaling behavior, where higher <*nrmsf*> corresponds to higher <*ε*>:

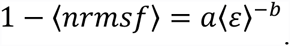 Such scaling is lost at the CP in the MCF-7 cell fates. This shows that order (scaling) and disorder (divergence) are balanced at the CP, which presents genomic avalanches. The scaling-divergent behavior reflects the co-existence of distinct response domains, i.e., critical states. In **Supplementary File S1**, we address the genuineness of the power-law scaling and the existence of collective behavior of gene expression in power-law scaling. Chromosomes exhibit a fractal organization; thus, the power law behavior may reveal a quantitative relation between the aggregation state of chromatin through *nrmsf* and the average expression of an ensemble of genes. The entity of gene expression likely scales with the topology-associated chromatin domains (TADs), such that the degree of *nrmsf* should be related to the physical plasticity of genomic DNA and the high-order chromatin structure.

## Dynamic Flux Analysis for an Open Thermodynamic Genomic System

Sloppiness in SOC control reveals that genome expression is self-organized into a few critical states through a critical transition. The emergent coherent-stochastic behavior (CSB) in a critical state corresponds to the scalar dynamics of its center of mass (CM), *X*(*t*_j_), where X ∈ {Super, Near, Sub}: Super, Near and Sub represent the corresponding critical states. Thus, the dynamics of *X*(*t*_j_) are determined by the change in the one-dimensional effective force acting on the CM. We consider the effective force as a net flux of incoming flux from the past to the present and outgoing flux from the present to the future. Based on this concept of flux, it becomes possible to evaluate the dynamical change in the genetic system in terms of flux among critical states through the environment.

### 1 Self-flux

The effective force is deduced as the decomposition of IN flux from the past (*t*_j-1_) to the current time (*t*_j_) and OUT flux from the current time (*t*_j_) to a future time (*t*_j+1_), so that the effective force as a net self-flux, *f* (*X*(*t*_*j*_)) from its temporal average, can be written as

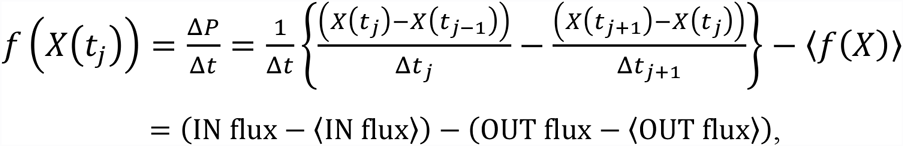

where Δ*P* is the change in momentum with a unit mass (i.e., the impulse: *F*Δ*t* = Δ*P*) for a time difference: Δ*t* = *t*_j+1_ − *t*_j−1_ and Δ*t*_j_ = *t*_j_ − *t*_j−1_; *t*_j_ is the natural log of the *j*^th^ experimental time point (refer to biological data sets); Δ*t*_j_ = *t*_j_ *- t*_j-1_, the CM of a critical state is 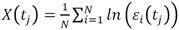 with the natural log of the *i^th^* expression 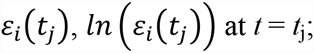 the temporal average of the net self-flux, <*f*(X)> = <INflux> - <OUTflux>, and *N* is the number of expressions in a critical state (Table 1).

As noted, the negative force, *f*(*X*(*t*_j_)), is taken such that a linear term in the nonlinear dynamics of *X*(*t*_j_) corresponds to a force under a harmonic potential energy. The sign of the net self-flux shows the net incoming force to *X*(*t*_j_) (X<=): net IN flux for *f*(*X*(*t*_j_))> 0, and net outgoing force from *X*(*t*_j_) (X=>): net OUT flux for *f*(*X*(*t*_j_))< 0, where the net IN (OUT) flux corresponds to activation (deactivation) flux (force), respectively.

### 2 Interaction flux

The degree of interaction can be evaluated as the exchange of effective force, so that the interaction force can again be decomposed into an IN-coming interaction flux of a critical state, *X*(*t*_j_) from another critical state or the environment, *Y*, from the past (*t*_j-1_) to the current time (*t*_j_), and an OUT-going interaction flux of X(*t*_j_) to *Y* from the current time (*t*_j_) to the future (*t*_j+1_), and the net interaction flux is defined as

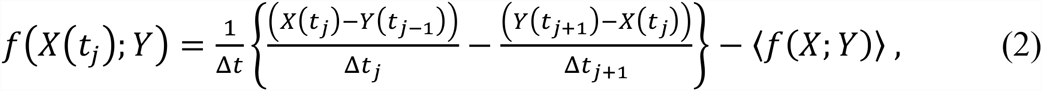

where the interaction flux can be similarly defined by the first and second terms, which represent IN and OUT flux (refer to Equation (1), respectively and *Y* ∈ {Super, Near, Sub, E} with *Y* ≠ *X*: E represents the environment, and 〈*f*(*X*; *Y*)〉 is the temporal average. The sign of the net interaction flux (equation (2)) corresponds to the direction of interaction: net IN interaction flux (Y=>X) for positive and net OUT interaction flux (X=>Y) for negative. The interaction flux between critical states can be defined when the number of expressions in a critical state is normalized.

### 3 The flux network

Due to the law of force, the net self-flux of a critical state, *X*(*t*_j_), is the summation of interaction fluxes (equation (2)) with the other critical states and the environment given by

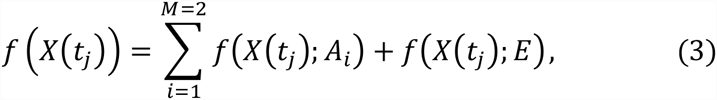

where *A*_*i*_∈ {Super, Near, Sub} with *A*_*i*_≠ *X*, and *M* is the number of internal interactions (*M* = 2). Equation (3) tells us that the sign of the difference between the net self-flux and the overall contribution from internal critical states, 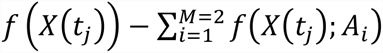 reveals incoming flux (positive) from the environment to a critical state or outgoing flux (negative) from a critical state to the environment; when the difference in all critical states becomes zero, the genome system itself is *closed* thermodynamically (no flux flow from the environment).

### 4 The average flux balance

If we take a temporal average of equation (1) or (3) and equation (2) for the data for both MCF-7 and HL-60 cells, we obtain the average flux balances of a critical state (Table 2), *X*, and its interaction with other states or the environment, respectively:

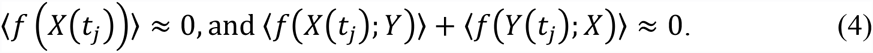

## Contributions

KY and MT initiated the project; MT, AG and KY designed the study; MT analyzed data; KY provided theoretical support; MT and AG wrote the manuscript; JE contributed research and a literature analysis of the biological interpretation of results; and MH performed data treatment to develop a systematic method.

## Acknowledgements

MT sincerely thanks the Institute for Advanced Biosciences, Keio University, Tsuruoka City, and the Yamagata prefectural government, Japan, as well as his family (especially to Taeko and Takako Tsuchiya) for allowing him to complete this research project at Keio University. The authors are thankful to Dr. Vyacheslav Lyashenko for a fruitful discussion.

## Supplementary File S1

Here, we further demonstrate that i) the power-law scaling in the scaling divergent behavior (e.g., Figure 4A) is not a statistical artifact and ii) the dynamics of the collective behavior of gene expression exist through interactions among genes in power-law scaling as follows.

We showed that a scaling region in scaling-divergent behavior (Figure 4A) does actually exist in the sub-critical state (bimodal frequency distribution in MCF-7 cells [Tsuchiya M, *et al*., 2015]).

Figure X1 below (for MCF-7 cells) shows that the bimodality coefficient of random samplings of gene expression converges to that of the sub-critical state, which reveals that, in the power-law scaling (Figure 4A: MCF-7 cells), a self-similar frequency distribution exists among random samplings of gene expression. This result provides additional supporting evidence to further confirm the existence of scaling behavior in the sub-critical state.

**Figure X1:**
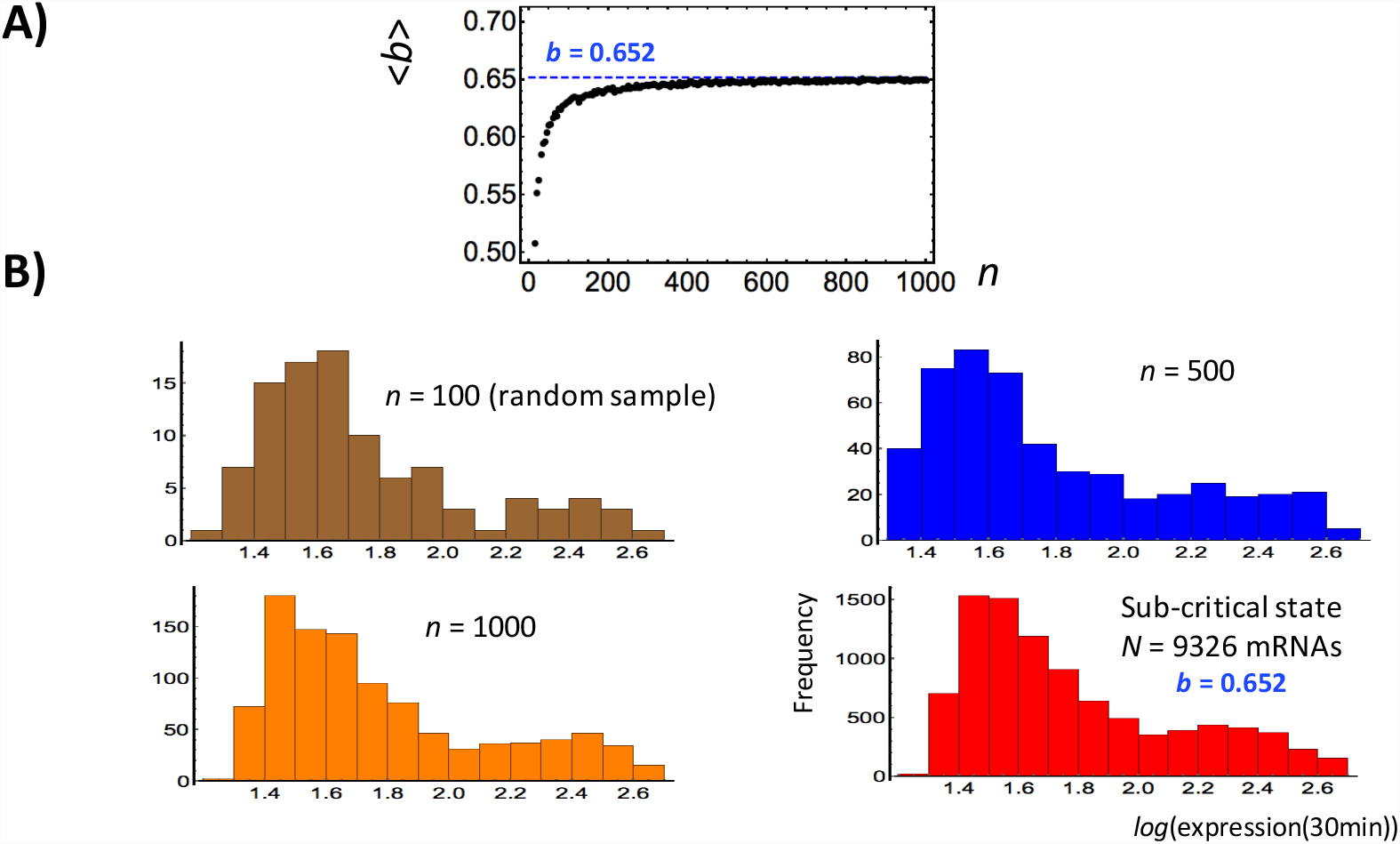
Evidence of scaling behavior in self-similarity of random samplings of gene expression: **A)** The bimodality coefficient of random samplings (*n*: number of randomly selected gene expressions) of gene expression converges to that of the sub-critical state (*b* = 0.652 at *t* =30min, as an example). *b* is Sarle’s bimodality coefficient for a finite sample, and the results show a bimodal or multimodal distribution when *b*> 5/9 (~0.556) An average bimodality coefficient, <*b*>, over 200 repeats is estimated for each random sampling. **B)** The frequency distribution of an ensemble of randomly selected gene expression (*n* = 100: Brown, 500: Blue, 1000 mRNAs: Orange) from the sub-critical state has a self-similar bimodal distribution to that of the sub-critical state (Red).

Next, we demonstrated [Tsuchiya M, *et al.,* 2015] that coherent behavior emerges in the ensemble of stochastic gene expression (coherent-stochastic behavior: CSB). The emergent coherent dynamics follow the dynamics of the center of mass (CM):

(i) The oscillatory coherent dynamics (ABS: Figure X2) have been shown to have a good correlation with the dynamics of its center of mass (see Algebraic correlation of CSB in [Tsuchiya M, *et al*., 2015], and

(ii) The coherent dynamics in random samplings shown in Figure X1 converge to the CM of the sub-critical state (Figure 10A **section IV**). Thus, scaling behavior in the sub-critical state stems from CSB through interactions among genes. In Figure X2, the dynamics of probability density profiles of gene expression in the sub-critical state represent the dynamics of CSB.

Therefore, the power-law scaling is not a statistical artifact and the collective behavior of expression as CSB through interactions among genes is present in power-law scaling.

**Figure X2:**
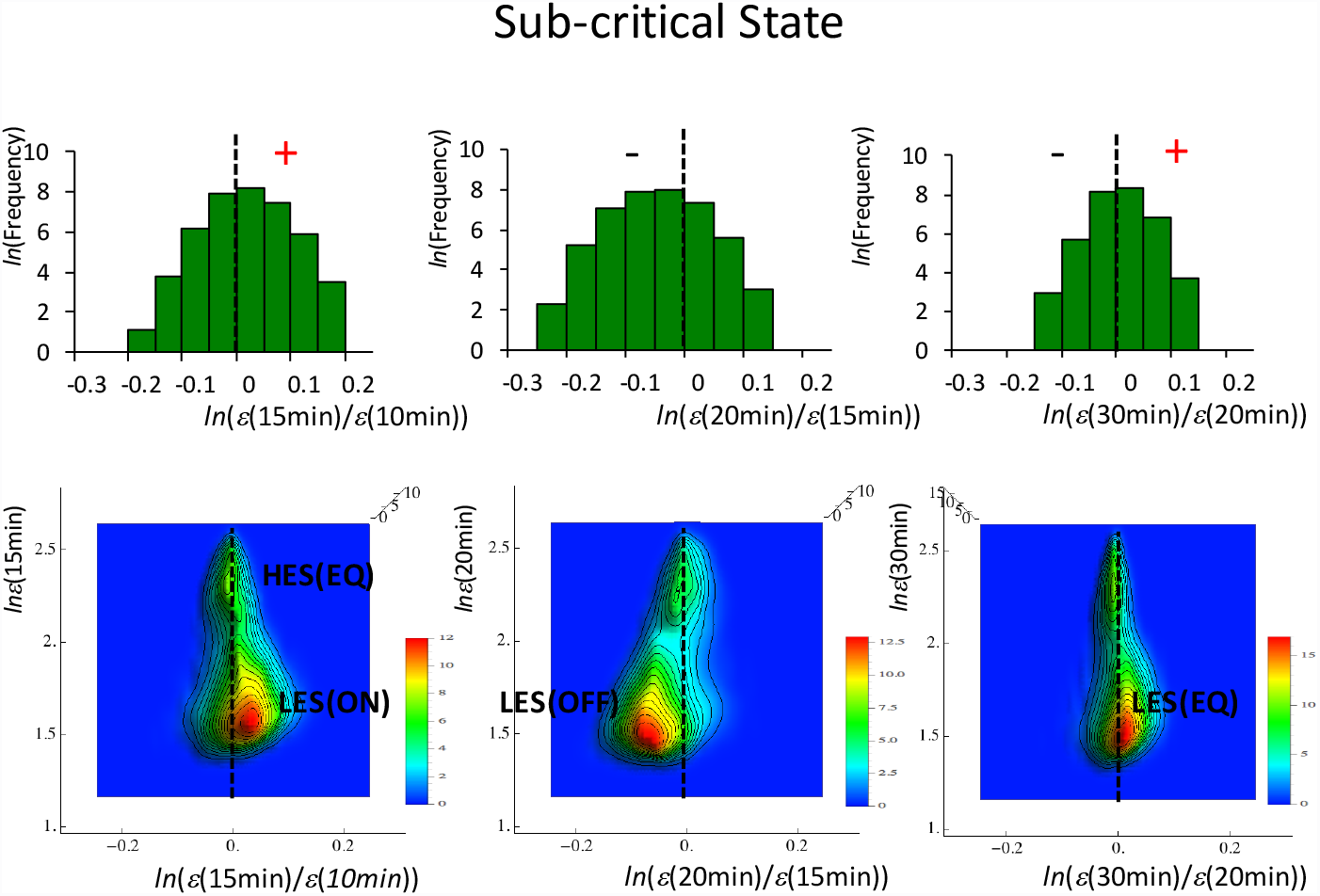
Dynamics of CSB: collective behavior in the sub-critical state (HRG-stimulated MCF-7 cells). Pseudo-3-D probability density profiles for the regulatory space show that two CESs form a pair to develop a pendulum-like oscillatory system, i.e., a low-expression state (LES) that swings around a high-expression state (HES) (Figure taken from Figure 7 in [Tsuchiya M, *et al*., 2014]; ON: up-regulation; OFF: down-regulation; EQ: zero change). This pendulum-like oscillatory system acting as a pair of CESs is defined as an autonomous bistable switch (ABS). Furthermore, the oscillatory dynamics of ABS are shown to be well correlated with the dynamics of the center of mass of ABS (see the detailed analysis of ABS dynamics in [Tsuchiya M, *et al.,* 2015]). *ε*(*t*) represents overall expression at time *t*. Note: Shu and colleagues [Shu G, et al., 2003] demonstrated, by means of density analysis of noisy gene-expression profiles, the robustness of gene-expression clustering.

## A List of Abbreviations

SOC: self-organized criticality;
CP: critical point;
CM: center of mass;
CES: coherent expression state; nrmsf; normalized root mean square fluctuation;
CSB: coherent-stochastic behavior;
ABS: autonomous bistable switch

## Reference

1. Takahashi K, Yamanaka S (2006) Induction of pluripotent stem cells from mouse embryonic and adult fibroblast cultures by defined factors. Cell 131: 861-872.

2. Maherali N, Sridharan R, Xie W, Utikal J, Eminli S, Arnold K, et al. (2007) Directly reprogrammed fibroblasts show global epigenetic remodeling and widespread tissue contribution. Cell Stem Cell 1: 55-70.

3. Efroni S, Duttagupta R, Cheng J, Dehghani H, Hoeppner DJ, Dash C, et al. (2008) Global transcription in pluripotent embryonic stem cells. Cell Stem Cell 2, 437-447.

4. Young RA (2011) Control of the embryonic stem cell state. Cell 144: 940-945.

5. Buganim Y, Dina AF, Rudolf J (2013) Mechanisms and models of somatic cell reprogramming, Nat Rev Genet, 14: 427-439.

6. Raser JM, O’Shea EK (2005) Noise in gene expression: Origins, consequences, and control. Science 309: 2010–2013.

7. Yoshikawa K (2002) Field hypothesis on the self-regulation of gene expression. J Biol Phys 28: 701-712.

8. Yamanaka S (2009) Elite and stochastic models for induced pluripotent stem cell generation. Nature 460: 49-52.

9. Transtrum MK, Machta B, Brown K, Daniels BC, Myers CR, Sethna JP (2015) Perspective: Sloppiness and emergent theories in physics, biology, and beyond. J Chem Phys 143: 010901-010913.

10. Tsuchiya M, Hashimoto M, Takenaka Y, Motoike IN, Yoshikawa K (2014) Global genetic response in a cancer cell: Self-organized coherent expression dynamics. PLOS One 9: e97411.

11. Tsuchiya M, Giuliani A, Hashimoto M., Erenpreisa J, Yoshikawa K (2015) Emergent Self-Organized Criticality in gene expression dynamics: Temporal development of global phase transition revealed in a cancer cell line. PLoS One 11: e0128565.

12. Turalska M, Luković M, West BJ, Grigolini P (2009) Complexity and synchronization. Phys Rev E 80: 021110-021122.

13. Luković M, Vanni F, Svenkeson A, Grigolini P (2014) Transmission of information at criticality, Physica A 416: 430-438.

14. Estrada J, DePace, A, Gunawardena J (2016) Information integration and energy expenditure in gene regulation. Cell 166: 234-244.

15. Li F, Long T, Lu Y, Ouyang Q, Tang C (2004) The yeast cell cycle network is robustly designed. Proc Natl Acad Sci USA 101: 4781-4786.

16. Brandman O, Ferrell Jr JE, Li R, Meyer T (2005) Interlinked fast and slow positive feedback loops drive reliable cell decisions. Science 310: 496-498.

17. Ma’ayan A, Jenkins SL, Neves S, Hasseldine A, Grace E, Dubin-Thaler B, et al. (2005) Formation of regulatory patterns during signal propagation in a mammalian cellular network. Science 309: 1078-1083.

18. Nykter M, Price ND, Aldana M, Ramsey SA, Kauffman SA, Hood LE, et al. (2008) Gene expression dynamics in the macrophage exhibit criticality. Proc Natl Acad Sci USA 105: 1897-1900.

19. Solé RV (2011) Phase Transitions. Princeton University Press.

20. Marković D, Gros C (2014) Power laws and self-organized criticality in theory and nature. Phys Rep 536: 41-74.

21. Lieberman-Aiden E, van Berkum NL, Williams L, Imakaev M, Ragoczy T, Telling A, et al. (2009) Comprehensive mapping of long-range interactions reveals folding principles of the human genome. Science 326: 289-293.

22. Sexton T, Bantignies F, Cavalli G (2009) Genomic interactions: Chromatin loops and gene meeting points in transcriptional regulation. Semin Cell Dev Biol 20: 849-855.

23. Gibcus JH, Dekker J (2013) Connecting the genome: Dynamics and stochasticity in a new hierarchy for chromosome conformation. Mol Cell 49: 773-782.

24. Nagano T, Lubling Y, Stevens TJ, Shoenfelder S, Yaffe E, Dean W, et al. (2013) Single-cell Hi-C reveals cell-to-cell variability in chromosome structure. Nature 502: 59–64.

25. Jost D, Carrivain P, Cavalli G, Vaillant C (2014) Modeling epigenome folding: formation and dynamics of topologically associated chromatin domains. Nucleic Acids Res 42: 9553-9561.

26. Huang S, Eichier G, Bar-Yam Y, Ingber DE (2005) Cell fates as high-dimensional attractor states of a complex gene regulatory network. Phys Rev Lett 94: 128701-128705.

27. Tsuchiya M, Piras V, Giuliani A, Tomita M, Selvarajoo K (2010) Collective dynamics of specific gene ensembles crucial for neutrophil differentiation: the existence of genome vehicles revealed. PLOS One 5: e12116

28. Nagashima T, Shimodaira H, Ide K, Nakakuki T, Tani Y, Takahashi K, et al. (2007) Quantitative transcriptional control of ErbB receptor signaling undergoes graded to biphasic response for cell differentiation. J Biol Chem 282: 4045–4056.

29. Nakakuki T, Birtwistle MR, Saeki Y, Yumoto N, Ide K, Nagashima T, et al. (2010) Ligand-specific c-Fos expression emerges from the spatiotemporal control of ErbB network dynamics. Cell 141: 884–896.

30. Guo H, Zhu P, Yan L, Li R, Hu B, Lian Y, et al. (2014) The DNA methylation landscape of early human embryos. Nature 511: 606-610.

31. MacArthur BD, Lemischka IR (2013) Statistical mechanics of pluripotency. Cell 154: 484–489

32. Schiessel H (2003) Topical Review: The physics of chromatin. J Phys Condens Matter 15: R699-R774.

33. Kornyshev AA (2010) Physics of DNA: unravelling hidden abilities encoded in the structure of ‘the most important molecule’. Phys Chem Chem Phys 12: 12352-12378.

34. Le Treut G, Képès F, Orland H (2016) Phase behavior of DNA in the presence of DNA-binding proteins. Biophys J 110: 51-62.

35. Tremethick DJ (2007) Higher-order structures of chromatin: The elusive 30nm fiber. Cell 128: 651-654.

36. Li G, Reinberg D (2011) Chromatin higher-order structures and gene regulation. Curr Opin in Genet Dev 21: 175-186.

37. Tsuchiya M, Wong ST, Yeo ZX, Colosimo A, Palumbo MC, et al. (2007) Gene expression waves: cell cycle independent collective dynamics in cultured cells. FEBS J 274: 2874–2886.

38. Lin CY, Lovén J, Rahl PB, Paranal RM, Burge CB, Bradner JE, Lee TI, Young RA (2012) Transcriptional amplification in tumor cells with elevated c-Myc. Cell 151: 56-67.

39. Sabò A, Kress TR, Pelizzola M, de Pretis S, Gorski MM, Tesi A, et al. (2014) Selective transcriptional regulation by Myc in cellular growth control and lymphomagenesis. Nature 511: 488-492.

40. Frank SR, Schroeder M, Fernandez P, Taubert S, Amati B (2001) Binding of c-Myc to chromatin mediates mitogen-induced acetylation of histone H4 and gene activation. Genes Dev 15: 2069-2082.

41. Saeki Y, Endo T, Ide K, Nagashima T, Yumoto N, Toyoda T, et al. (2009) Ligand-specific sequential regulation of transcription factors for differentiation of MCF-7 cells. BMC Genomics 10: 545.

42. Gilbert SF (2013) Developmental Biology 10^th^ ed. Sinauer Associates, US.

43. Tarella C, Ferrero D, Gallo E, Pagliardi GL, Ruscetti FW (1982) Induction of differentiation of HL-60 cells by dimethyl sulfoxide: evidence for a stochastic model not linked to the cell division cycle. Cancer Res 42: 445-449.

44. Tsiftsoglou AS, Wong W, Hyman R, Minden M, Robinson SH (1985) Analysis of commitment of human leukemia HL-60 cells to terminal granulocytic maturation. Cancer Res 45: 2334-2339.

45. Papp B, Brouland JP, Arbabian A, Gélébart P, Kovács T, Bobe R, et al. (2012) Endoplasmic reticulum calcium pumps and cancer cell differentiation. Biomolecules 2: 165-186.

46. Whitaker M (2006) Calcium at fertilization and in early development. Physiol Rev 86: 25-88.

47. Schaefer A, Magócsi M, Stöcker U, Kósa F, Marquardt H (1994) Early transient suppression of c-myb mRNA levels and induction of differentiation in Friend erythroleukemia cells by the [Ca2+]i-increasing agents cyclopiazonic acid and thapsigargin. J Biol Chem 269: 8786-8791.

48. Dolmetsch RE, Lewis RS, Goodnow CC, Healy JI (1997) Differential activation of transcription factors induced by Ca2+ response amplitude and duration. Nature 386: 855-858.

49. Yen A, Forbes M, DeGala G, Fishbaugh J (1987) Control of HL-60 cell differentiation lineage specificity, a late event occurring after precommitment. Cancer Res 47: 129-134.

50. Launay S, Giannì M, Kovàcs T, Bredoux R, Bruel A, Gélébart P, et al. (1999) Lineage-specific modulation of calcium pump expression during myeloid differentiation. Blood 93: 4395-4405.

51. Telford NA, Watson AJ, Schultz GA (1990) Transition from maternal to embryonic control in early mammalian development: a comparison of several species. Mol Reprod Dev 26: 90-100.

52. Wennekamp S, Mesecke S, Nédélec F, Hiiragi T (2013) A self-organization framework for symmetry breaking in the mammalian embryo. Nat Rev Mol Cell Biol. 14(7): 452-459.

53. Choy JS, Wei S, Lee JY, Tan S, Chu S, Lee TH (2010) DNA methylation increases nucleosome compaction and rigidity. J Am Chem Soc 132: 1782-1783.

54. Niakan KK, Han J, Pedersen RA, Simon C, Pera RA (2012) Human pre-implantation embryo development. Development 139: 829-841.

55. Yang CX, Liu Z, Fleurot R, Adenot P, Duranthon V, Vignon X, Zhou Q, Renard JP, Beaujean N (2103) Heterochromatin reprogramming in rabbit embryos after fertilization, intra-, and inter-species SCNT correlates with preimplantation development. Reproduction. 145: 149-159.

56. Peaston AE, Knowles BB, Hutchison KW (2007) Genome plasticity in the mouse oocyte and early embryo. Biochem Soc Trans 35: 618-622.

57. Clar S, Drossel B, Schwabl F (1996) Forest fires and other examples of self-organized criticality. J Phys Condens Matter 8: 6803-6824.

58. Jensen HJ (1998) Self-Organized Criticality. Cambridge Univ Press, UK.

59. Kinouchi O, Prado CPC (1999) Robustness of scale invariance in models with self-organized criticality. Phys Rev E 59: 4964-4969.

60. Turcotte DL (2001) Self-organized criticality: does it have anything to do with criticality and is it useful? Nonlinear Process Geophys 8: 193-196.

61. Halley JD, Burd M (2004) Nonequilibrium dynamics of social groups: insights from foraging argentine ants. Insect Soc 51: 226-231.

62. Halley JD, Winkler DA (2008) Critical-like self-organization and natural selection: two facets of a single evolutionary process? BioSystems 92: 148-158.

63. Plenz D, Niebur E (2014) Criticality in Neural Systems. Wiley-VCH, Germany.

64. Huang S, Guo YP, May G, Enver T (2007) Bifurcation dynamics in lineage-commitment in bipotent progenitor cells. Dev Biol 305: 695-713.

65. Cinquin O, Demongeot J (2005) High-dimensional switches and the modelling of cellular differentiation. J Theor Biol 233: 391-411.

66. Halley JD, Burden FR, Winkler DA (2009) Summary of stem cell decision making and critical-like exploratory networks. Stem Cell Res 2: 165-177.

67. Halley JD, Smith-Miles K, Winkler DA, Kalkan T, Huang S, Smith A (2012) Self-organizing circuitry and emergent computation in mouse embryonic stem cells. Stem Cell Res. 8: 324-33.

68. Langton CG (1990) Computation at the edge of chaos - phase transitions and emergent computation. Physica D 42: 12–37.

69. Waldrop MM (1992) Complexity: The Emerging Science at the Edge of Chaos. Simon and Schuster, New York.

70. Bonabeau E (1997) Flexibility at the edge of chaos: a clear example from foraging in ants. Acta Biotheor 45: 29-50.

71. de Oliveira PM (2001) Why do evolutionary systems stick to the edge of chaos? Theory Biosci 120: 1-19.

72. Takenaka Y, Nagahara H, Kitahata H, Yoshikawa K (2008) Large-scale on-off switching of genetic activity mediated by the folding-unfolding transition in a giant DNA molecule: An hypothesis. Phys Rev E 77: 031905.

73. Nagahara H, Yoshikawa K (2010) Large system in a small cell: A hypothetical pathway from a microscopic stochastic process towards robust genetic regulation. Chem Phys Lett 494: 88–94.

74. DeDecker BS (2000) Allosteric drugs: thinking outside the active-site box. Chem Biol 7: R103-R107.

75. Dror RO, Green HF, Valant C, Borhani DW, Valcourt JR, Pan AC, et al. (2013) Structural basis for modulation of a G-protein-coupled receptor by allosteric drugs. Nature 503: 295-299.

76. Ciofani M, Madar A, Galan C, Sellars M, Mace K, Pauli F, et al. (2012) A validated regulatory network for Th17 cell specification. Cell 151: 289-303.

77. Yan L, Yang M, Guo H, Yang L, Wu J, Li R, et al. (2013) Single-cell RNA-Seq profiling of human preimplantation embryos and embryonic stem cells. Nat Struct Mol Biol 20:1131-1139.

78. Deng Q, Ramsköld D, Reinius B, Sandberg R (2014) Single-cell RNA-seq reveals dynamic, random monoallelic gene expression in mammalian cells. Science 343:193–196.

79. Bolstad BM, Irizarry RA, Astrand M, Speed TP (2003) A comparison of normalization methods for high density oligonucleotide array data based on variance and bias. Bioinformatics 19: 185–193.

80. Irizarry RA, Hobbs B, Collin F, Beazer-Barclay YD, Antonellis KJ, Scherf U, et al. (2003) Exploration, normalization, and summaries of high density oligonucleotide array probe level data. Biostatistics 4: 249–264.

81. McClintick JN, Edenberg HJ (2006) Effects of filtering by present call on analysis of microarray experiments. BMC Bioinformatics 7: 49.

82. Tsuchiya M, Selvarajoo K, Piras V, Tomita M, Giuliani A (2009) Local and global responses in complex gene regulation networks. Physica A 388:1738–1746.

83. Tsuchiya M, Piras V, Choi S, Akira S, Tomita M, Giuliani A, et al. (2009) Emergent genome-wide control in wildtype and genetically mutated lipopolysaccharide-stimulated macrophages. PLOS One 4: e4905.

84. Bak P, Tang C, Wiesenfeld K (1987) Self-organized criticality: An explanation of 1/f noise. Phys Rev Lett 59(4): 381–384.

## References

1. Shu G, Zeng B, Chen YP, Smith OH (2003) Performance assessment of kernel density clustering for gene expression profile data. Comp Funct Genomics 4: 287–299.

2. Tsuchiya M,Hashimoto M, Takenaka Y, Motoike IN, Yoshikawa K (2014) Global genetic response in a cancer cell: Self-organized coherent expression dynamics. PLOS One 9: e97411.

3. Tsuchiya M, Giuliani A, Hashimoto M, Erenpreisa J, Yoshikawa K (2015) Emergent Self-Organized Criticality in gene expression dynamics: Temporal development of global phase transition revealed in a cancer cell line. PLoS One 11,

